# An ultrasensitive and modular platform to detect Siglec ligands and control immune cell function

**DOI:** 10.1101/2025.06.10.658684

**Authors:** Zeinab Jame-Chenarboo, Edward N. Schmidt, Madeline Crichton, Kei Takahashi-Yamashiro, Guilherme M. Lima, Liany Luna-Dulcey, Jaesoo Jung, Sabine Ivison, Chris D. St. Laurent, Jhon R. Enterina, Sung-Yao Lin, Susmita Sarkar, Reni John, Som G. Nanjappa, Stacy A. Malaker, Megan K. Levings, Jamey D. Marth, Ratmir Derda, Matthew S. Macauley

## Abstract

Siglecs are immunomodulatory receptors that regulate immune cell function. A fundamental challenge in studying Siglec-ligand interactions is the low affinity of Siglecs for their ligands. Inspired by how nature uses multivalency, we developed Siglec-liposomes as a highly multivalent and versatile platform for detecting Siglec glycan ligands in which recombinant Siglecs were conjugated to liposomes using the SpyCatcher-SpyTag system. Siglec-liposomes offer tunable multivalency and a modular assembly, enabling presentation of different Siglecs on the same liposome. Using Siglec-liposomes, we profiled Siglec ligands on human leukocytes, revealing new insights into Siglec ligands. Moreover, Siglec-liposomes are *in vivo* compatible, where we demonstrated that Siglec-7-liposomes bind to the brain vasculature in a mucin-dependent manner. Given the abundance of Siglec ligands on T cells, we investigated whether Siglec-liposomes modulate T cell function and find that Siglec-7-liposomes increase T cell proliferation in a ST3Gal1-dependent and CD43-independent manner. Taken together, Siglec-liposomes are a versatile and sensitive tool for detecting Siglec ligands and immunomodulation.

## Introduction

Sialic acid-binding immunoglobulin-type lectins (Siglecs) are a family of transmembrane receptors that regulate immune cell function through interactions with their glycan ligands (*1*). Siglec-ligand interactions play many important roles in human health and disease such as cancer (*2*). These interactions can take place with ligands on the same cell (*cis*) or another cell (*trans*). It is notable that interactions of Siglecs with their *trans* sialoglycan ligands occurs with high multivalency, which is essential to compensate for their inherent low monovalent affinity (*3*). The low affinity of Siglecs for their ligands makes it challenging to study Siglec-ligand interactions outside the context of a highly multivalent cell surface. Tools to leverage avidity of Siglecs are needed to study the biological roles of Siglecs outside the context of a lipid bilayer. Moreover, as Siglec family members are expressed together on immune cells (*4*), a modular platform to multiplex numerous Siglecs would more faithfully mimic how Siglecs function together in nature.

For over 30 years, Siglec-Fc chimeras have been the tool of choice to study Siglecs outside the context of a lipid bilayer, which have facilitated many important discoveries (*5–9*). Siglec-Fc proteins have also been extensively used for probing ligands on cancer cells, with many studies documenting the upregulation of Siglec ligands on a wide variety of cancers (*10*). These findings have motivated innovated approaches to block Siglec-ligand interactions to enhance anti-tumor immunity (*2, 3, 11*). Siglec-Fc proteins are likewise useful in drilling deeper into the precise biochemical nature of Siglec ligands; recently, Siglec-Fc proteins have been instrumental in the discovery that carbohydrate sulfation enhances the binding of Siglec-3, -5, -7, -8, and -15 to ligands presented on glycan microarrays (*12, 13*) and cells (*14–16*).

While Siglec-Fc proteins are valuable tools, they have limitations. For example, when used to stain human primary cells and tissues, Siglec-Fcs can produce high background staining, which is due to recognition of IgG adsorbed to the cells and tissues by ahuman IgG secondary antibody that is required to pre-complex the Siglec-Fc. To address the drawbacks associated with using a secondary antibody, we previously introduced Strep-Tactin to detect a Strep-Tag II (*5*), but this has limits on multivalency, therefore, increasing multivalency would have the potential to create much more sensitive detection probes.

Previously, liposomes have been used to present Siglec ligands, which have been successful in probing Siglec-ligand interactions and modulating immune cell function (*3, 17*). For example, liposomes have recently been used to dissect Siglec-glycolipid interactions (*7, 18*), and regulating the proximity of Siglecs to other immune cell receptors (*19, 20*). Importantly, liposomes are *in vivo* compatible (*21*), enabling liposomes displaying Siglec ligands to be used in many applications for targeting Siglecs and modulating immunity (*22, 23*). We hypothesized that liposomes could also be used as a useful multivalent presentation platform for displaying Siglecs. Testing this hypothesis requires robust bio-orthogonal chemistry to link the Siglec to liposome, ideally in a site-specific manner. Recently, we demonstrated that SpyCatcher-SpyTag (*24*) bioconjugation can successfully immobilize Siglec-7 on DNA-barcoded bacteriophages, resulting in constructs that can detect Siglec-7 ligands on cells (*8*). Based on a proven biocompatibility of liposomes *in vivo*, we reasoned that they could offer a more robust and sensitive platform for detecting Siglec-ligand interactions, particularly for *in vivo* studies.

Here, we leveraged SpyCatcher-SpyTag bioconjugation (*24*) to present Siglecs in a multivalent and modular manner from liposomes and demonstrate that the SpyCatcher-SpyTag system creates a tunable multivalent scaffold that can profile Siglec ligands on different cells and tissues in a sensitive and selective manner. Profiling Siglec ligands on regulatory T cells (Tregs) revealed the higher levels of multiple Siglec ligands on Tregs compared to conventional T cells (Tconv). Moreover, Siglec-liposomes can be multiplexed to simultaneously detect different Siglec ligands. Importantly, Siglec-liposomes can detect Siglec ligands *in vivo*, where we found that brain vasculature has abundant levels of mucin-dependent Siglec-7 ligands. Finally, we discovered an immunomodulatory role for Siglec-7, wherein Siglec-7-liposomes stimulated T cell proliferation in a ST3Gal1-dependent and CD43-independent manner.

## Results

### Design, expression, and purification of Siglec-SpyTag proteins

Aiming to use the SpyCatcher-SpyTag system (*24*) for efficient bioconjugation of Siglecs to a multivalent platform, we started from our first generation Siglec-Fc construct (**Fig. 1A**) and incorporated a SpyTag (ST) at two different locations (**Fig. 1B**). In Version 1 (V1), the ST was placed between the Siglec and the Fc (Siglec-ST-Fc) and in Version 2 (V2), the ST was placed at the C-terminus of the Fc (Siglec-Fc-ST). To test these constructs in liposomes, we initially focused on Siglec-7 because it expresses well (*5*), and shows consistent and reliable binding to cells in a mucin-type *O*-glycan-dependent manner (*14–16, 25*). Accordingly, V1 Siglec-7-ST-Fc and V2 Siglec-7-Fc-ST were expressed from CHO cells and doubly purified using the His_6_ and Strep-Tag II affinity tags (**Fig. S1A**). Moreover, TEV protease could be used to turn V1 into monomeric Siglec-ST (**Fig. S1B**).

**Fig. 1.**
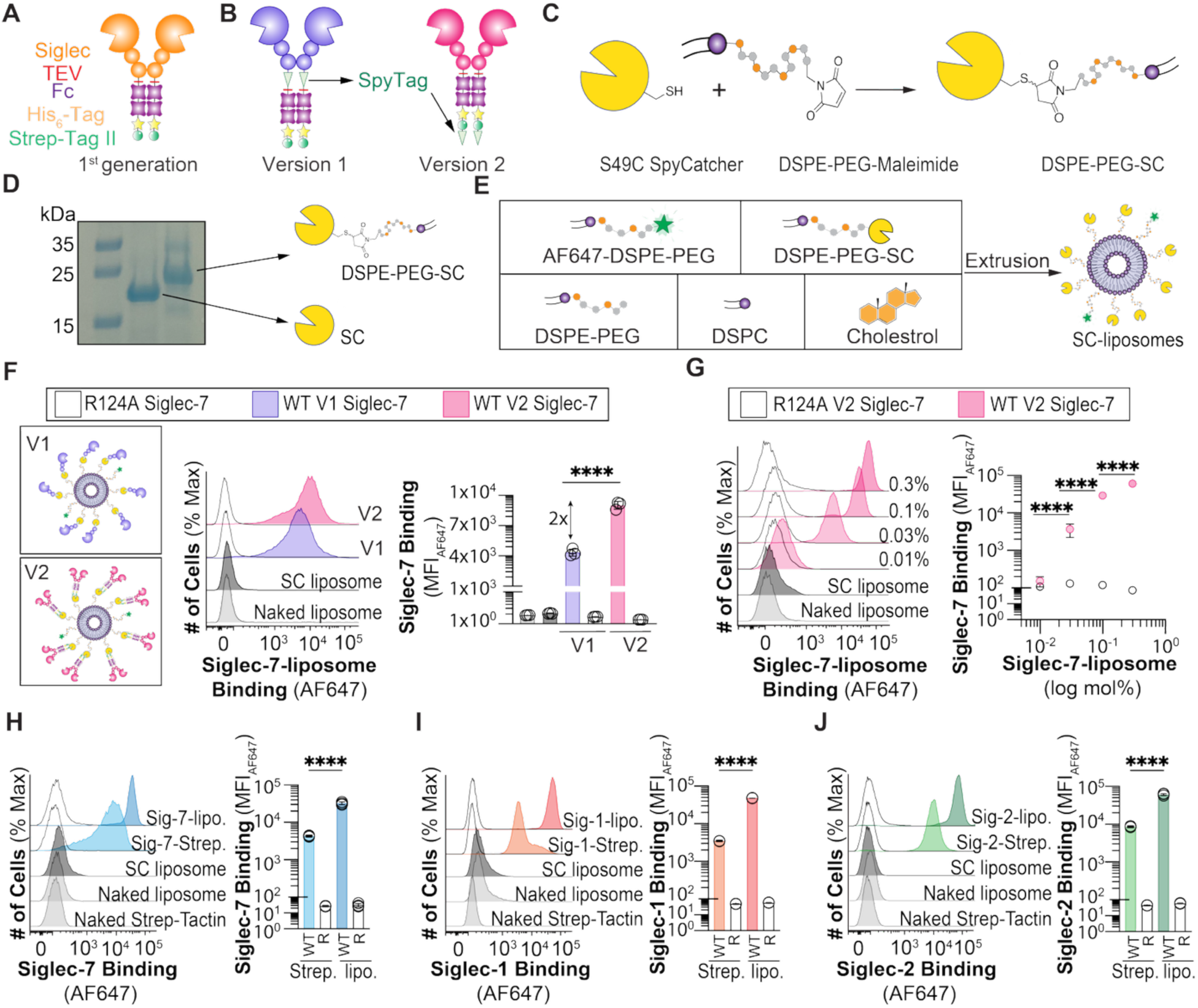
Development and validation of Siglec-liposomes. (**A**) First generation Siglec-Fc. (**B**) V1 and V2 constructs of Siglec-Fc with a genetically-encoded SpyTag. (**C**) Schematic of site-specific conjugation of SC to DSPE-PEG-Maleimide. (**D**) SDS-PAGE of SC and DSPE-PEG-SC. (**E**) Schematic of making SC-liposomes. (**F**) Binding of V1 and V2 WT Siglec-7- and R124A Siglec-7-liposomes to U937 cells. Data is represented as flow cytometry histograms and median fluorescent intensity (MFI) of Siglec-7 binding. (**G**) Binding of V2 of WT Siglec-7 and R124A Siglec-7 in different densities on liposomes to U937 cells. Data is represented as flow cytometry histograms and MFI of Siglec-7 binding. Binding of (**H**) WT and R124A V2 Siglec-7, (**I**) WT and R116A V2 Siglec-1, and (**J**) WT and R120A V2 Siglec-2 pre-complexed with Strep-Tactin (Strep.) and conjugated to liposomes (lipo.) to U937 cells. Data is presented as flow cytometry histograms and MFI of Siglec binding. The *p* value for three technical replicates was calculated using an unpaired one-way ANOVA. \*\*\*\**p*<0.0001.

### Preparation of liposomes displaying SpyCatcher

Aiming to develop multivalent Siglec-liposomes, a S49C SpyCatcher (SC) mutant (*8*) was used for site-specific conjugation to DSPE-PEG-Maleimide (**Fig. 1C**), which was shown to be successful by SDS-PAGE (**Fig. 1D**). To create multivalent SC-liposomes, we used the Doxil formulation (*26*) and substituted a portion of the 5% DSPE-PEG with DSPE-PEG-SC to tune the multivalency of SC (**Fig. 1E**). In addition, 0.1% AF647-DSPE-PEG was added to the liposomal formulation for detection by flow cytometry. SC- liposomes were conjugated with either V1 monomers or V2 dimers of Siglec-7. Conjugation of V2 Siglec-Fc-ST to SC protein and SC-liposomes was verified by SDS-PAGE (**Fig. S2A,B**). We also expressed and conjugated R124A Siglec-7, which cannot interact with its sialic acid-containing glycan ligands (*5, 14*).

To test the SC-liposomes conjugated with V1 Siglec-7 (Siglec-7-ST) or V2 Siglec-7 (Siglec-7-Fc- ST), we analyzed their binding to monocytic U937 cells by flow cytometry (**Fig. 1F**). Both WT versions of Siglec-7 displayed on the liposomes showed robust binding to U937 cells, with the V2 dimeric Siglec-7- liposomes showing significantly higher binding (2-fold) than the V1 monomeric version. This observation is consistent with our previously reported enhanced ligand binding properties of multivalent V2 Siglec-7 on bacteriophage when compared to multivalent display of V1 Siglec-7 (*8*). This enhancement may be the effect of local clustering of glycan binding (V-set) domains, resulting from the dimeric nature of V2 Siglec-7. In in either case, R124A Siglec-7 showed no binding to cells. To assess how the density of Siglec-7 on the liposomes impacts binding, the amount of V2 Siglec-7 was titrated in two different ways: by varying the amount of SC displayed on the liposomes and saturating it with V2 Siglec-7 (**Fig. S3**) or keeping the amount of SC displayed on the liposome constant and varying the amount of V2 Siglec-7 (**Fig. 1G**). For both titrations, a sharp increase of 20-40-fold in binding was observed when V2 Siglec-7 was increased from 0.01 to 0.03 mol%, corresponding to approximately 8 – 80 display copies of Siglec-Fc per liposome. Above 0.1 mol% and up to 0.33 mol% Siglec-Fc, which corresponds to approximately 200 copies of Siglec-Fcs, there was a modest increase (2-fold) in binding of V2 Siglec-7 to U937 cells. Compared to Siglec-7- Fc pre-complexed with Strep-Tactin (*5*), Siglec-7-liposomes demonstrated an 8-fold increase in detecting Siglec-7 ligands on the cell surface (**Fig. 1H**). Titrating the amount of AF647 on the Strep-Tactin and liposomes consistently demonstrated the superior detection sensitivity of Siglec-7-liposomes, ruling out differences in the amount of fluorophore (**Fig. S4**). We also confirmed that V2 Siglec-1- and V2 Siglec-2- liposomes showed enhanced ligand detection on U937 cells compared to using Strep-Tactin as a pre-complexing agent, confirming the compatibility of liposomes with other Siglecs (**Fig. 1I,J**).

### Enhancing sensitivity of the platform using Siglecs expressed from CMAS^-/-^ CHO cells

As Siglecs are naturally masked by *cis* ligands on cells (*7*), we hypothesized that *cis* masking may also be in effect for Siglec-liposomes and that eliminating the sialic acid on the V2 Siglec-Fc-ST proteins would enhance the ability of Siglec-liposomes to detect cellular ligands. To test this hypothesis, V2 Siglec- 7 was expressed from genetically-engineered CMP-sialic acid synthetase deficient (CMAS^-/-^) CHO cells. To compare the binding of V2 Siglec-7 expressed in CHO CMAS^-/-^ with the V2 Siglec-7 expressed in CMAS^+/+^ CHO cells, we conjugated both versions to our optimized SC-liposome formulation (0.3 mol% SC-liposome) and tested their binding to U937 cells (**Fig. 2A**). V2 Siglec-7 expressed in CHO CMAS^-/-^ showed higher binding to U937 cells compared to V2 Siglec-7 expressed in CMAS^+/+^ CHO cells.

**Fig. 2.**
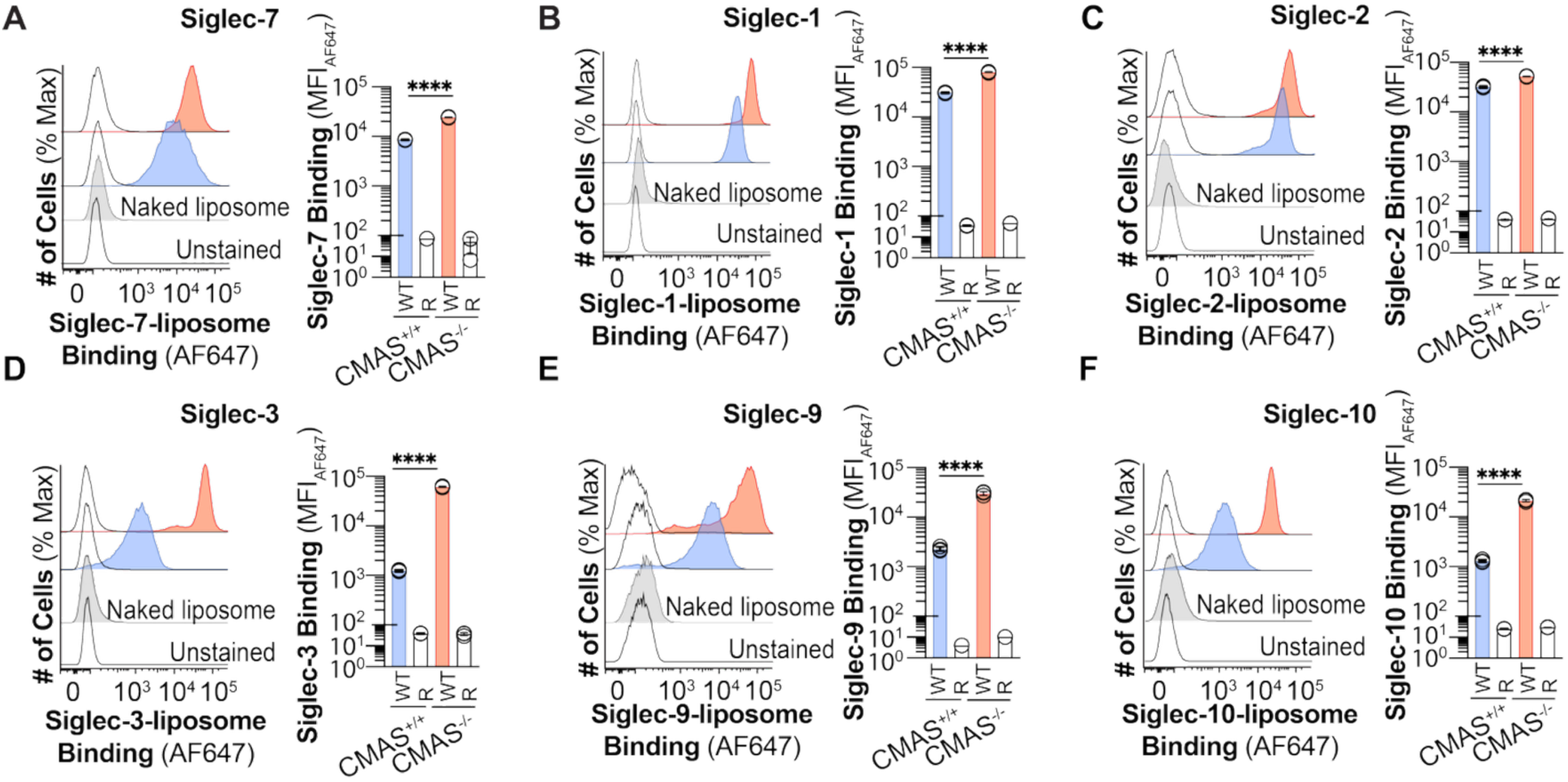
Siglecs expressed in sialic acid deficient CHO cells show improved binding toward their ligands. Binding of WT and arginine mutated of (**A**) Siglec-7-, (**B**) Siglec-1-, (**C**) Siglec-2-, (**D**) Siglec-3-, (**E**) Siglec-9-, and (**F**) Siglec-10-liposomes expressed in CMAS^+/+^ and CMAS^-/-^ CHO cells to U937 cells. Data is presented as flow cytometry histograms and MFI of Siglec binding. The *p* value for three technical replicates was calculated using an un-paired one-way ANOVA. \*\*\*\**p*<0.0001.

Accordingly, we expressed the entire human Siglec family as their V2 Siglec-Fc-ST proteins, along with an essential arginine mutants, from CMAS^-/-^ CHO cells (**Fig. S5**). These proteins were incorporated into SC-liposomes and tested for binding to cells by flow cytometry, which demonstrated that the sialic acid-deficient Siglec proteins showed enhanced binding compared to their counterparts with sialic acid, except Siglec-6 and -11 that did not bind U937 cells (**Fig. 2B-F** and **S6**). None of the arginine mutated versions of Siglecs showed significant binding to cells. To compare the sensitivity of Siglecs expressed in CMAS^-/-^ CHO cells in SC-liposome with Strep-Tactin, we pre-complexed all Siglecs expressed in CMAS^-/-^ CHO cells with Strep-Tactin and conjugated them to SC-liposome and tested their binding to cells. All Siglecs showed a better binding to cells in SC-liposomes compared to the Strep-Tactin, while no binding was observed for the arginine mutated constructs (**Fig. S7**). As an added benefit, sialic acid-deficient Siglec-Fc proteins expressed at a 2-4 times higher yield (**Fig. S8**).

### Using SC-liposomes to profile Siglec ligands on primary cells

Siglec-Fc proteins pre-complexed with secondary αhuman IgG show high background binding to primary immune cells due to the high concentration of IgG (6-17 g/L) present in blood (**Fig. S9**) (*27*). Siglec-liposomes avoid the need for using αhuman IgG as secondary. Therefore, we used Siglec-liposomes to probe Siglec ligands on immune cells from human peripheral blood. Flow cytometry was used to distinguish B cells (CD19^+^), CD4^+^ T cells (CD3^+^, CD4^+^), CD8^+^ T cells (CD3^+^, CD8^+^), mature natural killer (NK) cells (CD56^+^), neutrophils (CD15^+^, CD16^+^), and monocytes (CD14^+^) isolated from peripheral blood mononuclear cells (PBMCs), and initially compared the binding of Siglec-2 pre-complexed with Strep-Tactin with Siglec-2-liposome (**Fig. S10**). Siglec-2-liposomes showed a higher sensitivity for binding to all immune cells compared to Siglec-2 pre-complexed with Strep-Tactin, while no binding was observed for R120A Siglec-2 in either format.

Accordingly, Siglec ligands were analyzed on immune cells isolated from fresh peripheral blood of four healthy individuals (**Fig. 3A** and **S11-16**). Siglec-liposomes showed diverse and broad binding to immune cells, with several notable observations: Siglec-1, -2, -3, -4, -7, and -10 showed significant binding to all different immune cells, with the strongest binding of Siglec-1 to monocytes, Siglec-2 to B cells, Siglec-3 and Siglec-8 to monocytes and CD8^+^ T cells, Siglec-7 to T cells and NK cells, Siglec-9 to NK cells, and Siglec-10 to B cells (**Fig. 3B**). No binding was observed for arginine mutated versions of Siglecs, except for Siglec-6, where WT and R122A Siglec-6 bound at almost the same level to the B cells and CD8^+^ T cells. This latter finding is consistent with our prior observation that the Arg122 is dispensable for recognizing sialylated ligands (*7*).

**Fig. 3.**
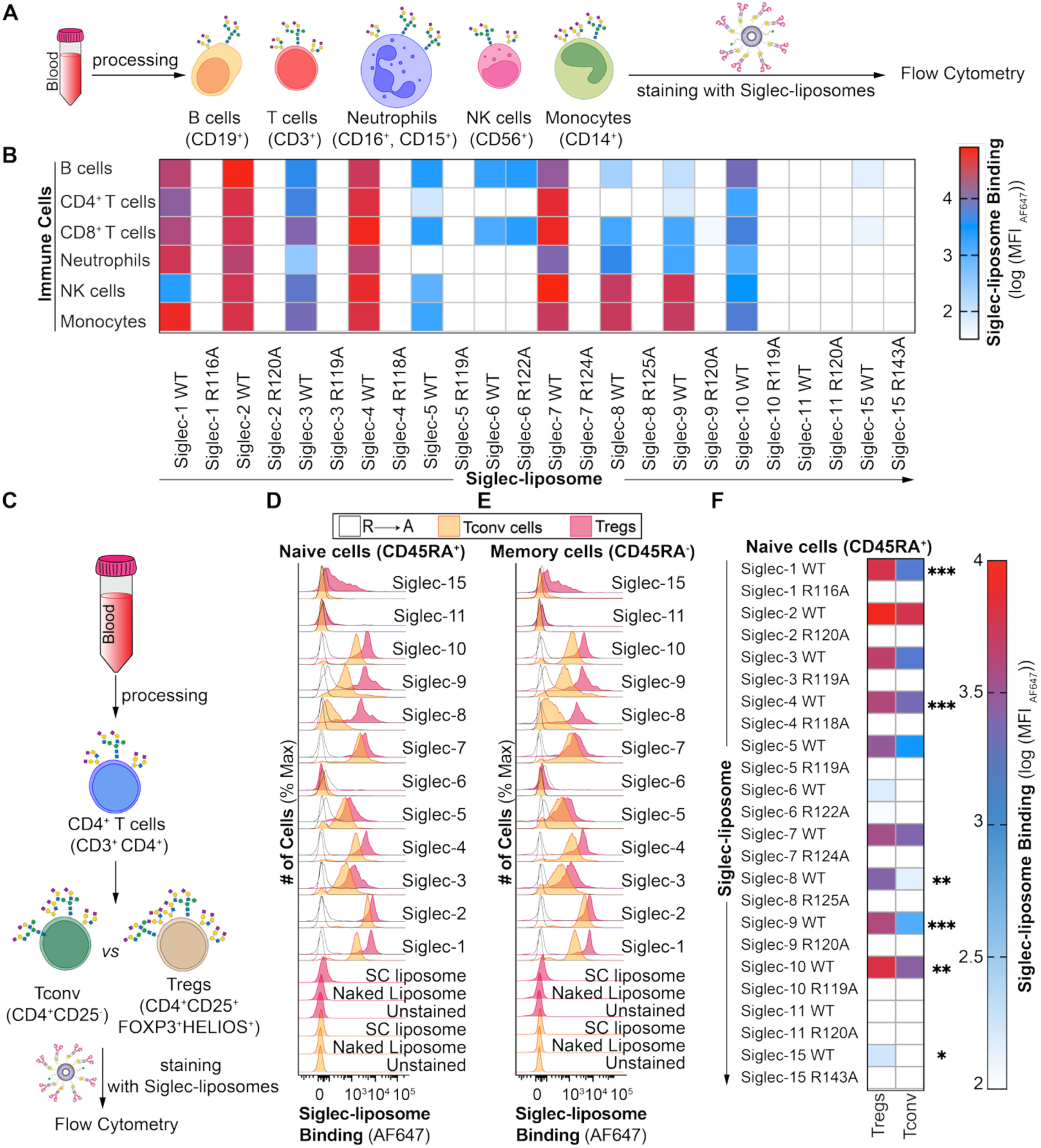
Profiling Siglec ligands on primary cells. (**A**) Schematic of staining immune cells isolated from human peripheral blood with Siglec-liposomes. (**B**) Siglec-liposomes binding to B cells, T cells, mature NK cells, monocytes, and neutrophils isolated from PBMCs. (**C**) Schematic of staining Treg and Tconv cells isolated from human peripheral blood with Siglec-liposomes. Binding of Siglec-liposomes to (**D**) naïve and (**E**) memory Treg and Tconv cells isolated from human peripheral blood. Data is presented as flow cytometry histograms. (**F**) Binding of Siglec-liposomes to naïve Treg and Tconv cells isolated from human peripheral blood. Data is presented as a heat map. The *p* value for four (panel B) and five (panel F) biological replicates was calculated using a paired one-way ANOVA. *0.05>*p* ≥0.01, **0.01>*p*≥0.001, ***0.001>*p*≥0.0001.

Profiling Siglec ligands on immune cells revealed that T cells have abundant Siglec ligands, prompting us to further investigate Siglec ligands on T cell subsets. We focused on Tregs, a subset of CD4^+^ T cells, because of their critical roles in maintaining immune homeostasis and preventing allergies, inflammation, and autoimmune diseases (*28*). Cancers exploit the suppressive function of Tregs by expanding these regulatory cells in the tumor microenvironment, thereby, inhibiting anti-tumor immunity (*29*). Notably, Tregs exhibit distinct glycosylation patterns compared to Tconv cells (*30, 31*) and efforts have been made to identify ligands of some Siglecs such as Siglec-1 on Tregs (*32*). To profile Siglec ligands on Tregs and compared them with Tconv cells, we isolated these cells from fresh peripheral blood of five individuals and stained them with Siglec-liposomes (**Fig. 3C**). Specifically, we compared the levels of Siglec ligands on naïve (CD45RA^+^) and memory (CD45RA^-^) Treg and Tconv cells (**Fig. 3D,E** and **S17**). Several Siglec-liposomes including Siglec-1, -8, -9, -10, and -15 showed increased binding to Tregs compared to Tconv cells and this difference was more pronounced in naïve (CD45A^+^) compared to memory (CD45A^-^) Tregs (**Fig. 3F**). As Tregs constitute ∼2-5% of periphery T cells, clinically relevant numbers of Tregs can be prepared for clinical and pre-clinical trials using *ex vivo* expanded Tregs (*28*). Therefore, we were curious if Siglec ligands on *ex vivo* expanded Tregs showed similar patterns, and profiled Siglec ligands on autologous *ex vivo* expanded Tregs and Tconv from five donors (**Fig. S18**). For the expanded Tregs and Tconv, only Siglec-10-liposomes showed higher binding to Treg compared to Tconv cells.

### Multiplexing Siglec-liposomes

The modular process by which Siglec-liposomes are formulated makes them suited for multiplexing in two different ways. The first way is to present different Siglecs on different liposomes that each contain a unique fluorophore to simultaneously detect multiple Siglec ligands in a single experiment (**Fig. 4A**). Accordingly, SC-liposomes were prepared with DSPE-PEG-AF488, -AF555, or -AF647, and conjugated with V2 Siglec-1, V2 Siglec-7, and V2 Siglec-2, respectively. These Siglecs were chosen because they have non-overlapping ligands (*16*). After conjugation, all three Siglec-liposomes were mixed and tested for binding to U937 cells by flow cytometry. In parallel, each individual Siglec-liposome was tested for binding to U937 cells. No significant difference in binding to cells was observed for AF488- Siglec-1-liposomes compared to when they were multiplexed with the other two liposomes (**Fig. 4B**). Similarly, binding of AF555-Siglec-7-liposomes and AF647-Siglec-2-liposomes showed no significant difference in binding compared to the multiplexed liposomes (**Fig. S19A,B**). We also created different combinations of multiplexed liposomes consisting of WT and arginine mutated Siglecs and tested their binding to U937 cells, which demonstrated that the presence of one Siglec does not interfere with the ability of another Siglec to engage their glycan ligands on cells (**Fig. 4C** and **S19C-H**).

**Fig. 4.**
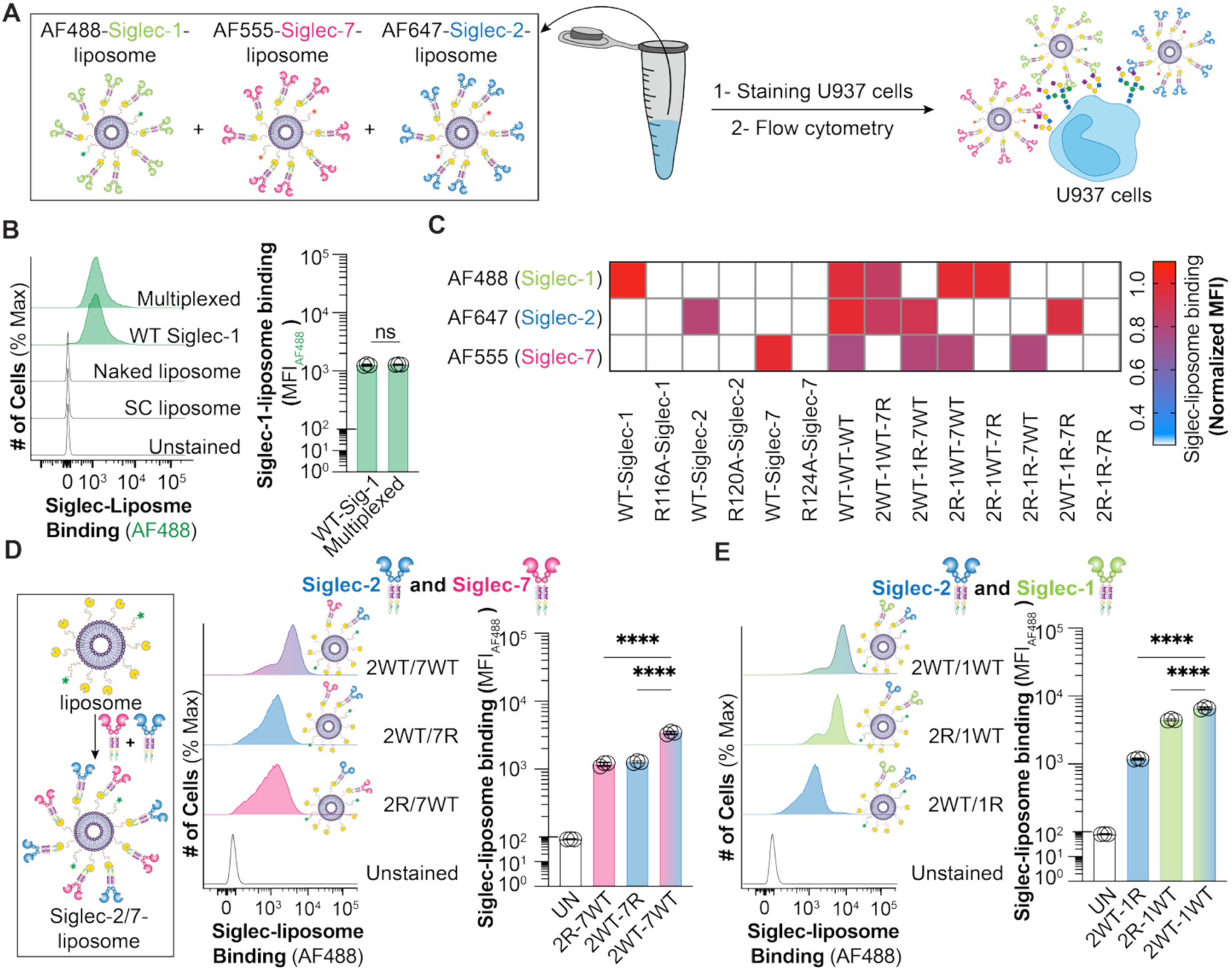
Siglec-liposomes as a multiplexing platform. (**A**) Schematic for multiplexing liposomes with different fluorophores. (**B**) Binding of single color and multiplexed of AF488-Siglec-1-liposome to U937 cells. Data is presented as flow cytometry histograms and MFI of Siglec binding. (**C**) Binding of multiplexed and single color of AF488-Siglec-1-, AF555-Siglec-7-, and AF647-Siglec-2-liposomes in different combinations to U937 cells. Data is presented as a heat map. Binding of (**D**) Siglec-2/Siglec-7, and (**E**) Siglec-1/Siglec-2 on the same liposomes to U937 cells. Data is presented as flow cytometry histograms and MFI of Siglec binding. The *p* value for three technique replicates was calculated using an unpaired one-way ANOVA. Not significant (ns) *p*>0.05, \*\*\*\**p*<0.0001.

The second way Siglec-liposomes were multiplexed is by displaying multiple Siglecs on the same liposome. To test this, we initially conjugated V2 Siglec-2 and V2 Siglec-7 on the same liposome and probe their binding to U937 cells via flow cytometry (**Fig. 4D**). Liposomes presenting V2 Siglec-2 and V2 Siglec- 7 bound to cells in a positively cooperative manner. Control multiplexed liposomes including liposomes displaying one Siglec (Siglec-2 or Siglec-7) or displaying WT and arginine mutated of these Siglecs were prepared and results demonstrate that Siglec-liposomes bind to their cellular ligands in all different combinations of multiplexed liposomes without interfering with each other (**Fig. S20A**). We also found that multiplexed V2 Siglec-1 and V2 Siglec-2 bound to their ligands on the cells in a positively cooperative manner (**Fig. 4E** and **S20B**). These results demonstrate that Siglec-liposomes can be used as a multiplexed platform to detect different Siglec ligands on the cell simultaneously or display different Siglecs on one liposome to mimic a genuine immune cell.

### In vivo compatibility of Siglec-liposomes

As the base liposome formulation used in our studies have a long *in vivo* circulatory half-life (*26, 33*), we felt they would be well suited for detecting their ligands in mice. We tested Siglec-2- and Siglec-7-liposomes as they show strong binding to immune cells isolated from mouse splenocytes (**Fig. S21**). As Siglecs expressed in CMAS^-/-^ CHO cells will have terminal galactose residues that can be taken up by the liver (*34*), we used V2 Siglec-2 and V2 Siglec-7 expressed From CMAS^+/+^ CHO cells to avoid this issue. Siglec-7-liposomes were formulated on SC-liposomes at different densities (0.003, 0.01, 0.03, and 0.1 mol%) and injected these into mice intravenously (**Fig. S22**). After one hour, mice were euthanized and binding of Siglec-7-liposomes to different immune cells in splenocytes were assessed by flow cytometry. Consistent with our *in vitro* staining of mouse immune cells with Siglec-7-liposomes, Siglec-7-liposomes showed *in vivo* binding to T cells in a manner that was dependent on the density of Siglec-7 on the SC-liposomes.

Two concentrations of Siglec-7-liposomes (**Fig. 5A** and **S23**) or Siglec-2-liposomes (**Fig. 5B** and **S23**), with 0.1 mol% V2 Siglec, were administered to mice. Mouse splenocytes were isolated and the binding of Siglec-liposomes to different immune cells including B cells (B220^+^), CD4^+^ T cells (CD3^+^, CD4^+^), CD8^+^ T cells (CD3^+^, CD8^+^), NK cells (NK1.1^+^), and neutrophils (LY6G^+^) were assessed by flow cytometry. WT Siglec-7-and WT Siglec-2-liposomes showed significant binding to T cells and B cells, respectively, at both 1.3 mM and 4 mM concentrations. In contrast, no binding was observed for R124A Siglec-7- or R120A Siglec-2-liposomes. The binding of Siglec-7-liposomes to T cells was also assessed in the spleen through immunofluorescence (IF) microscopy, which demonstrated clear co-localization with CD3^+^ T cells (**Fig. 5C** and **S24**).

**Fig. 5.**
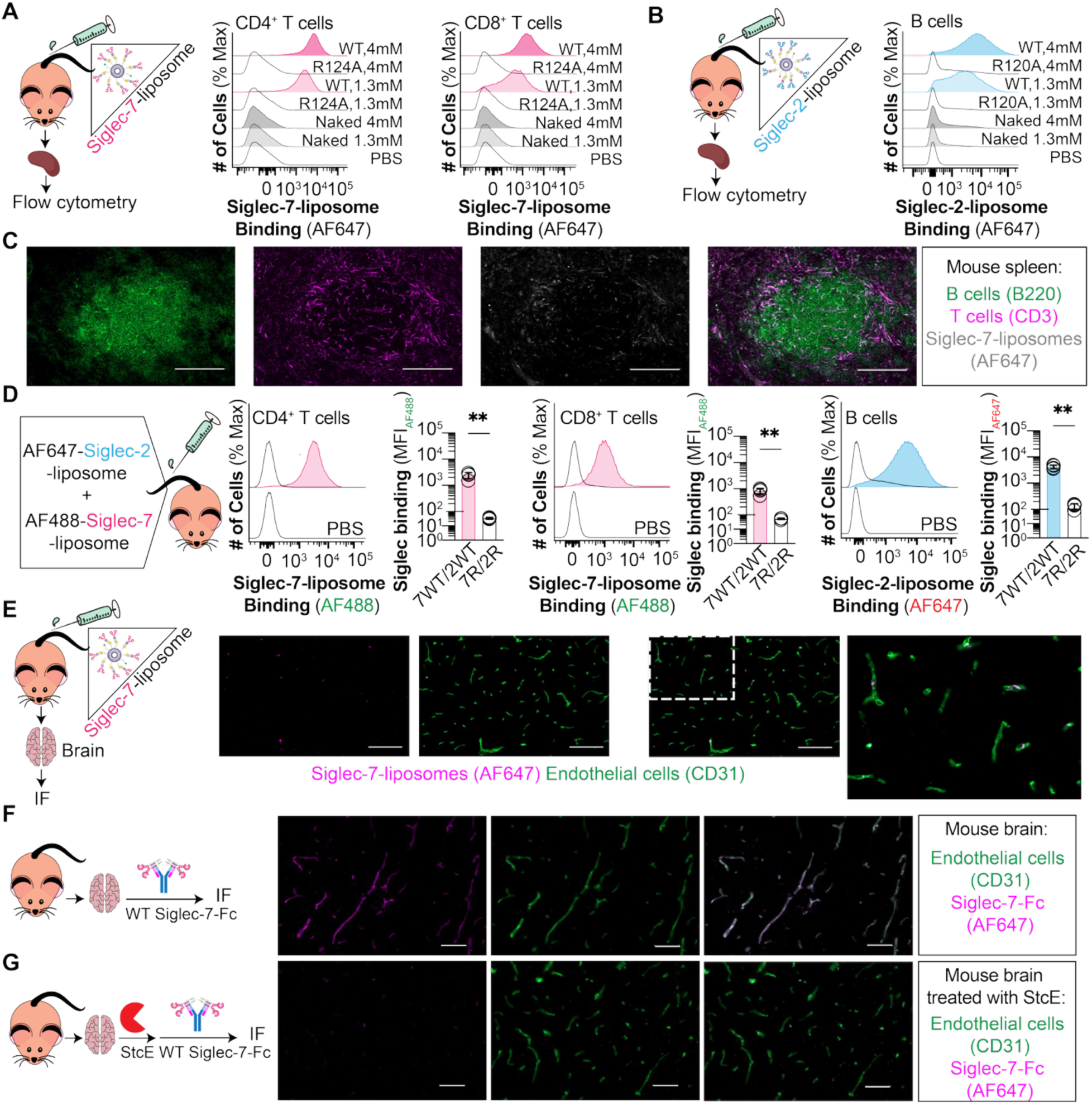
*In vivo* and *ex vivo* staining of cells and tissues using Siglec-liposomes. *In vivo* binding of WT and arginine mutated of (**A**) Siglec-7-liposome to CD4^+^ and CD8^+^ T cells and (**B**) Siglec-2-liposome to B cells isolated from mouse splenocytes. Data is presented as flow cytometry histograms. (**C**) IF microscopy images from mouse spleen representing *in vivo* binding of WT Siglec-7-liposomes to T cells. scale bar: 100 µm. (**D**) *In vivo* binding of multiplexed AF488-Siglec-7-liposomes to CD4^+^ and CD8^+^ T cells and AF647-Siglec-2-liposomes to B cells. Data is presented as flow cytometry histograms and MFI of Siglec binding. (**E**) IF microscopy images of mouse brain tissues representing *in vivo* binding of Siglec-7-liposomes to blood vessels. Scale bar: 100 µm. IF microscopy images of *ex vivo* staining of mouse brain tissues with (**F**) WT Siglec-7-Fc and αCD31, and (**G**) StcE mucinase enzyme treatment followed by WT Siglec-7-Fc and αCD31. Scale bar: 50 µm. The *p* value for four biological replicates was calculated using a paired Student’s *t* test. **0.01>*p*≥0.001.

We also assessed *in vivo* binding of multiplexed liposomes, where multiplexed Siglec-2-liposomes (AF647) and Siglec-7-liposomes (AF488) were administered into mice, and their binding to immune cells from mouse splenocytes were assessed (**Fig. 5D** and **S25**). Multiplexed WT Siglec-7- and WT Siglec-2- liposomes showed strong binding to T cells and B cells, respectively. Multiplexed R124A Siglec-7- and R120A Siglec-2-liposomes showed no binding to immune cells isolated from mice splenocytes (**Fig. 5D** and **S25**). This observation was consistent with our *in vitro* staining of immune cells from mouse splenocytes, where Siglec-7- and Siglec-2-liposomes showed strong binding to T cells and B cells, respectively, while arginine mutated of these Siglecs showed no significant binding.

In addition to spleen, we also examined Siglec ligands in other tissues. Mice injected with Siglec- 7-liposomes showed a pattern of fluorescent signal from the liposomes in the brain and kidney that resembled the vasculature. Therefore, we co-stained these tissues with αCD31 antibody to detect endothelial cells and found that CD31 staining overlapped with the signal from Siglec-7-liposomes, strongly suggesting that Siglec-7-liposomes bind to endothelial cells on blood vessels (**Fig. 5E** and **S26,27**). In follow-up *ex vivo* staining of mouse brain slices, Siglec-7-Fc and CD31 signal also overlapped (**Fig. 5F**), while no signal was observed for R124A Siglec7-Fc (**Fig. S28**). Mouse brain slices pre-treated with the mucinase StcE (*35*) demonstrated that the ability of Siglec-7-Fc to recognize vasculature was abolished (**Fig. 5G** and **S28**). Taken together, these results show that Siglec-liposomes enable Siglecs to detect Siglec ligands *in vivo* in both single color and multiplexed formats. Notably, Siglec-7-liposomes detect mucin ligands in brain vasculature.

### Immunomodulation by Siglec-liposomes

Antigen presenting cells (APCs) (dendritic cells, macrophages, and B cells) express Siglecs (*4*), and T cells express ligands for Siglec-1, -2, -3, -5, -7, -9, and -10, which suggest a functional role for Siglec-ligand interactions in cell-cell interactions between APCs and T cells. We felt that Siglec-liposomes would be a suitable tool to test this by determining if they can modulate T cell proliferation. Therefore, we isolated human T cells from peripheral blood of healthy donors and activated them with αCD3/28 beads along with co-culturing with Siglec-liposomes (**Fig. 6A**). As a control, T cells were co-cultured with PBS, naked liposome, SC liposome, or the arginine mutated versions of the Siglecs. We analyzed T cell proliferation by labeling CD4⁺ and CD8⁺ T cells with cell trace violet (CTV) fluorophore and measured fluorescence dilution by flow cytometry after 96 hours. Co-culturing T cells with liposomes displaying Siglec-3, -5, -9, or -10 did not affect their proliferation compared to controls (**Fig. S29A-F**). In contrast, liposomes displaying Siglec-7 significantly enhanced the proliferation of both CD4⁺ and CD8⁺ T cells (**Fig. 6B,C)**. The effects of Siglec-1- and Siglec-2-liposomes on T cell proliferation were donor-dependent, with some biological samples showing that Siglec-1- and Siglec-2-liposomes could stimulate T cell proliferation, but overall, these Siglec-liposomes did not significantly increase proliferation relative to controls (**Fig. S29G-J**). Co-culturing T cells with soluble Siglec-7-Fc increased the T cell proliferation, though to a lesser extent than Siglec-7-liposomes (**Fig. S30**).

**Fig. 6.**
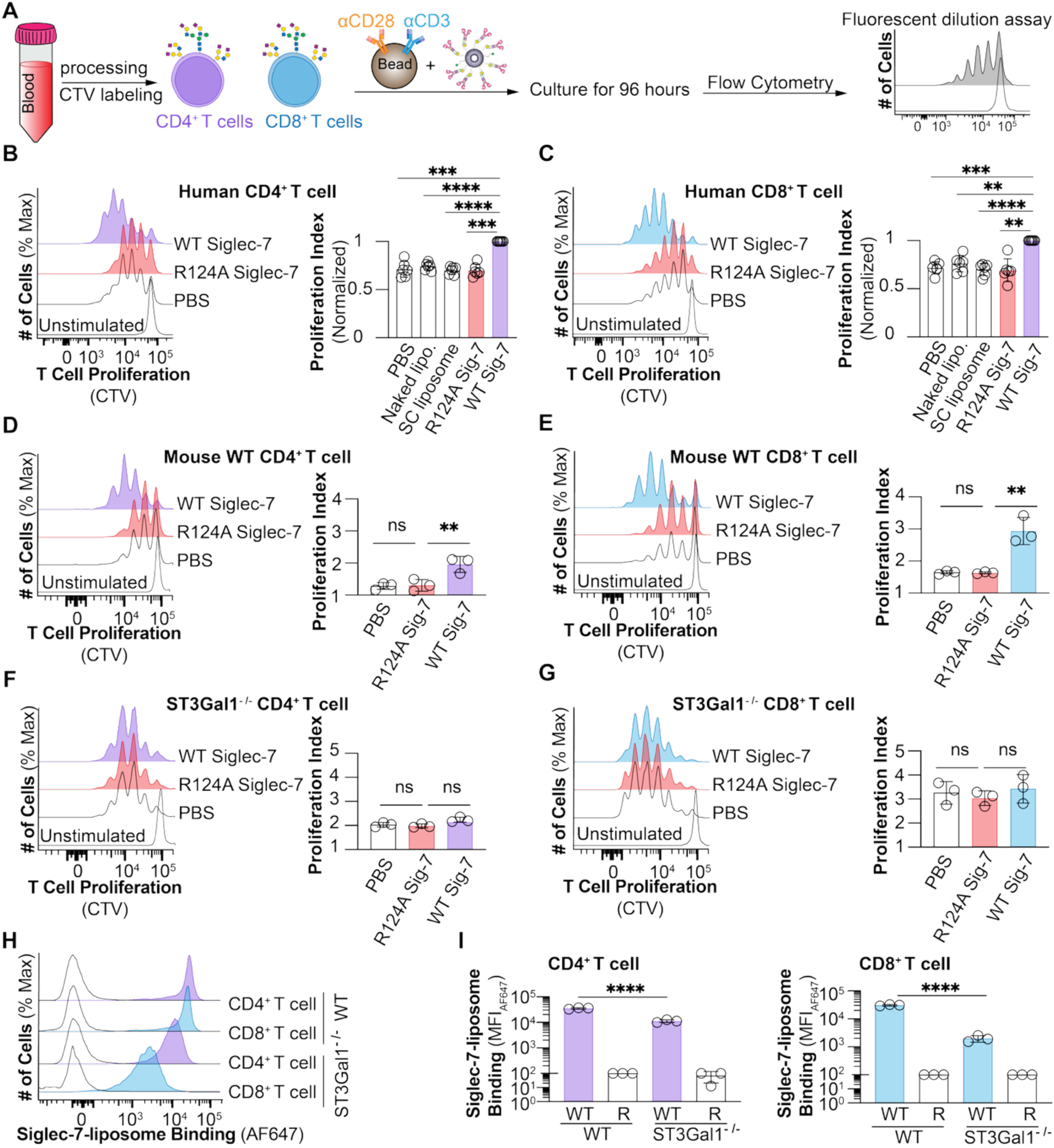
Modulation of T cell proliferation by Siglec-7-liposomes. (**A**) Schematic of T cell isolation from human blood and co-culturing with αCD3/28 beads and Siglec-liposomes. T cell proliferation of human (**B**) CD4^+^ and (**C**) CD8^+^ T cells co-cultured with WT and R124A Siglec-7-liposomes. Data is presented as flow cytometry histograms and proliferation index. T cell proliferation of (**D**) WT CD4^+^, (**E**) WT CD8^+^, (**F**) ST3Gal1^-/-^ CD4^+^, and (**G**) ST3Gal1^-/-^ CD8^+^ T cells isolated from mouse solenocytes. Data is presented as flow cytometry histograms and proliferation index. Binding of WT and R124A Siglec-7-liposomes to WT and ST3Gal1^-/-^ T cells. Data is presented as (**H**) flow cytometry histograms and (**I**) MFI of Siglec-7 binding. The *p* value for six (panel B,C) and three (panel D-G) biological replicates was calculated using a paired one-way ANOVA. The p value for three technical replicates was calculated using an unpaired ANOVA. Not significant (ns) *p*>0.05, **0.01>*p*≥0.001, ***0.001>*p*≥0.0001, \*\*\*\**p*<0.0001.

To study the effect of Siglec-7- liposomes in increasing T cell proliferation in more detail, we investigated the effect of Siglec-7-liposomes in T cells isolated from mouse spleen and found that they had a similar stimulatory effect as human T cells (**Fig. 6D,E**). ST3Gal1 is a major contributor to the biosynthesis of sialylated mucins, and mucin-type *O*-glycans are strongly implicated as ligands for Siglec-7 (*16*). Therefore, we tested the ability of Siglec-7-liposomes to stimulate proliferation in T cells from ST3Gal1^-/-^ mice (*36*). To do this, we isolated CD4^+^ and CD8^+^ T cells from WT and ST3Gal1^-/-^ mice, co-cultured them with WT or R124A Siglec-7-liposomes, and assessed T cell proliferation. Compared to T cells from WT mice, T cells from ST3Gal1^-/-^ mice displayed a hyperproliferation phenotype in both CD4^+^ and CD8^+^ T cell subsets (*36*), which was previously attributed to enhanced apoptosis of these cells. Therefore, we investigated T cell proliferation after 72 hours and found that Siglec-7-liposomes did not increased T cell proliferation in T cells from ST3Gal1^-/-^ mice (**Fig. 6F,G**). We also analyzed T cell proliferation in WT mouse CD4^+^ and CD8^+^ T cell after 96 hours and observed that the effect of Siglec-7-liposomes on T cell proliferation remained consistent between 72 and 96 hours (**Fig. S31**). To test the binding of Siglec-7- liposomes to ST3Gal1^-/-^ vs WT mice, we stained the T cells from WT and ST3Gal1^-/-^ mice with Siglec-7- liposomes and found that Siglec-7-liposomes bound to ST3Gal1^-/-^ T cells 3-fold less to CD4^+^ T cells and 15-fold less to CD8^+^ T cells, suggesting that ST3Gal1 plays a major role in generating ligands for Siglec-7 in mouse T cells (**Fig. 6H,I**). Nevertheless, it is interesting that there are ST3Gal1-independent Siglec-7 ligands, particularly on CD4^+^ T cells.

CD43 is a glycoprotein that has been demonstrated to be a carrier of ligands for Siglec-7 (*25, 37–39*). To test if CD43 contributes to the immunostimulatory role we observed for Siglec-7-liposomes, we isolated CD4^+^ and CD8^+^ T cells from WT and CD43^-/-^ mice along with co-culturing them with WT and R124A Siglec-7-liposomes. The T cell proliferation was analyzed after 96 hours (**Fig. S32A-D**). Analyzing T cell proliferation of CD4^+^ and CD8^+^ T cell from WT and CD43^-/-^ mice demonstrated that WT Siglec-7- liposomes increased T cell proliferation in CD4^+^ and CD8^+^ T cell subsets from both WT and CD43^-/-^ mice, while no effect was observed for R124A Siglec-7-liposomes. We confirmed that Siglec-7-liposome binding was modestly decreased on CD43^-/-^ T cells compared to T cells from WT mice (**Fig. S32E,F**). Overall, these results implicate an ST3Gal1-dependent and CD43-independent, mechanism by which Siglec-7- liposomes enhance T cell proliferation.

## Discussion

Similar to other glycan-binding proteins, multivalency is a key factor in Siglec-glycan interactions (*3, 40*). Siglec-liposomes enable multivalency to be experimentally controlled, which was accomplished by robust site-specific conjugation using the SpyCatcher-SpyTag system (*24*). Highly multivalent presentation of Siglecs enabled greatly improved detection of Siglec ligands on cells and tissues. A key feature that helped unlock the full potential of Siglec-liposomes was the expression of recombinant Siglecs from CHO cells that lack sialic acid. Production of Siglec-Fc proteins from Lec2 CHO cells or treatment of Siglec-Fc proteins with neuraminidase is carried out in some laboratories to prevent undesired *cis* ligand interactions that might mask the ability of Siglec-Fc proteins to recognize *trans* ligands on cells (*41*). However, there is limited experimental evidence outside the context of a cell surface that such masking is at play. We provide clear evidence that sialic acid-deficient Siglec-liposomes are much more capable of binding *trans* ligands on cells, indicating that masking of Siglecs by *cis* ligands is present on a multivalent platform such as the liposomes.

Although Siglec-Fc proteins are dimeric, they require further multimerization to detect ligands on cells, with ahuman IgG pre-complexing being the most common approach (*3*). Siglec-liposomes avoid the use of ahuman IgG and, therefore, avoids detection of adsorbed IgG on human samples. This feature enabled a detailed investigation of Siglec ligands on primary human immune cells. Each Siglec-liposome showed a unique pattern for binding to human immune cells. For example, we observed strong binding of Siglec-2-liposomes to B cells, Siglec-7-liposomes to T cells and NK cells, Siglec-9-liposomes to NK cells and monocytes, and Siglec-10 liposomes to B cells. Results with Siglec-9 and -10 were not observed in our previous study using pre-complexed Strep-Tactin (*5*), likely reflecting the enhanced sensitivity of Siglec- liposomes. It is intriguing that in these examples, the cell types expressing high levels of specific Siglec ligands also typically express that Siglec at high levels (*4*). These results are consistent with the ability of *cis* ligands to regulate the function of Siglecs; for instance, masking of the Siglec to only allow for *trans* ligand interactions under the appropriate physiological circumstance, such as within an immunological synapse (*42–45*).

The ability of *trans* interactions between Siglecs and their ligands on other cells to regulate immunobiology is quickly gaining recognition as a major role for Siglecs (*19, 37*). For example, numerous lines of evidence have been put forward for Siglecs inducing effects in T cells (*37, 46, 47*). Our results with Siglec-liposomes demonstrate that Siglec-7 engaging its ligands on T cells can stimulate T cell proliferation. It was previously demonstrated that Siglec-7-Fc inhibits T cell proliferation (*37*), which is the opposite of what we observed. Differences between the studies that could explain this are the enhanced multivalency of liposomes and the precise T cell stimulation protocol; the previous study used soluble aCD3/28 antibodies to activate T cells (*37*), while we activated T cells using aCD3/28 antibodies adsorbed to beads. Previous studies demonstrated that α2-3 linked sialic acids on mucin-type *O*-glycans serve as ligands for Siglec-7 (*1, 16*). While ST3Gal1^-/-^ T cells did not completely abrogate Siglec-7 ligands, the T cell stimulatory effect of Siglec-7-liposomes was completely abolished, demonstrating that the increase in T cell stimulation by Siglec-7-liposomes is mediated by ligands created by ST3Gal1. It is interesting that ST3Gal1^-/-^ CD8^+^ T cells were hyperproliferative, which has not been previously reported, but may be directly connected to the enhanced rate of apoptosis observed previously on ST3Gal1^-/-^ CD8^+^ T cells (*36*). CD43 is a glycoprotein ligand for Siglec-7 on K562 CML (chronic myeloid leukemia) cells (*25, 38*), healthy human T cells (*37*), and CLL (chronic lymphocytic leukemia) cancer cells (*39*). CD43^-/-^ T cells displayed moderately attenuated Siglec-7-liposome binding, confirming the previous observations that CD43 is a carrier for Siglec-7 ligands. However, CD43^-/-^ T cells are still susceptible to the stimulatory effect of Siglec- 7-liposomes, demonstrating that other ST3Gal1-dependent ligands mediate this stimulatory effect. Aside from CD43, other Siglec-7 ligands have also been identified, such as CD162 and CD45 (*37, 39*).

While our results demonstrate that Siglec-liposomes can modulate T cell proliferation, the implications of these findings within the context of an immunological synapse between a T cell and APC (*48*) are complex because Siglec ligands and receptors that activate T cells (*e.g*. TCR) will be co-engaged.

One study found that Siglecs on APCs inhibit T cell responses (*49*). However, assessing the impact of Siglecs in the complex environment of an immunological synapse is not straight-forward and, in this regard, APCs express diverse and broad levels of Siglecs (*4*). For example, macrophages express Siglec-1 and -15, dendritic cells express Siglec-2, -3, -5, -7, -9, and -15, and B cells express Siglec-2 and -10. Our liposome platform is suited to exploring this in greater detail in future studies by combining Siglecs and molecules that activate T cells in order to co-engage/co-cluster these receptors on T cells. The implications of our work on Siglec-liposomes inducing T cell proliferation are also intriguing considering our observation that Tregs express higher levels of Siglec ligands compared to Tconv cells, including Siglec-1, -8, -9, -10, and -15, and that roles for Siglecs engaging their ligands on Tregs have been proposed (*32*). It was previously demonstrated that compared to Tconv cells, Tregs have altered expression of *N*-glycans (*50, 51*), which is relevant because many Siglecs (Siglec-1, -9, -10, and -15) recognize *N*-glycans as their sialoglycan ligands (*15, 16, 52*). Consistent with our findings for Tregs isolated from PBMCs, *ex vivo* expanded Tregs showed more ligands for Siglec-10, however, we did not observe an increase in binding of the other Siglecs in *ex vivo* expanded Tregs compared to Tconv cells. This difference may be because the *ex vivo* expanded Treg and Tconv cells were derived from human thymus, which may differ from Treg and Tconv cells isolated from human blood (*28*). As tumor associated macrophages express high levels of Siglec-10 (*53, 54*), our observation of higher levels of Siglec-10 ligands on Tregs may have important implication in maintaining an immunosuppressive tumor microenvironment.

A major advantage of Siglec-liposomes is their ability to be multiplexed, which we demonstrated in two ways: (i) mixing Siglec-liposomes with different fluorophores or (ii) creating liposomes with different Siglecs on the same liposome. The ability to study multiple Siglecs functioning together is intriguing in light of differences in glycan specificity, with most Siglecs recognizing a non-overlapping set of sialoglycan ligands (*4*). For our multiplexing studies, we used Siglec-1-, Siglec-2-, and Siglec-7- liposomes because these Siglecs have non-overlapping glycan specificity: Siglec-1 prefers a2-3-linked sialosides of many classes of glycans (*55–57*), Siglec-2 prefers a2-6-linked sialosides on *N*-glycans (*42, 58*), and Siglec-7 prefers sialylated mucin-type *O*-glycans (*16, 37, 59*). Indeed, our results demonstrated that binding of these Siglecs to their respective glycan ligands on cells is not competitive in both types of multiplexing. In fact, co-presenting Siglec-2 and Siglec-7, or Siglec-1 and Siglec-7 on one liposome showed positive cooperative binding. In principle, multiplexing of Siglec-liposomes with different fluorophores is limited by the availability of small molecule fluorophores, although spectral flow cytometry makes it feasible to use many more fluorophores. An intriguing approach warranting future investigation is a genetic barcode associated with each liposome that could combine the advantages of working with bacteriophage and the advantages of the *in vivo* compatibility of liposomes.

Most immune cells express multiple Siglecs, and an open question is how Siglecs function together to recognize *trans* ligands. For example, NK cells express Siglec-7 and Siglec-9 (*60, 61*), while B cells express Siglec-2 and Siglec-10 (*42, 53, 54, 62, 63*). Our modular liposome platform provides an opportunity to address this physiologically relevant question, which would not be possible to systematically address with previously developed approaches to pre-complex and detect Siglecs. In the future, this multiplexing approach can be highly useful to investigate the functional impact on immune cells, like the immunostimulatory effect we observed on T cells for individual Siglecs.

Most pre-complexed forms of Siglecs involve non-covalent assembly with ahuman IgG or Strep-Tactin, which are not suitable for *in vivo* applications. Recently, we demonstrated that Siglecs can be displayed on genetically-encoded bacteriophage for *in vivo* applications (*8*). However, the extremely short circulatory half-life of bacteriophage (*64*) limits *in vivo* applications using this platform. Liposomes are commonly used for different *in vivo* applications such as mice immunization (*62, 65*), immunomodulation (*62*), and drug delivery in the laboratory mouse (*66, 67*). Indeed, we demonstrated that Siglec-7-liposomes and Siglec-2-liposomes showed robust *in vivo* binding to T cells and B cells from mice, respectively. Moreover, we demonstrated that these two liposomes can be multiplexed with different fluorophores to detect their ligands on T and B cells within the same mouse. Recently, it was shown that brain vasculature has a dense array of mucins (*68*). Our findings that Siglec-7-liposomes bind to endothelial cells in the brain, via mucins, are highly consistent with these findings. A potential application of these findings would be active delivery of a drug to the blood brain barrier (BBB). Moreover, as engagement between Siglecs and their ligands on endothelial cells can induce *trans* signaling (*69, 70*), another application for Siglec-7- liposome would be modulating the function of endothelial cells. We speculate that this could relate to a natural role for Siglec-7 in modulating the function of endothelial cells, which is particularly intriguing in light of diminished mucins on the BBB in the context of neurodegeneration (*68*).

In summary, we have developed Siglec-liposomes as a multivalent, modular, and versatile platform that enables the detection and functional manipulation of Siglec ligands on cells. Most importantly, Siglec- liposomes are highly sensitive and can be used to detect their glycan ligands *in vivo*. Moreover, owing to the modular assembling of Siglec-liposomes, multiplexing enables multiple Siglec ligands to be detected simultaneously. Having a complete set of human Siglecs with a genetically-encoded SpyTag will facilitate more innovative approaches in the future.

## Materials and Methods

### Animal

All mice used in this study were on a C57BL/6J genetic background. Wild type B6J mice (The Jackson laboratory) were bred in house. The CD43^-/-^ and ST3Gal1^-/-^ mice were provided to us by Dr. Som G. Nanjappa (University of Illinois Urbana-Champaign, USA) and Dr. Jamey Marth (Sanford Burnham Prebys, USA), respectively. The animals used in this study were bred and maintained under specific pathogen-free conditions. All animal procedures described in the study were approved by the University of Alberta Animal Care committee in accordance with the Canadian Council on Animal Care (CCAC) guidelines (#AUP00002885).

### Experiments using human blood

All experiments were approved by the human research ethics board (REB Pro00092144) biomedical panel at the University of Alberta and the University of British Columbia (REB H17-01490). Donors used in this study aged between 20-50 years old, were non-smoker, and were in good overall health.

### Cell lines and cell culture

Chinese hamster ovary (CHO) cells were cultured in DMEM/F12 media (ThermoFischer), supplemented with 5% (v/v) heat-inactivated fetal bovine albumin (Gibco), 100 U/mL penicillin (Gibco), and 100 µg/mL streptomycin (Gibco). Human monocytic U937 cells (ATCC) were grown in RPMI 1640 (ThermoFischer), supplemented with 10% (v/v) heat-inactivated fetal bovine albumin, 100 U/mL penicillin, and 100 µg/mL streptomycin. Cells were grown at 37 °C in 5% carbon dioxide (CO_2_) incubator. To generate *ex vivo* expanded Treg and Tconv cells, human thymuses were processed and T cells were isolated, stimulated and expanded as previously described.(*71, 72*) Cells were cryopreserved 3 days after restimulation on day 11 of the expansion. *Ex vivo* expanded Tregs and Tconv cells were cultured in ImmunoCult™-XF T Cell Expansion Medium (Stemcell Technologies) with 100 IU/mL of hIL2 (Stemcell Technologies).

### Cloning of Version 1 and Version 2 constructs

To make version 1 (Siglec-ST-Fc) and version 2 (Siglec-Fc-ST), two different plasmid backbones were generated. For the version 1, two successive PCR reactions were performed. The first PCR reaction was performed using primers P1 and P2 (**Table S1**), in which the pcDNA-5 containing Fc, Strep Tag II and His-tag was used as a template. The PCR product size was validated using a 1 % agarose gel and was excised from the gel. The PCR product was then purified using the GeneJET Gel Extraction Kit (Thermo Fisher). The second PCR reaction was performed using primers P2 and P3 (**Table S1**), and used the PCR product form the 1^st^ PCR as a template. An 1 % agarose gel was used to validate the size of the 2^nd^ PCR. The PCR product was then excised and cleaned up as described above. For the version 2, one PCR reaction was performed to add the SpyTag peptide to the C-terminus of the Fc contract using primers P4 and P5 (**Table S1**) and pcDNA-5 containing Fc, Strep Tag II and His_6_-tag as a template. The size of the PCR product was validated by using an 1 % agarose gel and it was cleaned up as described above. In both Version 1 and 2, *AgeI* and *Xmal* restriction sites were added to 5’ and 3’ ends, respectively. Both final PCR products for the Versions 1 and 2 were treated with *AgeI* and *Xmal* restrictions enzymes (New England Biolabs) for 1 hour at 37 °C. The double digested products were then run on a 1 % agarose gel and the gene was excised and cleaned up as described above. To insert Version 1 and 2 into pcDNA5, the plasmid was treated with *AgeI* restriction and phosphatase enzymes. The Version 1 and 2 were then ligated into the pcDNA-5 and transformed into chemically competent *Escherichia coli* DH5α (New England Biolabs). Colonies were then picked and grown in liquid culture (lysogeny broth plus 100 μg/mL ampicillin) overnight in a shaking incubator at 37 °C. The plasmids were then purified using the GeneJET Plasmid Miniprep Kit (Thermo Fisher). The successful incorporation of Versions 1 and 2 were then confirmed by restriction enzyme digestion and Sanger sequencing.

The pcDNA-5 plasmids containing genes of the WT and arginine mutant of Siglec-1, -2, -3, -4, -5, -6, -7, -8, -9, -10, -11, and -15 from the 1^st^ generation of Siglec-Fc-Strep Tag II(*5*) were treated with *NheI* and *AgeI* restrictions enzymes (New England Biolabs) for 1 hour at 37 °C to yield the Siglec genes containing 5’ *NheI* and 3’ *AgeI* restriction sites. The restriction digestion was confirmed by using an 1 % agarose gel. The double digested genes were then excised and purified as above. To incorporate the Siglec genes into pcDNA5 containing Fc and ST, the plasmid was digested with *NheI* and *AgeI* enzyme, ran on an 1 % agarose gel and the double cut plasmid was excised from the gel. WT Siglec-7 and R124A Siglec-7 were then ligated into both pcDNA5-Fc-ST and pcDNA5-ST-Fc. The plasmids were then transformed, grown up, and purified as described above. The successful incorporation of Siglec-7 (WT and R124A) was then confirmed by restriction enzyme digestion and Sanger sequencing. All other above-mentioned Siglecs in both WT and arginine mutated versions were ligated into pcDNA5-Fc-ST and transformed, grown up, purified, and confirmed by Sanger sequencing, as described above.

### Generation of CMAS^-/-^ CHO Flip-In cells using CRISPR/Cas9 gene editing

Custom crRNA (Integrated DNA Technologies; IDT) was designed to target exon 5 of hamster CMAS (sequence = ATTGATGGATGTCTCACCAA). CHO Flip-In cells were seeded at 500,000 cells per well the day of transfection in a 6-well tissue culture plate, in 1.3 mL growth media (DMEM/F12 containing 5% (v/v) heat- inactivated fetal bovine albumin (Gibco), 100 U/mL penicillin (Gibco), and 100 µg/mL streptomycin (Gibco)). For one well of a 6-well plate, 20 pMol of Cas9 nuclease (IDT), 20 pMol of ATTO-647 labeled crRNA:tracrRNA (IDT) duplex, 8µL Cas9 Plus reagent (Thermo Fisher), and 16 μL CRISPRMAX reagent (Thermo Fisher) in 665 μL Opti-MEM medium (Thermo Fisher), were incubated together for 15 minutes, and then added to the cells. One day after transfection, cells were removed from the plate, washed, resuspended in 300 μl of cell sorting medium (PBS, 1% FBS, 1 mM EDTA), and stored on ice until sorting. Cells were sorted using a FacsMelody cell sorter (BD). The brightest 10% of cells stained with ATTO-647 were sorted as single cells into 96-well plates containing regular culture medium. Cells were grown for two weeks until colonies were large enough to be screened. For the screening, CMAS^-/-^ clones were screened with diCBM40^-^ and Peanut agglutinin (PNA^+^). The clones which were PNA^+^ and diCBM40^-^ were then expanded. Relevant portion of the genome was amplified by PCR using P6 and P7 primers (**Table S1**), and CRISPR^-/-^ was validated by Sanger sequencing of the PCR product.

### Stable Siglec-Fc transfection

WT or CMAS^-/-^ CHO Flip-In cells were cultured as described above. To stably transfect recombinant Siglec-Fcs in CHO cells, 400,000 cells were seeded per well in a 6-well cell culture dish. The following day, 2 µg of PcDNA-5-Siglec-Fc plasmid, 2 µg of pOG44 plasmid, and 4.7 µL of Lipofectamine plus reagent (Thermo Fisher) were added to 500 µL of Opti-MEM media (Gibco) and the mixture was incubated at room temperature for 15 minutes. Next, 15.7 µL of Lipofectamine LTX reagent (Thermo Fisher) was added to the mixture and incubated for 30 minutes at room temperature. During the incubation of the DNA mixture, cells were washed with 1.5 mL of Opti-MEM media (Gibco). After 30 minutes, the DNA mixture was added to the cells and cells were left in the growing conditions described above. In the next day, 2 mL of DMEM/F12 media with the above-mentioned supplements was added to cells. The cells were then selected by gradually increasing the Hygromycin (Thermo Fisher) concentration (from 0.5 mg/mL to 1 mg/mL) for two weeks. Media for the cells was also replaced every other day.

### Recombinant Siglec-Fc expression and purification

Cells were cultured in three 15 cm dishes. Once cells reached confluency, they were distributed into ten 15 cm dishes and placed at 32 °C incubators with 5% CO2. Ten days after cells reached confluency, the media was then harvested, put through a 0.22 µm filter, and stored at 4 °C for later purification. The Siglec-Fcs were purified using a Ni^2+^-Affinity chromatography (Cytiva) and Strep-Tactin affinity column (iba Lifescience). For the Ni^2+^-Affinity chromatography, the column was first equilibrated with 20 CV (column volumes) of equilibrium buffer (20 mM sodium phosphate, 0.5 M NaCl, pH 7.4). Next, the supernatant was loaded on the column, followed by 20 CV of wash buffer (50 mM imidazole, 20 mM sodium phosphate, 0.5 M NaCl, pH 7.4). The protein was then eluted using 20 CV of elution buffer (500 mM imidazole, 20 mM sodium phosphate, 0.5 M NaCl, pH 7.4) into fractions of 1 mL. Fractions were then combined and diluted 10X with buffer W (100 mM Tris-HCl, 150 mM NaCl, 1 mM EDTA, pH 8). The Strep-Tactin affinity column was equilibrated with 15 CV of buffer W, following by loading the protein. Next, column was washed with 15 CV of buffer W and the protein was eluted using 15 CV of elution buffer (100 mM Tris-HCl, 150 mM NaCl, 1 mM EDTA, 10 mM Desthiobiotin, pH 8). Fractions were then combined and dialyzed against 1X PBS, pH 7 overnight. The following day, protein was concentrated using AMICON Ultra Centrifugal Filters (Thermo Fisher). Protein’s size was validated using SDS-PAGE and protein concentration was then measured using a BCA assay (Thermo Fisher). The proteins were then either stored at 4 °C or lyophilized.

### Production of Version 1 construct

To yield the monomeric Siglec-ST, the purified protein was incubated with 10 molar excess of TEV protease enzyme overnight at 4 °C. To remove Fc and TEV, Ni^2+^-Affinity chromatography was used (Cytiva). The column was first equilibrated with 20 CV (column volume) of equilibrium buffer (20 mM sodium phosphate, 0.5 M NaCl, pH 7.4) and then protein was loaded onto the column. The flowthrough of the column containing Siglec-7-ST was collected and dialyzed against 1X PBS, pH 7 overnight. The protein was then concentrated using AMICON Ultra Centrifugal Filters (Thermo Fisher). The protein’s size was validated by SDS-PAGE and the protein concentration was measured using a BCA assay. Protein was then stored at 4 °C.

### Expression and purification of S49C SpyCatcher

S49C SpyCatcher plasmid was transformed into chemically competent E. coli BL21 (DE3) cells (New England Biolabs). The cells were then cultured overnight into 100 mL of liquid culture (lysogeny broth plus 100 μg/mL ampicillin). The following day, the culture was added to two 500 mL of liquid cultures (lysogeny broth plus 100 μg/mL ampicillin) and incubated at 37 °C, 220 rpm shaking until the OD reached 0.6. Once OD∼0.6, the S49C SpyCatcher expression was induced using a final concentration of 1mM IPTG and incubated at 37 °C, 220 rpm for 3 hours. Cells were then collected and centrifuged at 3,400 *xg* for 20 minutes, 4 °C, and the pellets were resuspended in binding buffer (50 mM Tris, 500 mM NaCl, 5 mM imidazole, pH 8) with 300ug/mL lysozyme and incubated for one hour on ice. Cells were then sonicated using constant pulse for 15 seconds on and 45 seconds off on ice for ten times. Cells were centrifuged for 35 minutes at 18,000 *xg* and at 4 °C. The supernatant was then filtered and stored at 4 °C for purification. For protein purification, a 5 mL Ni^2+^-affinity chromatography column (Cytiva) was equilibrated with 15 column volumes of binding buffer, and the supernatant was then loaded onto the column. The column was then washed with 100 mL of wash buffer (50 mM Tris, 500 mM NaCl, 60 mM imidazole, pH 8). The S49C SpyCatcher protein was then eluted from the column using elution buffer (50 mM Tris, 500 mM NaCl, 200 mM imidazole, pH 8) into 1 mL fractions. The fractions containing protein were then combined and the protein dialyzed against 1X PBS, pH 7 overnight at 4 °C. The protein was then concentrated using AMICON Ultra Centrifugal Filters and the size of the protein was validated by SDS-PAGE. The concentration of the protein was then validated using a BCA assay.

### Expression and purification of StcE enzyme

StcE plasmid was transformed into chemically competent E. coli BL21 (DE3) cells (New England Biolabs). The cells were then cultured overnight into 100 mL of liquid culture (lysogeny broth plus 100 μg/mL kanamycin). The following day, culture was added to 1L liquid culture (lysogeny broth plus 100 μg/mL kanamycin) incubated at 37 °C, 220 rpm shaking until it reached OD 0.6. Once OD∼0.6, the StcE expression was induced using a final concentration of 1mM IPTG and incubated at 16 °C, overnight. The following day, cells were centrifuged at 3,000 *xg* for 35 minutes, at 4 °C, and the pellets were collected and resuspended in binding buffer (50 mM Tris, 500 mM NaCl, 5 mM imidazole, pH 8) with 300 ug/mL lysozyme and incubated for one hour on ice. Cells were then sonicated using constant pulse for 15 seconds on and 45 seconds off on ice for ten times. Cells were centrifuged for 35 minutes at 18,000 *xg* and at 4 °C. The supernatant was then filtered and stored at 4 °C for purification. A 5 mL Ni^2+^-affinity chromatography column (Cytiva) was equilibrated with 15 column volumes of binding buffer; the supernatant was then loaded onto the column. The column was then washed with 100 mL of wash buffer (50 mM Tris, 500 mM NaCl, 60 mM imidazole, pH 8). The StcE protein was then eluted from the column using elution buffer (50 mM Tris, 500 mM NaCl, 200 mM imidazole, pH 8) into 1 mL fractions. The fractions containing protein were then combined and the protein dialyzed against 1X PBS, pH 7 overnight at 4 °C. The protein was then concentrated using AMICON Ultra Centrifugal Filters and the size of the protein was validated by SDS-PAGE. The concentration of the protein was then validated using a BCA assay.

### Spycatcher liposome preparation

Lipids including, 57 mol% 1,2-distearoyl-sn-glycero-3-phosphocholine (DSPC) (Avanti, CAS: 816-94-4), 38 mol% cholesterol (Avanti, CAS: 57-88-5), and 5 mol% polyethylene glycol distearoylphosphatidylethanolamine (PEG_45_–DSPE MW 2000) (Avanti, CAS: 474922-77-5) were dissolved in chloroform to yield the desired mol% used in liposome formulation. After dissolving lipids, the chloroform was evaporated using nitrogen gas to obtain a lipid thin film. Once dry, 100 µL of DMSO and the appropriate amount of fluorophore conjugated with DSPE-PEG-2000(*7*) was added to the lipid thin film from their respect DMSO solution, and it was stored at -80 °C until completely frozen. The DMSO was then removed through lyophilization.

### Liposome extrusion

Lipid thin film was hydrated with a total amount of 1 mL of 1X PBS and the appropriate amount of SpyCatcher-DSPE-PEG. The lipids in 1X PBS were then sonicated in a sonicator for cycles of one min on, five min off, for five times. Lipids were then extruded using an Avanti mini extruder in 1000µl syringe (SKU: 610017-1Ea), first with an 800 nm filter (cytiva, 10417304) and then with a 100 nm filter (cytiva, 800309), each for 20 times each to yield 100 ± 35 nm liposomes. The liposomes hydrodynamic diameter was then measured by a Malvern Panalytical Zetasizer Nano S. Liposomes were then stored at 4 °C for usage.

### Preparation of Siglec-liposomes and Siglec-Strep-Tactin

Lyophilized Siglec-Fcs, either 1^st^ generation (Siglec-Fc-Strep Tag II) or Version 2 (Siglec-Fc-ST), were hydrated using nuclease free water. Siglecs were then added to AF647-Strep-Tactin and AF647-SC-liposomes with the molar ratio of 1:5 and 1:1.2 (monomer Strep-Tactin: Siglec-Fc and SpyCatcher: Siglec-Fc), respectively, where the final concentration of Siglec-Fc was 16 µg/mL. The pre-complexing of Siglec-Strep-Tactin was performed for 30 minutes on ice and all Siglec-liposome conjugation was performed for one hour at room temperature.

### Flow cytometry assays on cell lines

To stain the cells, 200,000 cells/well were plated into 96-well U-bottom microplates and centrifuged for 5 minutes at 300*xg*. The cells were then resuspended in 40 µL of either pre-complexed Siglec-Strep-Tactin or Siglec-liposomes and incubated for 30 minutes on ice. 150 µL of 1X PBS was then added to cells to stop the reaction. Cells were then centrifuged at 300*xg* for 5 minutes and washed with 1X PBS two times. Cells were then resuspended into 200 µL of flow buffer (PBS, 0.1% BSA, 0.1% EDTA) and proceed to flow cytometry. Cells were kept on ice for the duration of flow cytometry. At least 10,000 events were collected for the flow cytometry experiments. Flow cytometry data was analyzed using FlowJo (10.5.3) software (BD Biosciences). Cell lines used for Siglec-Fc staining in this paper are as followed: WT U937 cells for Siglec-1, -2, -6, -7, and -11, CHST1 overexpressing (OE) U937 cells(*14*) for Siglec-3, -5, -8, and -15, CHST2 OE U937 cells(*14*) for Siglec-9, CMAH OE U937 cells for Siglec-10, and WT CHO cells for Siglec-4.

### Staining of ex vivo expanded Tregs and Tconv cells

Siglec-liposomes were prepared as described above with a final concentration of Siglec-1, -2, -4, -7, and -9 of 8 µg/mL, and for the rest of the Siglecs, was 16 µg/mL. *Ex vivo* expanded Treg and Tconv cells were grown for 2 days in 37 °C incubator with 5% CO2. Cells were then seeded in a 96-well U-bottom microplate and centrifuged for 5 minutes at 300 *xg*. Siglec-liposomes were then added to the cells, and the binding to Tregs and Tconv cells was measured by flow cytometry as described above.

### Flow cytometry assay on primary cells isolated from human peripheral blood

Human blood was first aliquoted into 5 mL volumes in 50 mL falcon tubes and 45 mL of 1X ACK (Ammonium–chloride– potassium) lysis was added to cells and incubated at room temperature for 5 minutes to deplete red blood cells. Cells were then centrifuged for 5 minutes at 400*xg*, 4 °C. The pellets were then resuspended in 5 mL of 1X ACK buffer and incubated at room temperature for 2 minutes. Cells were then centrifuged for 5 minutes at 400*xg*, 4 °C and the pellets were resuspended in 1X ACK buffer and were centrifuged for 5 minutes at 400xg, 4 °C. The pellet was then resuspended in 1X PBS to neutralize the lysis buffer, and cells were centrifuged for 5 minutes at 400*×g*, 4°C. Cells were then seeded into 96-well U-bottom microplate and centrifuged for 5 minutes at 400*xg*, 4 °C. The cells were then resuspended in 40 µL of antibody cocktail (**Table S2**) and incubated on ice for 30 minutes. 150 µL 1X PBS were then added to the cells and the microplate was centrifuged for 5 minutes at 400*xg*, 4 °C. The cells were then washed 2 times with 150 µL of 1X PBS and centrifuged for 5 minutes at 400*xg*, 4 °C. Siglecs were conjugated to liposomes as described above, and the cell pellets were then resuspended in 40 µL of Siglec-liposomes, where the final concentration for Siglec-1, -2 and -7 was 8 µg/mL and for the rest of the Siglecs, was 16 µg/mL. Cells were then incubated on ice for 30 minutes. 150 µL of 1X PBS was then added to cells and cells were centrifuged and washed as described above. After, cells were resuspended into 200 µL of flow buffer (PBS, 0.1% BSA, 0.1% EDTA) and proceed to flow cytometry. Cells were kept on ice for the duration of flow cytometry. At least 500,000 events for each Siglec-liposomes were collected.

### Flow cytometry assay on Tregs and Tconv cells isolated from human peripheral blood

For staining Tregs and Tconv cells from PBMCs with Siglec-liposomes, all the steps described above were followed. After staining the cells with antibody cocktail (**Table S2**) and Siglec-liposomes, cells were fixed with 1% PFA (Thermo Fischer) on ice for 10 minutes. 200 µL of permeabilization solution (Biolegend) was added to cells and cells were centrifuged for 5 minutes at 450*xg*, and this step was repeated for three times. Anti-FOXP3 and anti-HEIOS antibodies **(Table S2)** were incubated with cells overnight and at 4 °C. The following day, cells were washed two times with the permeabilization solution and then 300 µL of flow buffer (PBS, 0.1% BSA, 0.1% EDTA) was added to the cells and analyzed using flow cytometry. 1,000,000 events for each sample were recorded.

### Flow cytometry assay on primary cells from mouse splenocytes

Spleens from mice were dissociated into single cells. Cells were then centrifuged for 5 minutes at 400*xg*, 4 °C. After, 1 mL of 1X ACK lysis buffer was added per spleen and the cells were incubated at room temperature for one minute. Cells were then centrifuged for 5 minutes at 400*xg*, 4 °C. Cells were then washed with 10 mL of 1X PBS to neutralize the lysis buffer and centrifuged for 5 minutes at 400*xg*, 4 °C. The pellet was then resuspended in 1X PBS, and the cells were plated into a 96-well U-bottom microplate and the microplate was centrifuged for 5 minutes at 400*xg*, 4 °C. Cells were resuspended in 40 µL of antibody cocktail (**Table S3**) and incubated on ice for 30 minutes. After 30 minutes, 150 µL of 1X PBS was added to the cells and the microplate was centrifuged for 5 minutes at 400*xg*, 4 °C. The cells were then washed with 150 µL of 1X PBS and centrifuged for 5 minutes at 400*xg*, 4 °C. Cells were resuspended into 40 µL of Siglec-2- and Siglec-7-liposomes in final concentration of 8 µg/mL. Cells were then incubated on ice for 30 minutes. 150 µL of 1X PBS was then added to cells and cells were centrifuged and washed as described above. Cells were resuspended into 200 µL of flow buffer (PBS, 0.1% BSA, 0.1% EDTA) and proceed to flow cytometry. Cells were kept on ice for the duration of flow cytometry. At least 500,000 events for each Siglec-liposomes were collected.

### Multiplexing experiments with Siglec-liposomes

To make multiplexed liposomes, WT Siglec-2 was conjugated to AF647-liposomes, R120A Siglec-2 was conjugated to AF488-liposomes and AF647- liposomes, WT Siglec-7 was conjugated to AF555-liposomes, R124A Siglec-7 was conjugated to AF488- liposomes and AF555-liposomes, WT Siglec-1 was conjugated to AF488-liposomes, and R116A Siglec-1 was conjugated to AF488-liposomes and AF555-liposomes into final concentration of 500 µM. Each Siglec-liposome was then run on a CL4B column (Cytiva) and fractions of 200 µL were collected. The presence of liposomes in each fraction was then validated by staining WT U937 cells and running flow cytometry as described above. The fractions containing liposomes were then mixed to make multiplexed liposomes. 40 µL of multiplexed liposomes was added to WT U937 cells and flow cytometry was performed as described above. At least 10,000 events were collected for each multiplexed liposome. For *in vivo* multiplexed liposomes, WT and R124A Siglec-7 were conjugated to AF488-liposomes and WT and R120A Siglec-2 were conjugated to AF647-liposomes. To create multiplexed liposomes, WT Siglec-7, WT Siglec-2 and R124A Siglec-7, R120A Siglec-2 were mixed with the final concentration of 1.3 mM for each Siglec-liposome.

### In vivo liposome injection and IF microscopy analysis

The Siglec-liposomes with appropriate concentrations for *in vivo* experiments were prepared as described above. Siglec-liposomes were injected into C57BL/6J mice intravenously using a 30 G needle. One hour after injection, mice were euthanized by CO_2_ inhalation and organs were collected. Spleens were homogenized and analyzed by the flow cytometry as previously described. For immunofluorescence (IF) analysis, organs (brain, spleen, kidney) were embedded in the Optimal-Cutting-Temperature (OCT) (Thermo Fischer, Catalog No. 23730571) compound and were snap-frozen by immersing into ice cold isopentane on dry ice. Using a cryostat, 10 µm sections were prepared on the slides. For co-staining of other cell markers, slides were fixed with 4 % paraformaldehyde (PFA) for 10-15 min after 1X PBS washing. Sections were then blocked with bovine serum albumin (BSA, 10 mg/ml in PBS) for 1 hour at room temperature. Then, the sections were stained with 1^st^ antibody overnight at 4°C, following staining with the 2^nd^ antibody for 2 hours at room temperature. For co-staining of vascular endothelial cells, αCD31 antibody (R&D systems, AF3628, 1:50, goat) and agoat AF488 antibody (Thermo Fisher, A11055, 1:400-500) were used. For co-staining CD45R and CD3e on spleen sections, aCD45R (B220, BioLegend Catalog No. 103225, 1:200) conjugated with AF488 and aCD3e antibody (BD Catalog No. 553238, 1:200) conjugated with AF568 (1:400) were used. All the steps were performed without any detergents (i.e. Triton-X) to avoid collapse of liposomes. The microscopy LSM 700 (Zeiss) was used for capturing images with 20x magnification. The images were analyzed and processed with Zen 2.6 Blue edition software (Zeiss).

### Immunohistochemistry for mice tissues with Siglec-Fc

One year old mice were anesthetized with isoflurane and perfused with 20 mL of 1X PBS (Thermo Fischer). Brain tissue blocks and 10 µm brain sections on slides were prepared as described above. Siglec-7-Fc in PBS was pre-complexed with a polyclonal human antibody (Invitrogen, Ref-A21445) by incubating on ice for 1 hour. Sections on the slides were fixed with 4 % paraformaldehyde (PFA) for 10 min. Sections were washed with 1X PBS, then treated with StcE. Tissues were incubated with either StcE (40 µg/mL) or 1X PBS for 2 hours at 37 °C. Following incubation slides were washed with 1X PBS. Sections were then blocked with bovine serum albumin (BSA, 10 mg/ml in PBS) at room temperature for one hour. For co-staining with CD31, αCD31 (AF3628, R&D systems, 1:50) antibody with agoat antibody conjugated AF488 (A11055, Thermo Scientific, 1:500) were used as described above. Siglec-Fc complexes were then added on the slides and incubated at room temperature for two hours. The LSM 700 microscope (Zeiss) was used for capturing images with 20 x magnification. The images were analyzed and processed with Zen 2.6 Blue edition software (Zeiss).

### Isolation of human T cells from human peripheral blood, activating, and co-culturing with Siglec-liposomes

Human T cells were isolated based on the manufacturer’s protocol using human Pan T cell isolation kit (Miltenyi Biotech, order no: 130-097-095). Isolated T cells were resuspended in pre-warmed 1X PBS and stained with 5 μM of cell trace violet (CTV) (Invitrogen, Catalog No. C34557) for 6 min at 37°C and in the dark. 5 mL of Pre-warmed RPMI media (10% (v/v) heat-inactivated fetal bovine albumin (Gibco) was added to the T cells. T cells were then incubated for two minutes at room temperature to remove unbound CTV. 1 mL of pre-warmed RPMI (10% (v/v) heat-inactivated fetal bovine albumin (Gibco), 100 Units/mL Penicillin (Gibco), 100 μg/mL Streptomycin (Gibco), 0.02% Beta-mercaptoethanol (βME) (Gibco), 1% HEPES (Gibco), 1% Sodium pyruvate (Gibco), 1% MEM non-essential amino acids (Gibco)) was then added to T cells. Number of T cells for each donor was measured, and human αCD3/CD28 dynabeads were added to cells (1:0.25 T cell: bead) (Thermo Fisher, Catalog No. 11161D). Siglec-1, -2, and -7 (8 µg/mL) and Siglec-3, -5, -9, and -10 (16 µg/mL) conjugated to liposomes (as described above) were added to a 96-well V bottom microplate. Following this, T cells were cultured at density of 50,000 cells/well in the above-mentioned media with 100 U/mL hIL-2 (BioLegend, Catalog No. 589104) for 96 hours. T cell proliferation was measured by CTV dilution and via flow cytometry using an antibody cocktail (**Table S2**). For co-culturing T cells with Siglec-7-Fc, 8 µg/mL of either WT or R124A Siglec-7 was co-cultured with T cells as described above.

### Isolation of WT mouse T cells from splenocytes and co-culturing with Siglec-7-liposomes

Mouse Spleens were dissociated into single cells, and red blood cells were lyzed as described above. White blood cells where then incubated with the biotin-Antibody cocktail (**Table S3**) for 45 minutes on ice. Cells were then washed with 5 mL of 1X PBS and incubated with 20 µL of anti-biotin microbeads per spleen (Milteny Biotec, order no: 130-090-485), and incubated on ice for 15 minutes. While incubating, LS column (Milteny Biotec, order no: 130-042-401) was washed using 3 mL of MACS buffer (1% BSA, 0.1% EDTA in PBS). Cells were then loaded into the LS column and the flow through containing T cells was collected. T cells were then labeled with CTV (Invitrogen, Catalog No. C34557) as described above and activated with mouse αCD3/CD28 dynabeads (1:1 T cell: bead). Siglec-7-liposomes with a final concentration of 8 µg/mL in both WT and R124A were added to a 96-well V bottom microplate. T cells were then seeded to each well with density of 50,000 cells/well in RPMI (10% (v/v) heat-inactivated fetal bovine albumin (Gibco), 100 Units/mL Penicillin (Gibco), 100 μg/mL Streptomycin (Gibco), 0.02% Beta-mercaptoethanol (βME) (Gibco), 1% HEPES (Gibco), 1% Sodium pyruvate (Gibco), 1% MEM non-essential amino acids (Gibco), and cultured for 72 or 96 hours. T cell proliferation was then measured by CTV dilution and via flow cytometry using an antibody cocktail (**Table S3**).

### Isolation of CD43^-/-^ mouse T cells from splenocytes and co-culturing with Siglec-7-liposomes

To test the contribution of CD43 glycoprotein on T cell proliferation towards Siglec-7-liposomes, we lethally irradiated wild type C57 mice (WT) and then reconstituted with the bone marrow collected either from CD43^-/-^ mice or from WT mice (as control group). Mice were euthanized after eight weeks those, spleen and various lymph nodes (Inguinal, axillary, ilial, renal & popliteal) were collected. T cells were then separated from splenocytes of CD43^-/-^ mice as described above. T cells were then labeled with CTV, activated with mouse αCD3/CD28 dynabeads (1:1 T cell: bead), and co-cultured with the Siglec-7-liposomes as described above. For T cell proliferation analysis, flow cytometry was used with an antibody cocktail (**Table S3**).

### Isolation of ST3Gal1^-/-^ mouse T cells from splenocytes and co-culturing with Siglec-7-liposomes

To study the role of ST3Gal1 in effect of Siglec-7-liposomes on T cell proliferation, we used ST3Gal1^-/-^ with suitable wild type control mice. After acclimatization, those mice were euthanized, and spleen and various lymph nodes (Inguinal, axillary, ilial, renal & popliteal) were collected. T cells were then separated from splenocytes of ST3Gal1 ^-/-^ mice as described above. T cells were then labeled with CTV, activated with αCD3/CD28 dynabeads (1:1 T cell: bead), and co-cultured with the Siglec-7-liposomes as described above. For T cell proliferation analysis, flow cytometry was used with an antibody cocktail (**Table S3**).

## Acknowledgments

We are grateful to Kayla Graham, for her help obtaining blood samples for this study. We thank Taylor E. Gray for providing the diCBM40 lectin.

## Author contributions

Conceptualization: Z. J-C., R. D., and M. S. M.

Methodology: Z. J-C., and M. S. M.

Investigation: Z. J-C., E. N. S., M. C., K. T-Y., G. M. L., L. L-D., J. J., C. D. S., J. R. E., S. S.

Visualization: Z. J-C., and M. S. M. Supervision: M. S. M.

Writing—original draft: Z. J-C., and M. S. M.

