## Supplementary Information for "An ultrasensitive and modular platform to detect Siglec ligands and control immune cell function"

**This PDF file includes:**

Figs. S1 to S35
Tables S1 to S3

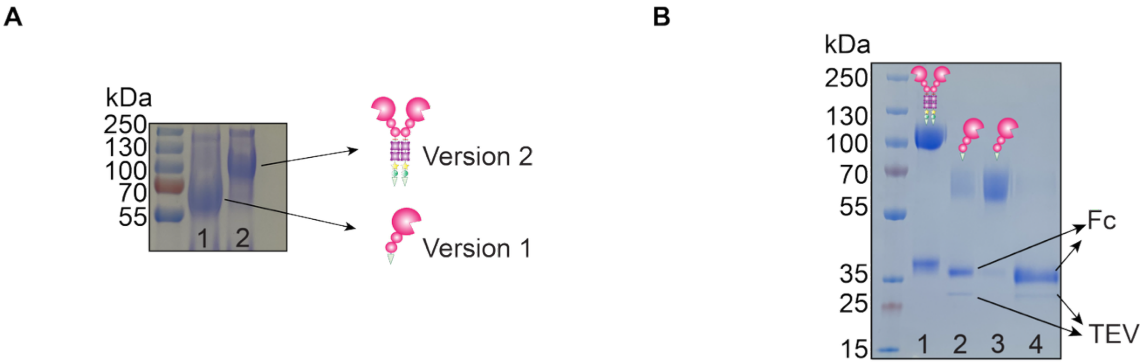


**Fig. S1. SDS-PAGE for the monomer of Version 1 (Siglec-7-ST-Fc) and Version 2 (Siglec-7-Fc-ST) of Siglec-7 and Version 1 treated with TEV protease.** (**A**) SDS-PAGE for the monomer (Siglec-7-ST) of Version 1 (Siglec-7-ST-Fc) in lane 1 and Version 2 (Siglec-7-Fc-ST) in lane 2. (**B**) SDS-PAGE of the steps for creating monomer of Version 1 (Siglec-7-ST). Lane 1: Siglec-ST-Fc. Lane 2: Siglec-ST-Fc treated with TEV enzyme. Lane 3: Siglec-7-ST after treatment with TEV and cleaned with $Ni^{2+}$ column. Lane 4: Elution from the $Ni^{2+}$ column.

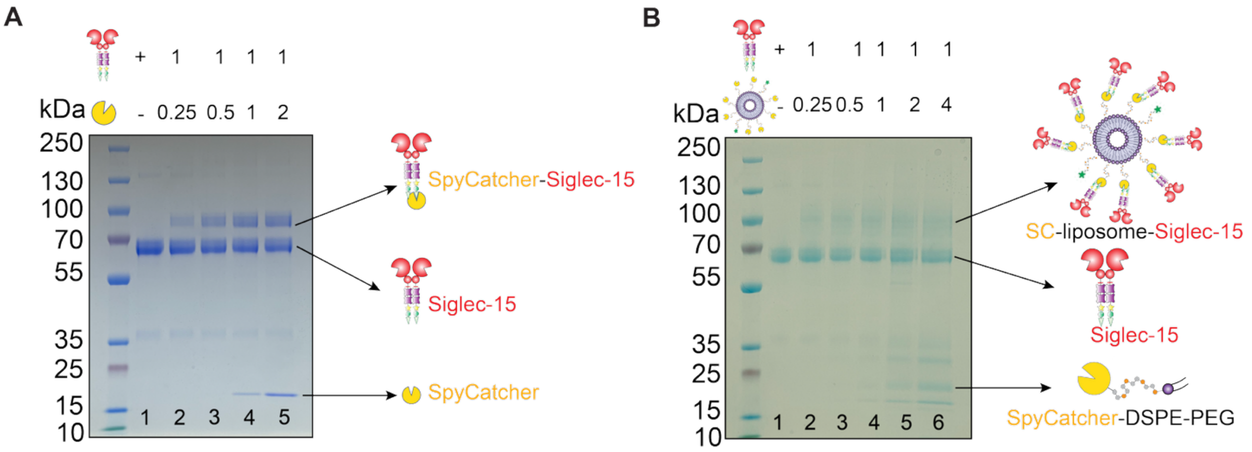


**Fig. S2. SDS-PAGE for Siglec conjugation to SC and SC-liposome.** (**A**) SDS-PAGE for conjugation of Siglec-15-Fc to SC protein in different molar ratios. Lane 1: Siglec-15-Fc. Lane 2: Siglec-15-Fc conjugated with SpyCatcher in 1:0.25 ratio (Siglec-15-Fc:SpyCatcher). Lane 3: Siglec-15-Fc conjugated with SpyCatcher in 1:0.5 ratio (Siglec-15-Fc:SpyCatcher). Lane 4: Siglec-15-Fc conjugated with SpyCatcher in 1:1 ratio (Siglec-15-Fc:SpyCatcher). Lane 5: Siglec-15-Fc conjugated with SpyCatcher in 1:2 ratio (Siglec-15-Fc:SpyCatcher). (**B**) SDS-PAGE for conjugating Siglec-15-Fc to SC-liposomes in different molar ratios. Lane 1: Siglec-15-Fc. Lane 2: Siglec-15-Fc conjugated with SC-liposome in 1:0.25 ratio (Siglec-15-Fc:SC-liposome). Lane 3: Siglec-15-Fc conjugated with SC-liposome in 1:0.5 ratio (Siglec-15-Fc:SC-liposome). Lane 4: Siglec-15-Fc conjugated with SC-liposomes in 1:1 ratio (Siglec-15-Fc:SC-liposome). Lane 5: Siglec-15-Fc conjugated with SC-liposome in 1:2 ratio (Siglec-15-Fc:SC-liposome). Lane 6: Siglec-15-Fc conjugated with SC-liposomes in 1:4 ratio (Siglec-15-Fc:SC-liposome).

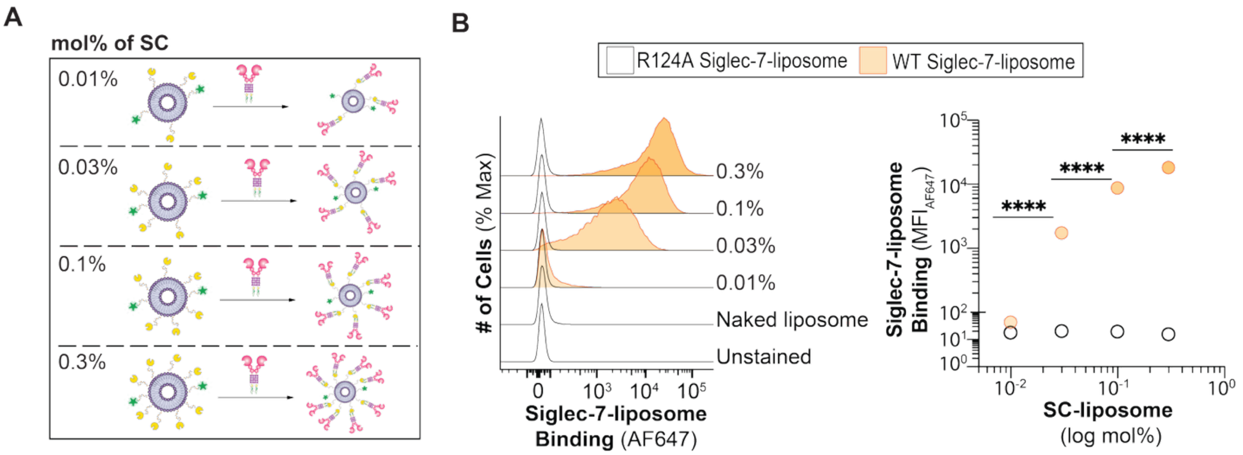


**Fig. S3. Titration of SC on liposome.** (**A**) Schematic of different densities of SC on liposome and conjugating to V2 Siglec-7. (**B**) Binding of WT and R124A V2 Siglec-7-liposomes to U937 cells in different densities of SC displayed on liposome. Data is presented as flow cytometry histograms (*left* panel) and median fluorescent intensity (MFI) of Siglec-7 binding (*right* panel). The *p* value for three technical replicates was calculated using an unpaired one-way ANOVA. ****$p$<0.0001.

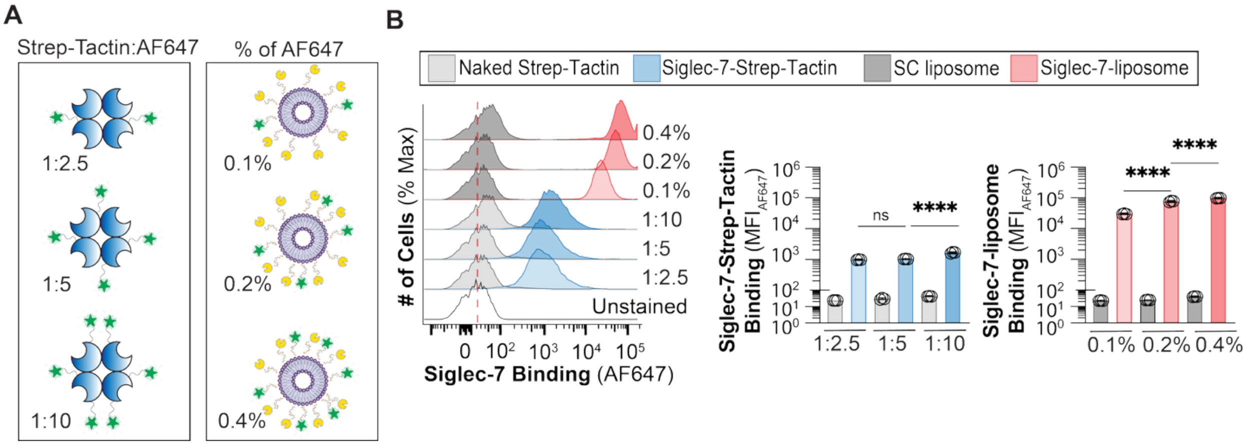


**Fig. S4. AF647 titration on Strep-Tactin and SC-liposomes.** (**A**) Schematic of AF647 titration on Strep-Tactin and 0.3 mol% SC-liposomes. (**B**) Binding of Siglec-7-Fc to U937 cells with different ratios of AF647 on Strep-Tactin and liposomes. Data is presented as flow cytometry histograms (*left* panel) and MFI of Siglec-7-Fc binding (*right* panels). The *p* value for three technical replicates was calculated using an unpaired one-way ANOVA. Not significant (ns) $p>0.05$, ****$p<0.0001$.

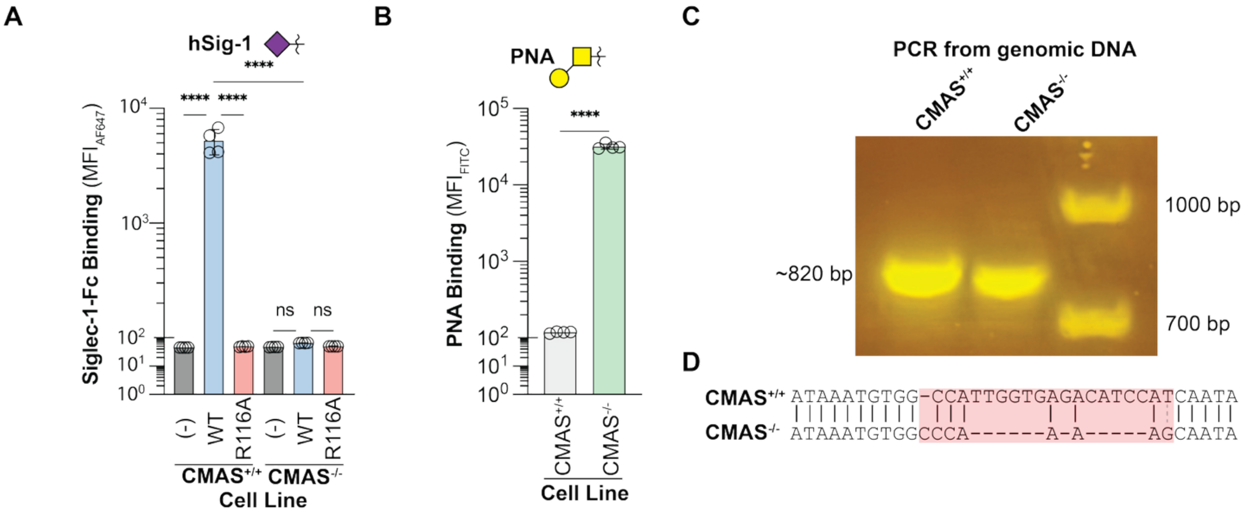


**Fig. S5. Validation of CMAS$^{-/-}$ CHO cells.** (**A**) Binding of WT and R116A Siglec-1-Fc to CMAS$^{+/+}$ and CMAS$^{-/-}$ CHO cells. Data is presented as MFI of Siglec-1 binding. (**B**) PNA binding to CMAS$^{+/+}$ and CMAS$^{-/-}$ CHO cells. Data is presented as MFI of PNA binding. (**C**) Agarose gel for PCR from genomic DNA of CMAS$^{+/+}$ and CMAS$^{-/-}$ CHO cells. (**D**) Sequencing result of PCR from genomic DNA of CMAS$^{+/+}$ and CMAS$^{-/-}$ CHO cells. The $p$ value for four technical replicates was calculated using an unpaired one-way ANOVA (a) and unpaired Student's $t$ test (b). ****$p$<0.0001.

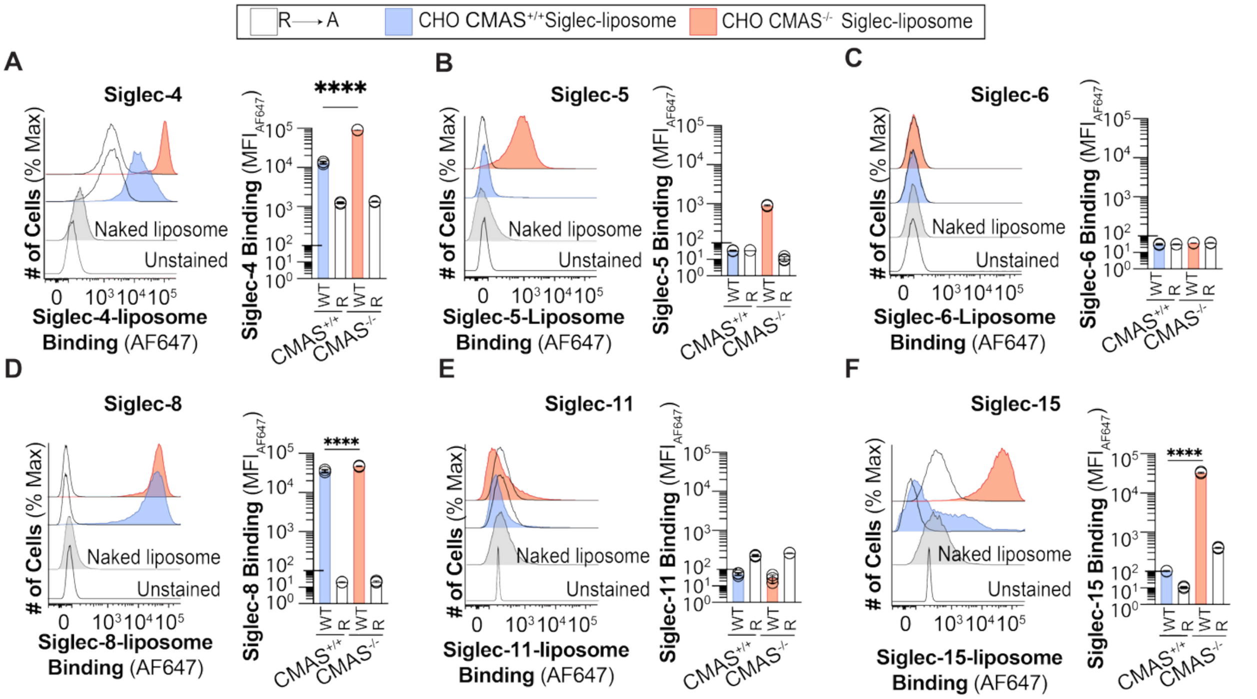


**Fig. S6. Binding of Binding of Siglec-4, -5, -6, -8, -11, and -15 expressed from CMAS$^{+/+}$ and CMAS$^{-/-}$ CHO cells.** Binding of 8 µg/mL of WT and arginine mutated of (**A**) Siglec-4-, (**B**) Siglec-5-, (**C**) Siglec-6-, (**D**) Siglec-8-, (**E**) Siglec-11-, and (**F**) Siglec-15-liposomes expressed from CMAS$^{+/+}$ and CMAS$^{-/-}$ CHO cells. Data is presented as flow cytometry histograms (*left* panel) and MFI of Siglec-liposomes binding (*right* panel). The *p* value for three technical replicates was calculated using an unpaired one-way ANOVA. ****$p$<0.0001.

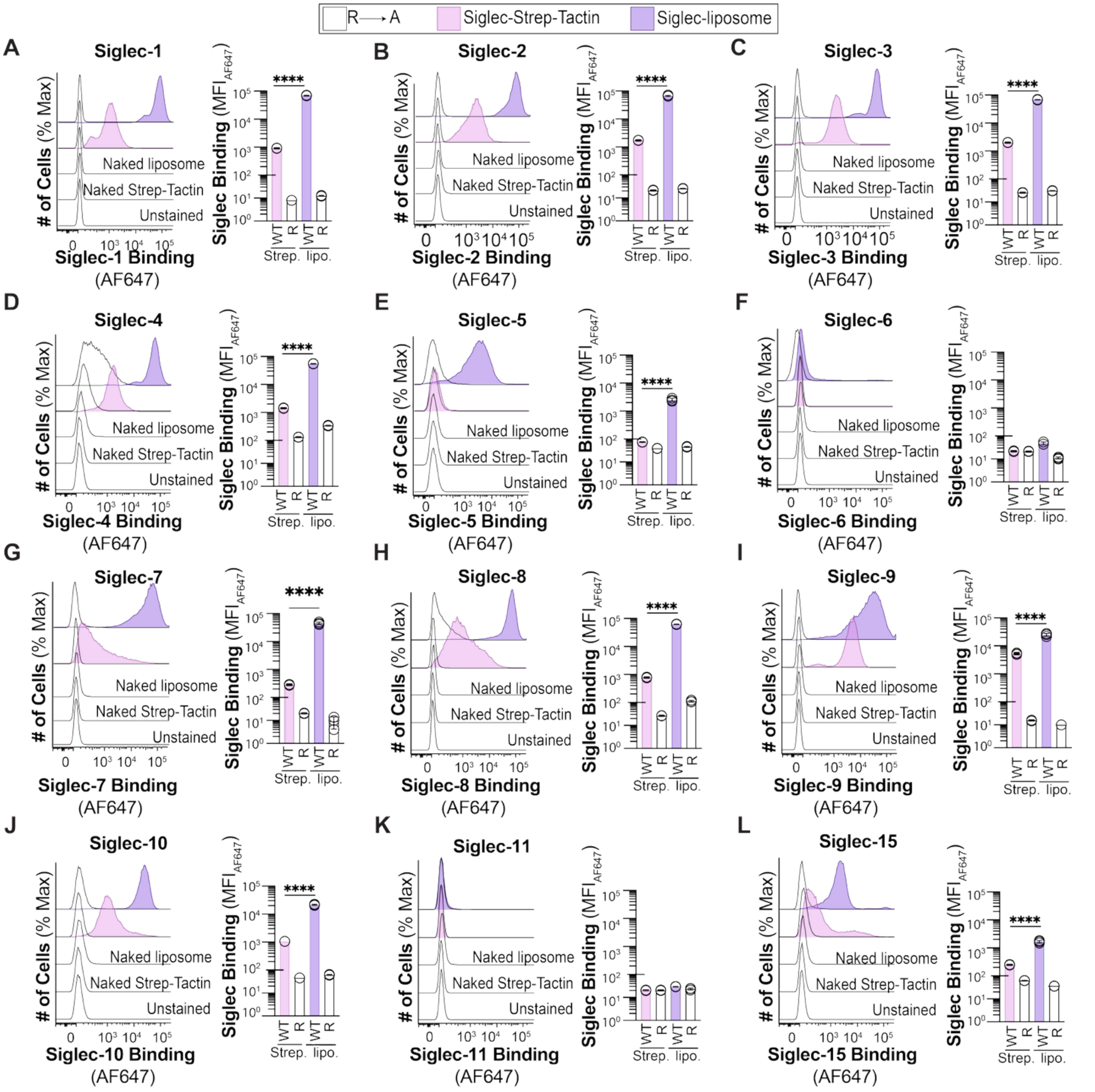


**Fig. S7. Binding of the human Siglec family expressed in CHO CMAS$^{-/-}$ cells and presented from liposome or Strep-Tactin platforms.** Binding of 8 μg/mL of WT and arginine mutated of (**A**) Siglec-1, (**B**) Siglec-2, (**C**) Siglec-3, (**D**) Siglec-4, (**E**) Siglec-5,(**F**) Siglec-6, (**G**) Siglec-7, (**H**) Siglec-8, (**I**) Siglec-9, (**J**) Siglec-10, (**K**) Siglec-11, and (**L**) Siglec-15 presented from liposome and Strep-Tactin. Data is presented as flow cytometry histograms and MFI of Siglec binding. The *p* value for three technical replicates was calculated using an unpaired one-way ANOVA. ****$p$<0.0001.

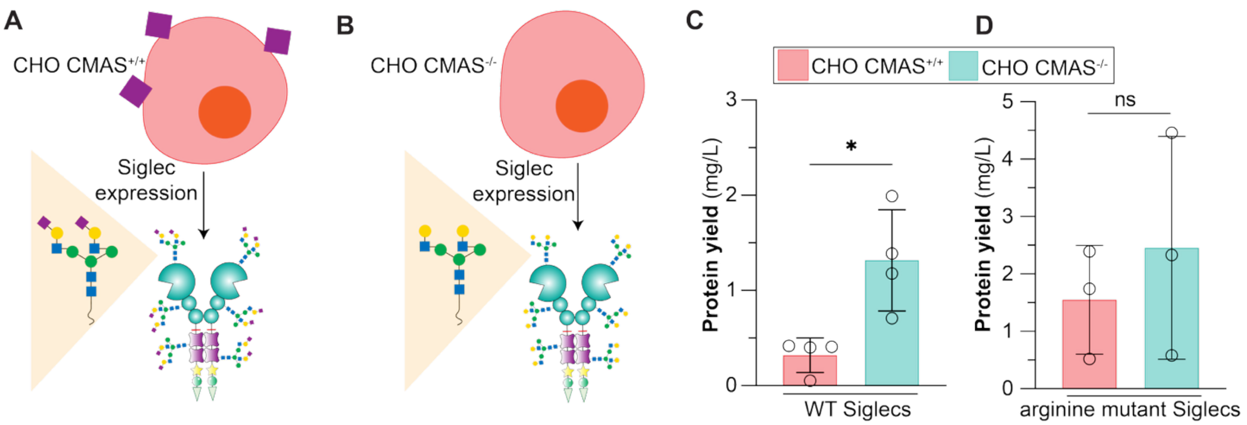


**Fig. S8. Siglec expression yields from CMAS$^{+/+}$ and CMAS$^{-/-}$ CHO cells.** (A) Schematic of Siglec expression from CMAS$^{+/+}$ CHO cells. (**B**) Schematic of Siglec expression from CMAS$^{-/-}$ CHO cells. (**C**) Protein expression yield of WT Siglecs in CMAS$^{+/+}$ CHO vs CMAS$^{-/-}$ CHO cells. (**D**) Protein expression yield of arginine mutated Siglecs from CMAS$^{+/+}$ CHO and CMAS$^{-/-}$ CHO cells. Siglecs used for expression comparison were Siglec-3, Siglec-5, Siglec-7, and Siglec-9. The *p* value for four (panel c) and three (panel d) technical replicates was calculated using an unpaired Student's *t* test. Not significant (ns) $p>0.05$, * $0.05>p\geq0.01$.

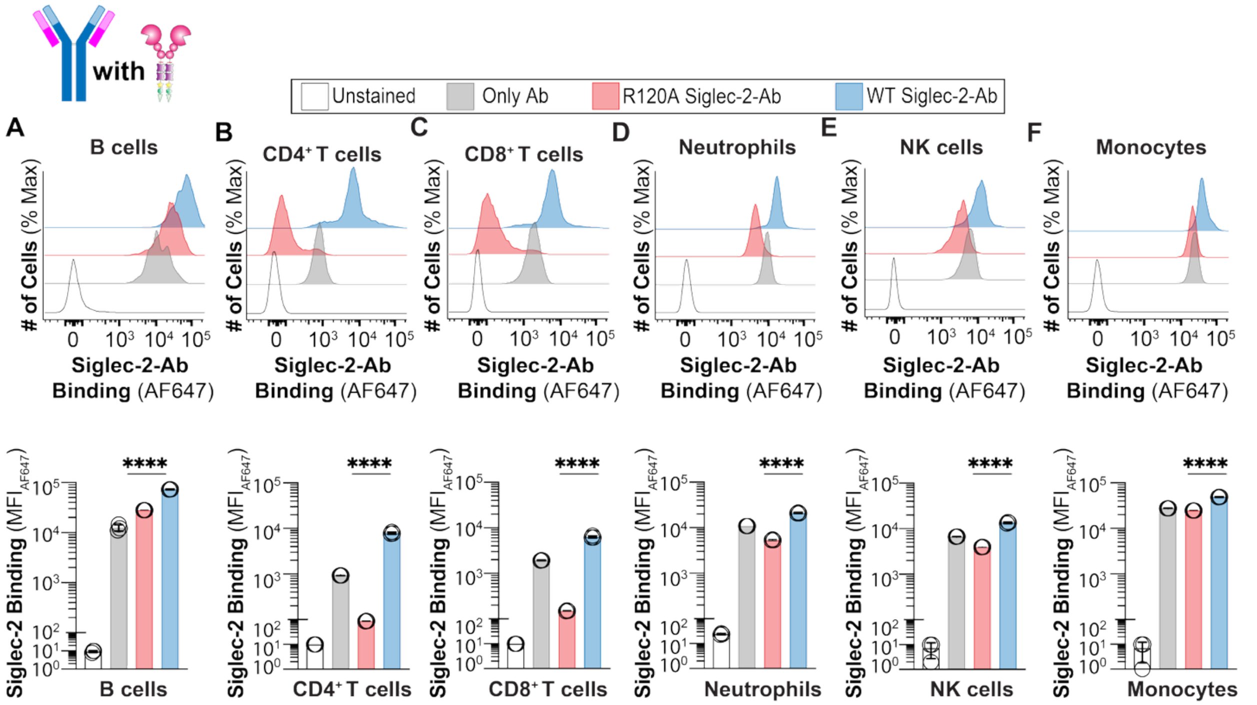


**Fig. S9. Binding of Siglec-2-Fc pre-complexed with secondary αhuman IgG1 antibody to immune cells isolated from human peripheral blood.** Binding of WT and R120A Siglec-2-Fc pre-complexed with αhuman IgG to (**A**) B cells, (**B**) $CD4^+$ T cells, (**C**) $CD8^+$ T cells, (**D**) Neutrophils, (**E**) NK cells, and (**F**) Monocytes isolated from human blood. Data is presented as flow cytometry representative histograms (*upper* panels) and MFI of Siglec-2-Fc binding (*lower* panels). The *p* value for three technical replicates was calculated using an unpaired one-way ANOVA. ****$p<0.0001$.

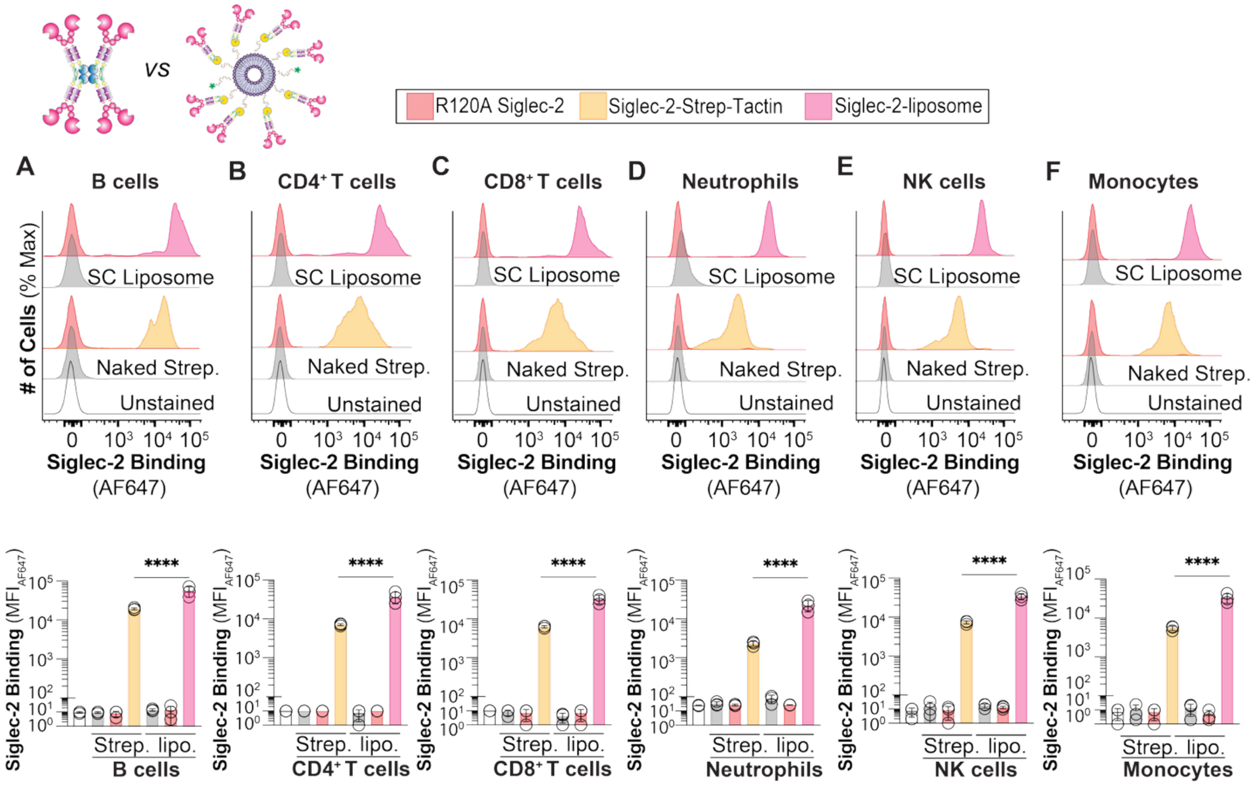


**Fig. S10. Binding of Siglec-2 in liposome and pre-complexed with Strep-Tactin to immune cells isolated from peripheral human blood.** Binding of WT and R120A Siglec-2 in liposomes and pre-complexed with Strep-Tactin to (**A**) B cells, (**B**) CD4$^+$ T cells, (**C**) CD8$^+$ T cells, (**D**) Neutrophils, (**E**) NK cells, and (**F**) Monocytes isolated from human blood. Data is presented as flow cytometry representative histograms (*upper* panels) and MFI of Siglec-2 binding (*lower* panels). The *p* value for three technical replicates was calculated using an unpaired one-way ANOVA. ****$p$<0.0001.

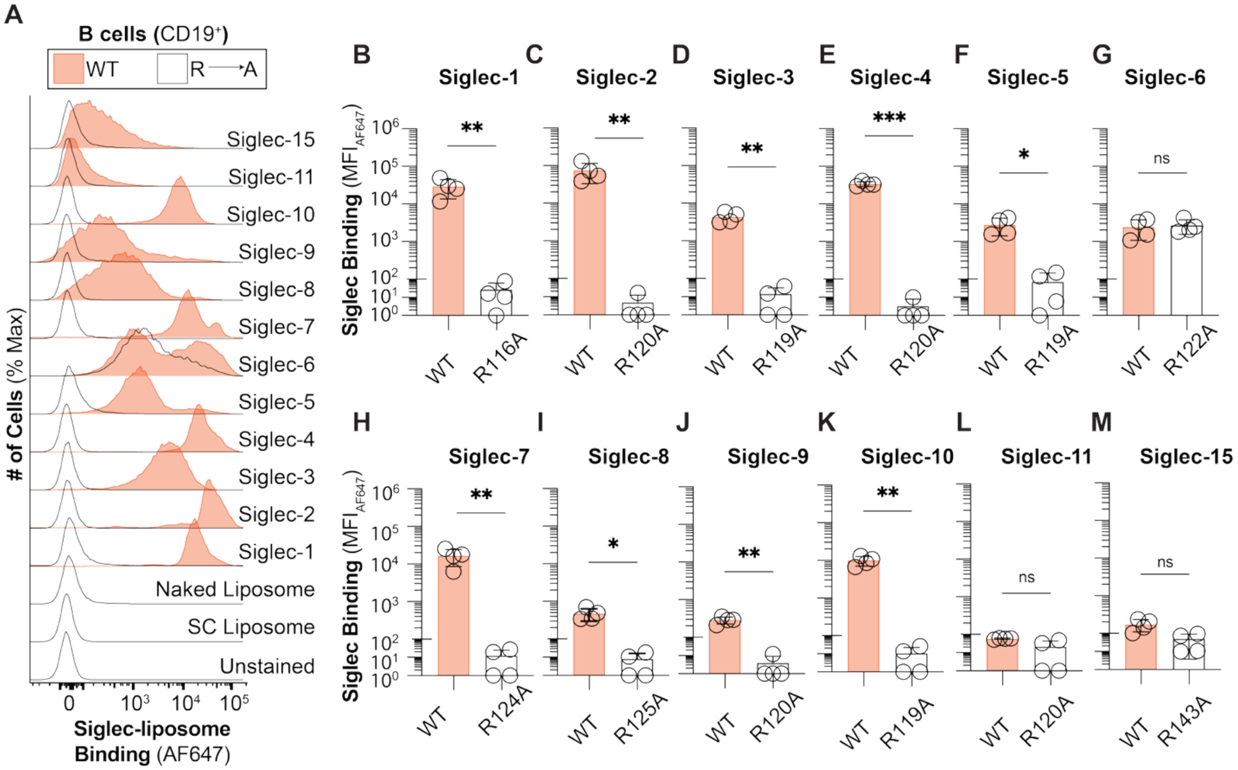


**Fig. S11. Binding of human Siglec family in liposome to B cells from peripheral human blood.** (**A**) Flow cytometry representative histograms of WT and arginine mutated of Siglec-liposomes binding to B cells isolated from human blood. MFI of (**B**) WT and R116A Siglec-1- (**C**) WT and R120A Siglec-2, (**D**) WT and R119A Siglec-3-, (**E**) WT and R118A Siglec-4-, (**F**) WT and R119A Siglec-5-, (**G**) WT and R122A Siglec-6-, (**H**) WT and R124A Siglec-7-, (**I**) WT and R125A Siglec-8-, (**J**) WT and R120A Siglec-9-, (**K**) WT and R119A Siglec-10-, (**L**) WT and R120A Siglec-11-, and (**M**) WT and R143A Siglec-15-liposomes to B cells. The $p$ value for four biological replicates was calculated using a paired one-way ANOVA. Not significant (ns) $p>0.05$, $*0.05>p\geq0.01$, $**0.01>p\geq0.001$, $***0.001>p\geq0.0001$.

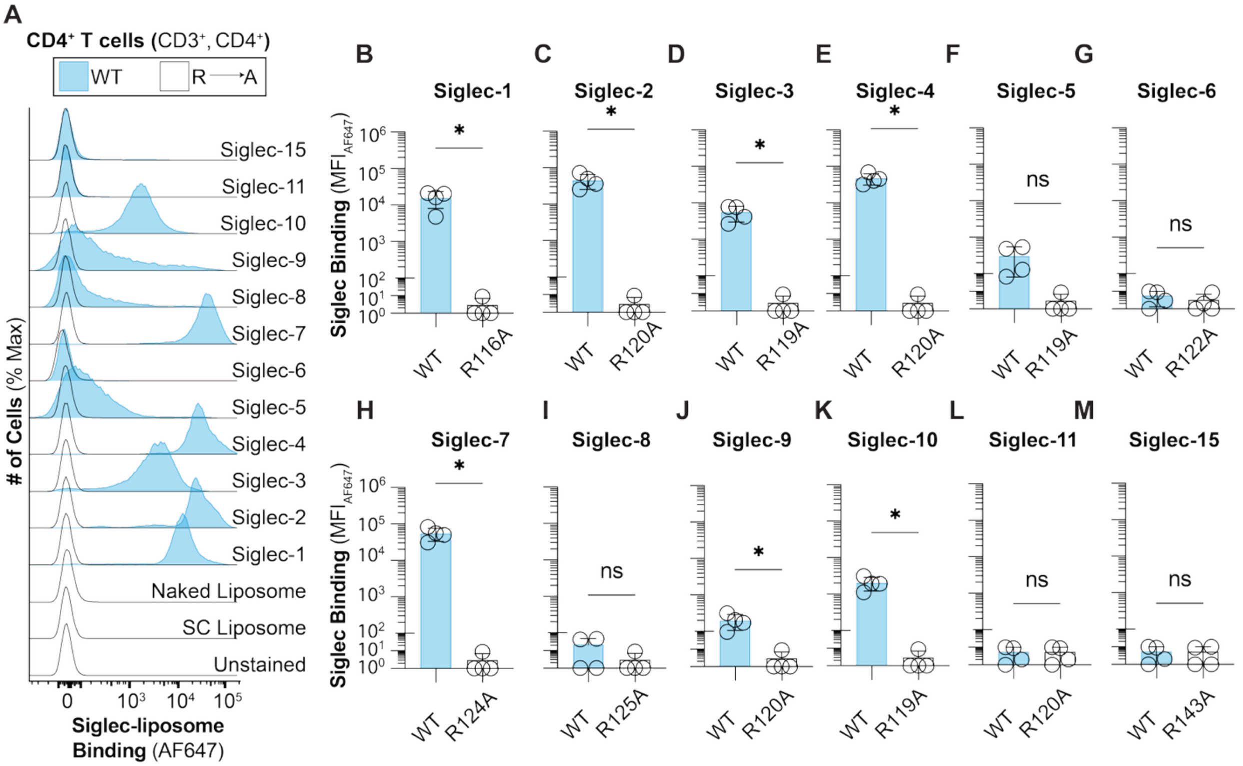


**Fig. S12. Binding of human Siglec family in liposome to CD4⁺ T cells from peripheral human blood.** (**A**) Flow cytometry representative histograms of WT and arginine mutated of Siglec-liposomes binding to CD4⁺ T cells isolated from human blood. MFI of (**B**) WT and R116A Siglec-1- (**C**) WT and R120A Siglec-2-, (**D**) WT and R119A Siglec-3-, (**E**) WT and R118A Siglec-4-, (**F**) WT and R119A Siglec-5-, (**G**) WT and R122A Siglec-6-, (**H**) WT and R124A Siglec-7-, (**I**) WT and R125A Siglec-8-, (**J**) WT and R120A Siglec-9-, (**K**) WT and R119A Siglec-10-, (**L**) WT and R120A Siglec-11-, and (**M**) WT and R143A Siglec-15-liposomes to CD4⁺ T cells. The $p$ value for four biological replicates was calculated using a paired one-way ANOVA. Not significant (ns) $p>0.05$, $*0.05>p\geq 0.01$.

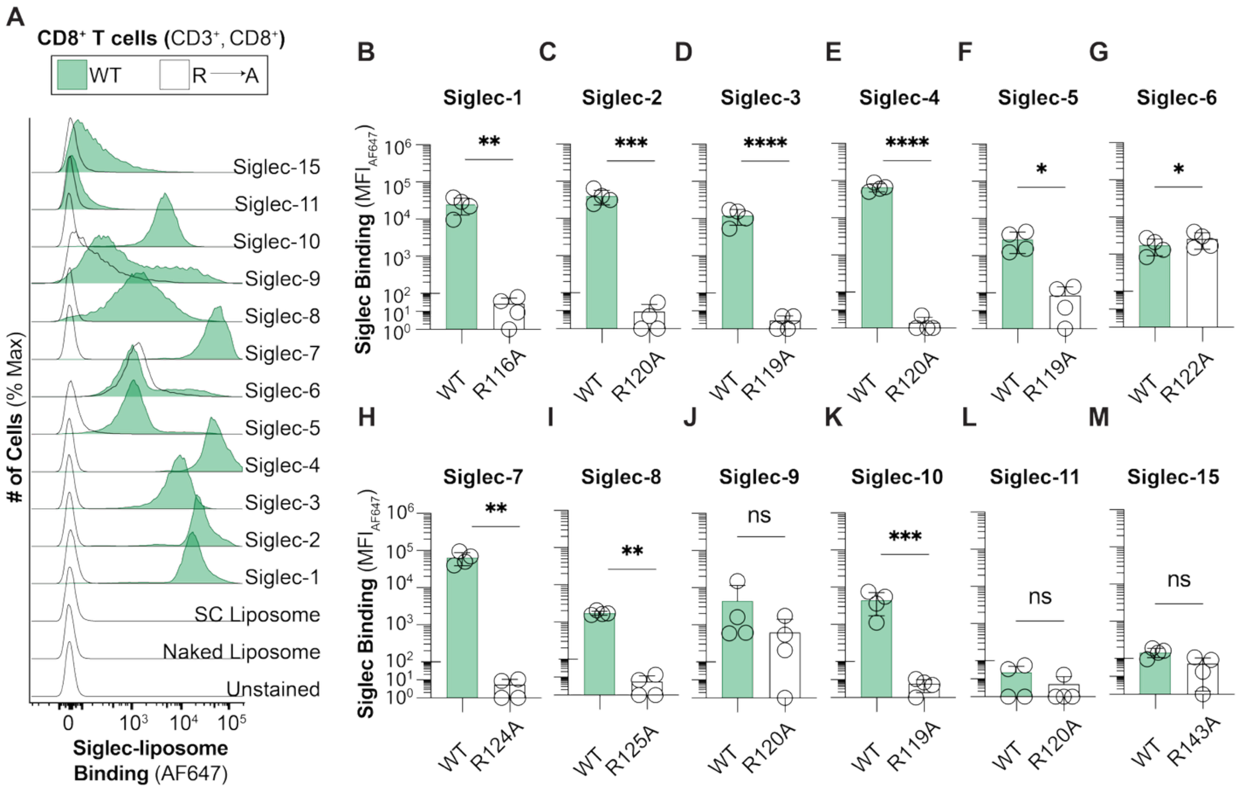


**Fig. S13. Binding of human Siglec family in liposome to CD8+ T cells from peripheral human blood.** (**A**) Flow cytometry representative histograms of WT and arginine mutated of Siglec-liposomes binding to CD8+ T cells isolated from human blood. MFI of (**B**) WT and R116A Siglec-1 (**C**) WT and R120A Siglec-2-, (**D**) WT and R119A Siglec-3-, (**E**) WT and R118A Siglec-4-, (**F**) WT and R119A Siglec-5-, (**G**) WT and R122A Siglec-6-, (**H**) WT and R124A Siglec-7-, (**I**) WT and R125A Siglec-8-, (**J**) WT and R120A Siglec-9-, (**K**) WT and R119A Siglec-10-, (**L**) WT and R120A Siglec-11-, and (**M**) WT and R143A Siglec-15-liposomes to CD8+ T cells. The *p* value for four biological replicates was calculated using a paired one-way ANOVA. Not significant (ns) $p>0.05$, $*0.05>p\geq0.01$, $**0.01>p\geq0.001$, $***0.001>p\geq0.0001$, $****p<0.0001$.

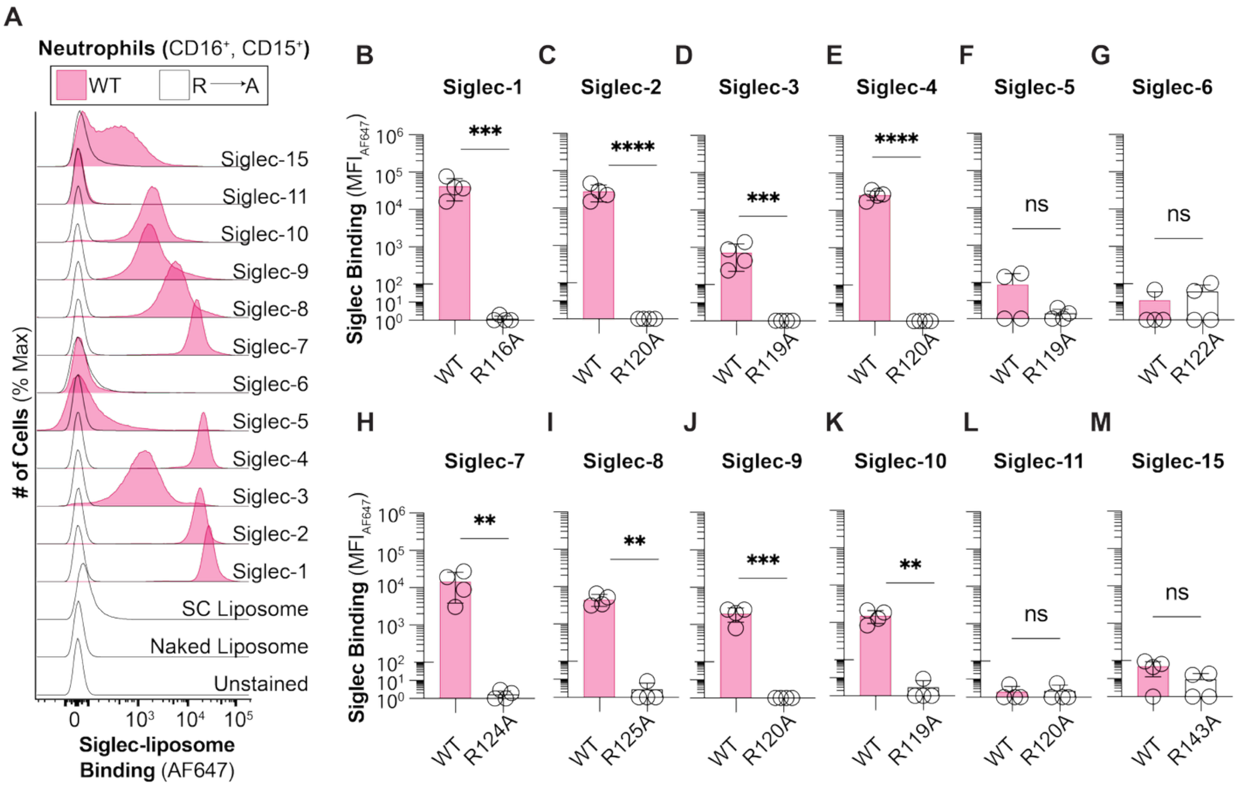


**Fig. S14. Binding of human Siglec family in liposome to neutrophils from peripheral human blood.** (**A**) Flow cytometry representative histograms of WT and arginine mutated of Siglec-liposomes binding to neutrophils isolated from human blood. MFI of (**B**) WT and R116A Siglec-1- (**C**) WT and R120A Siglec-2-, (**D**) WT and R119A Siglec-3-, (**E**) WT and R118A Siglec-4-, (**F**) WT and R119A Siglec-5-, (**G**) WT and R122A Siglec-6-, (**H**) WT and R124A Siglec-7-, (**I**) WT and R125A Siglec-8-, (**J**) WT and R120A Siglec-9-, (**K**) WT and R119A Siglec-10-, (**L**) WT and R120A Siglec-11-, and (**M**) WT and R143A Siglec-15-liposomes to neutrophils. The *p* value for four biological replicates was calculated using a paired one-way ANOVA. Not significant (ns) $p>0.05$, $^{**}0.01>p\geq0.001$, $^{***}0.001>p\geq0.0001$, $^{****}p<0.0001$.

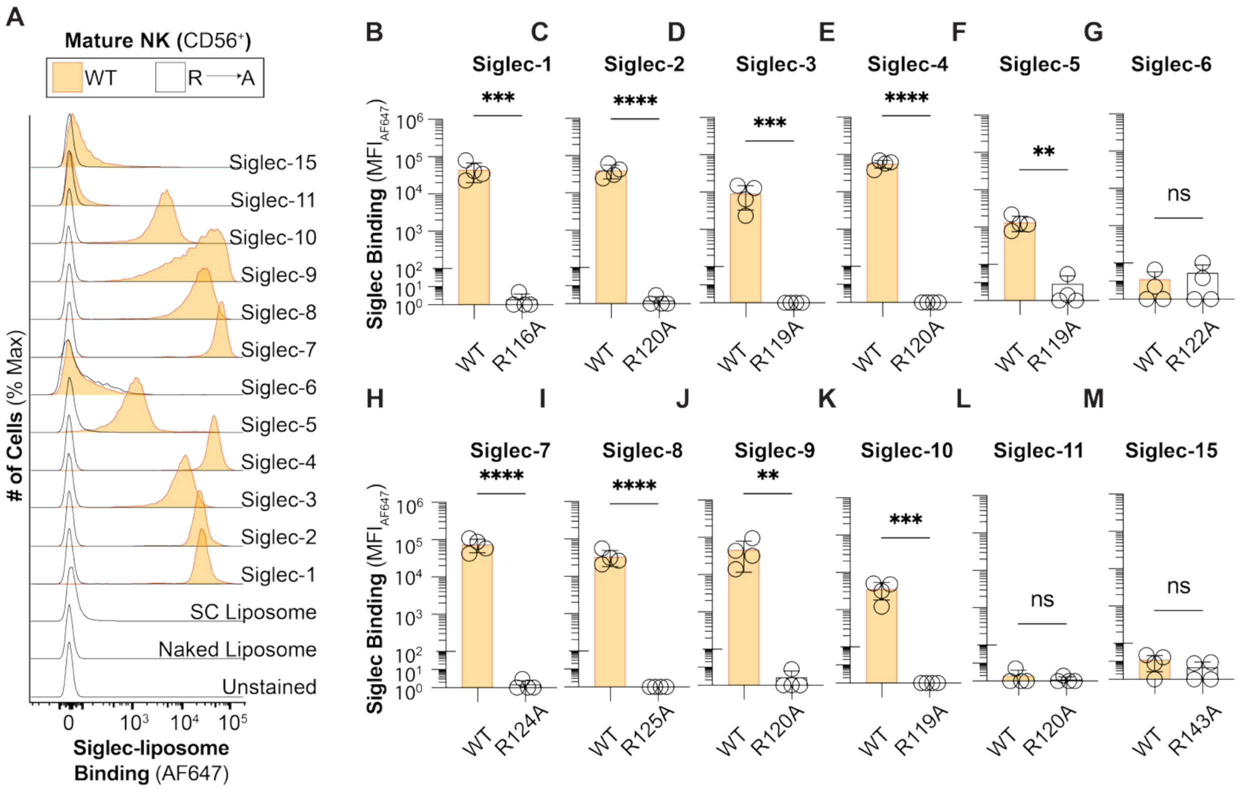


**Fig. S15. Binding of human Siglec family in liposome to mature NK cells from peripheral human blood.** (**A**) Flow cytometry representative histograms of WT and arginine mutated of Siglec-liposomes binding to mature NK cells isolated from human blood. MFI of (**B**) WT and R116A Siglec-1- (**C**) WT and R120A Siglec-2-, (**D**) WT and R119A Siglec-3-, (**E**) WT and R118A Siglec-4-, (**F**) WT and R119A Siglec-5-, (**G**) WT and R122A Siglec-6-, (**H**) WT and R124A Siglec-7-, (**I**) WT and R125A Siglec-8-, (**J**) WT and R120A Siglec-9-, (**K**) WT and R119A Siglec-10-, (**L**) WT and R120A Siglec-11-, and (**M**) WT and R143A Siglec-15-liposomes to mature NK cells. The $p$ value for four biological replicates was calculated using a paired one-way ANOVA. Not significant (ns) $p>0.05$, **$0.01>p\geq0.001$, ***$0.001>p\geq0.0001$, ****$p<0.0001$.

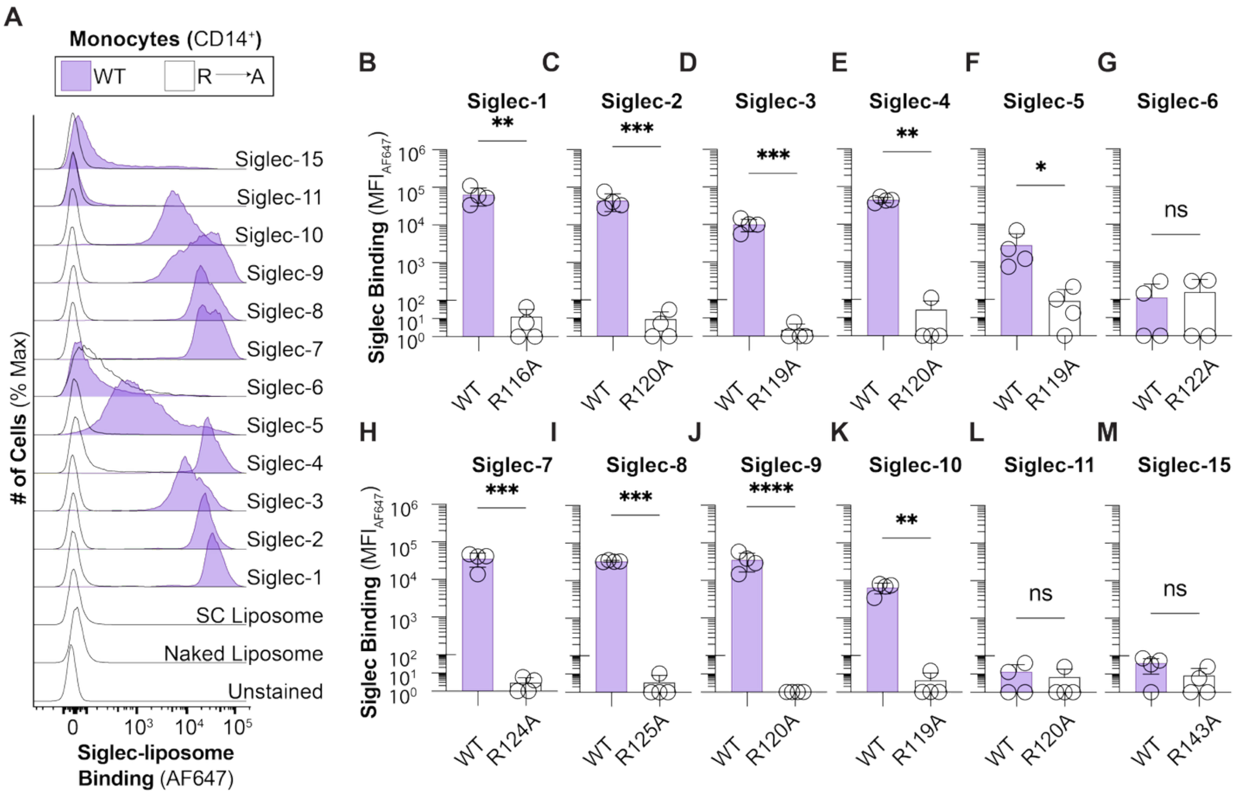


**Fig. S16. Binding of human Siglec family in liposome to monocytes from peripheral human blood.** (**A**) Flow cytometry representative histograms of WT and arginine mutated of Siglec-liposomes binding to monocytes isolated from human blood. MFI of (**B**) WT and R116A Siglec-1- (**C**) WT and R120A Siglec-2-, (**D**) WT and R119A Siglec-3-, (**E**) WT and R118A Siglec-4-, (**F**) WT and R119A Siglec-5-, (**G**) WT and R122A Siglec-6-, (**H**) WT and R124A Siglec-7-, (**I**) WT and R125A Siglec-8-, (**J**) WT and R120A Siglec-9-, (**K**) WT and R119A Siglec-10-, (**L**) WT and R120A Siglec-11-, and (**M**) WT and R143A Siglec-15- to monocytes. The $p$ value for four biological replicates was calculated using a paired one-way ANOVA. Not significant (ns) $p>0.05$, *$0.05>p\geq0.01$, **$0.01>p\geq0.001$, ***$0.001>p\geq0.0001$, ****$p<0.0001$.

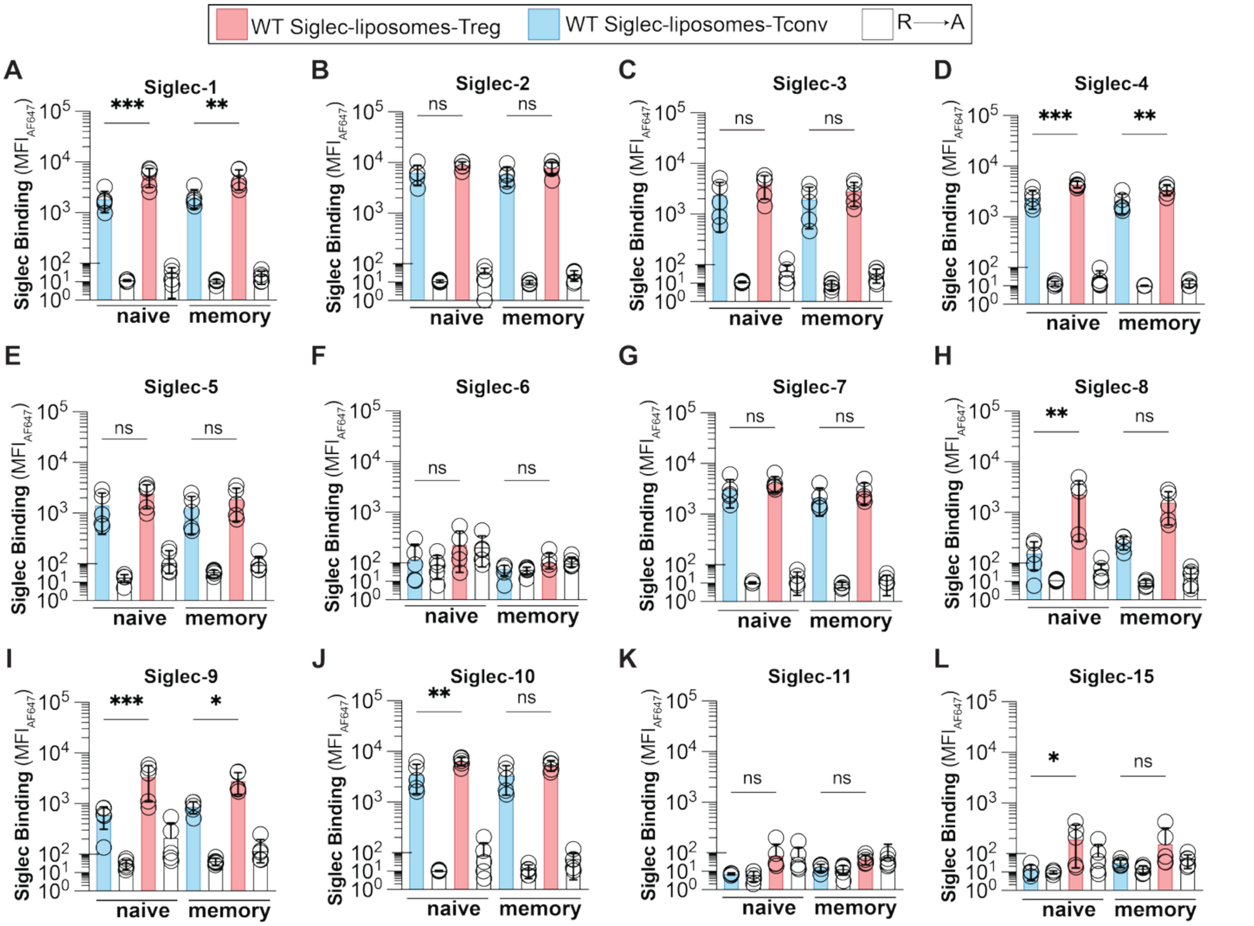


**Fig. S17. Binding of human Siglec family to naïve and memory Treg and Tconv CD4⁺ T cells isolated from peripheral human blood.** (**A-L**) Binding of WT and arginine mutated Siglec-liposomes to naïve and memory Treg and Tconv cells isolated from peripheral human blood. Data is presented as MFI of Siglec binding. The $p$ value for five biological replicates was calculated using a paired one-way ANOVA. Not significant (ns) $p>0.05$, *$0.05>p\geq 0.01$, **$0.01>p\geq 0.001$, ***$0.001>p\geq 0.0001$.

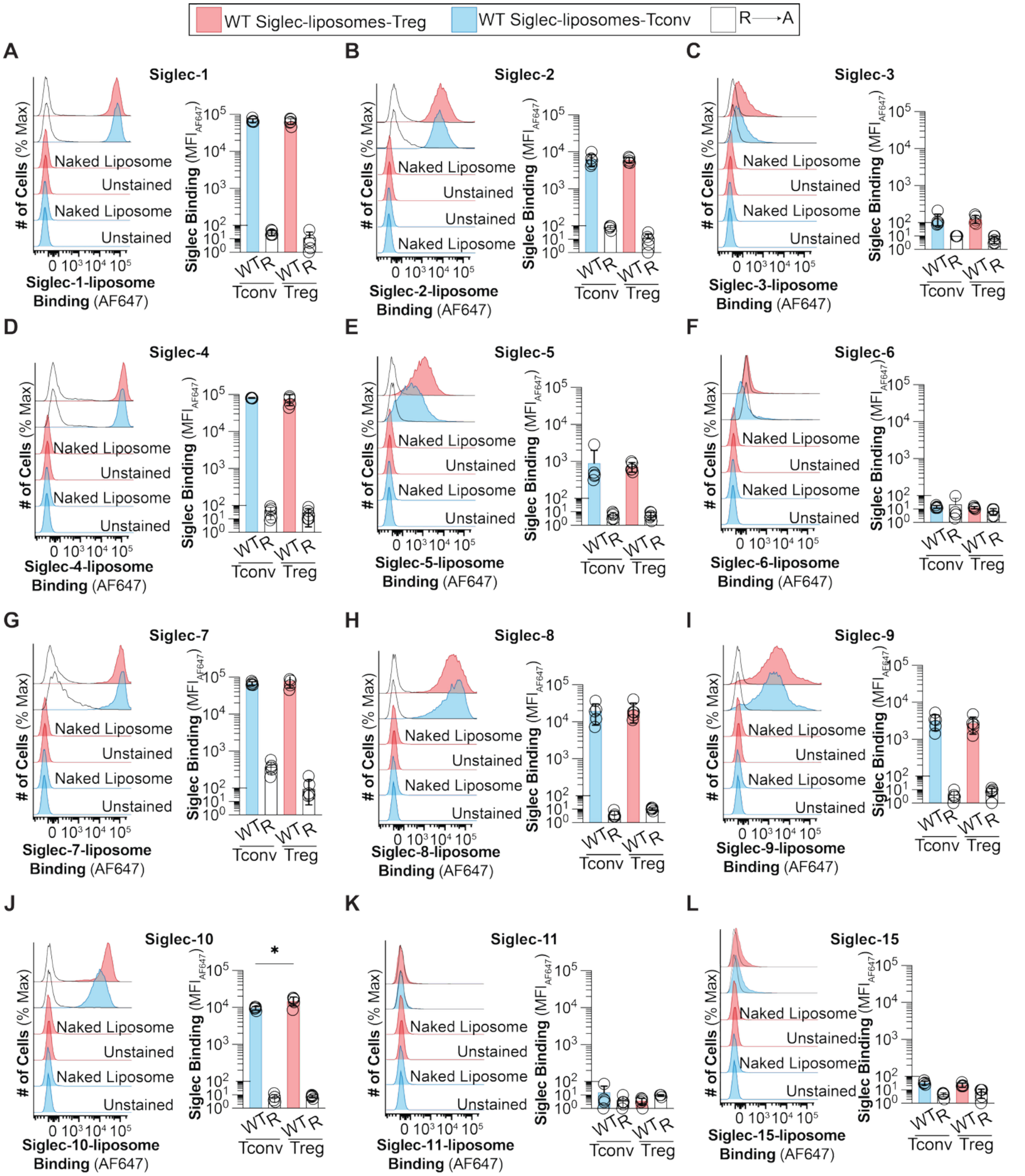


**Fig. S18. Binding of human Siglec family to *ex vivo* expanded Treg and Tconv CD4⁺ T cells.** (A-L) Binding of WT and arginine mutated Siglec-liposomes to *ex vivo* expanded Treg and Tconv cells. Data is presented as representative flow cytometry histograms and MFI of Siglec binding. The $p$ value for five biological replicates was calculated using a paired one-way ANOVA. *0.05>$p$≥0.01.

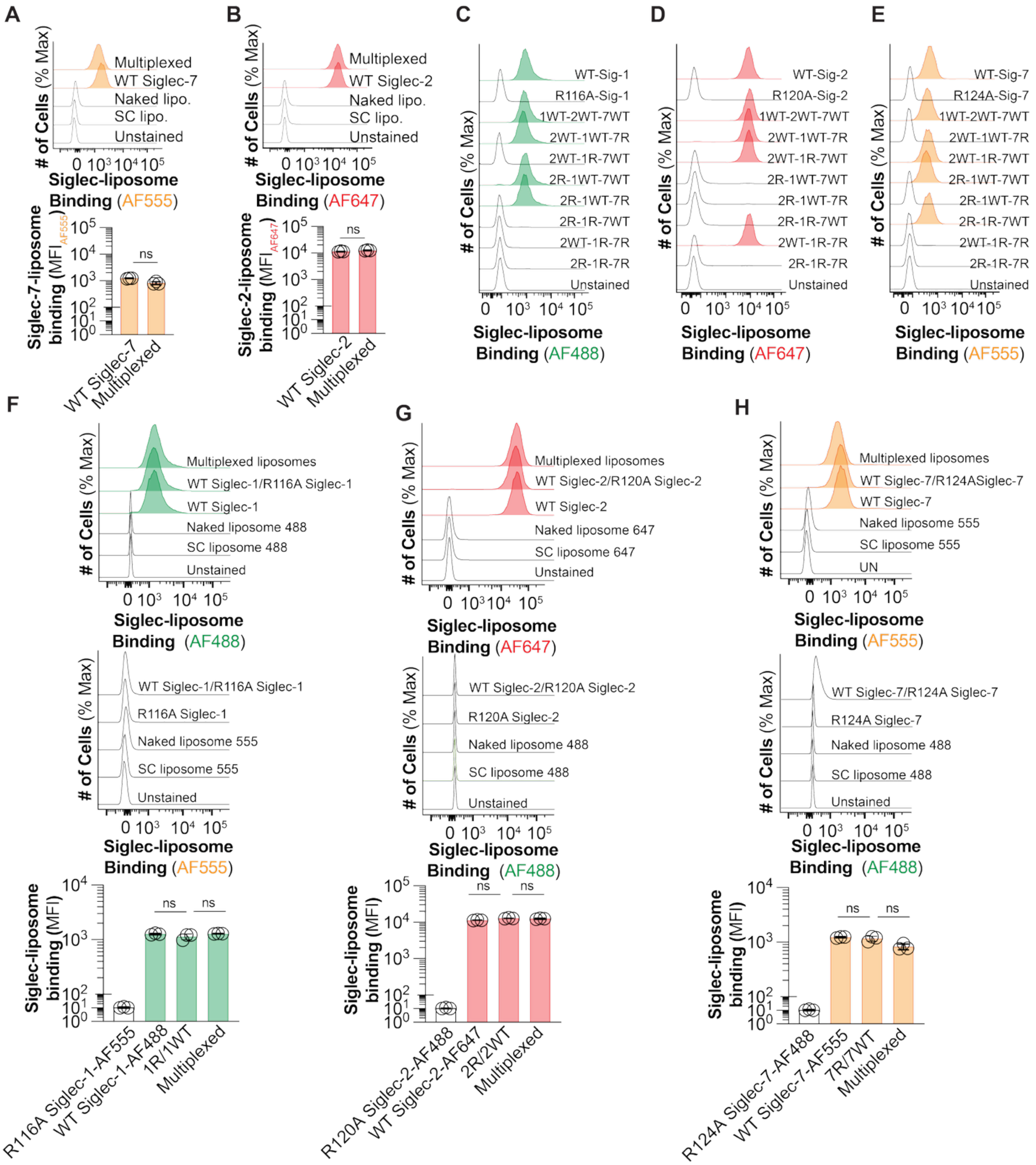


**Fig. S19. Binding of multiplexed Siglec-1-, Siglec-2-, and Siglec-7-liposomes to U937 cells.** Binding of multiplexed and single color of (**A**) AF55-Siglec-7-, and (**B**) AF647-Siglec-2-liposomes to U937 cells. Data is presented as flow cytometry histograms (*upper* panel) and MFI of Siglec binding (*lower* panel). Binding of multiplexed liposomes of (**C**) AF488-Siglec-1-, (**D**) AF647-Siglec-2-, and (**E**) AF555-Siglec-7-liposomes in different combinations to U937 cells. Data is presented as flow cytometry histograms. Binding of multiplexed WT and arginine mutated of (**F**) AF488-WT Siglec-1- and AF555-R116A Siglec-1-, (**G**) AF647-WT Siglec-2- and AF488-R120A Siglec-2-, and (**H**) AF555-WT Siglec-7- and AF488-R124A Siglec-7-liposomes. Data is presented as flow cytometry histograms (*upper* panels) and MFI of Siglec binding (*lower* panels). The *p* value for three technical replicates was calculated using an unpaired Student's *t* test (panel A, B) and an unpaired one-way ANOVA (panel F-H). Not significant (ns) $p>0.05$.

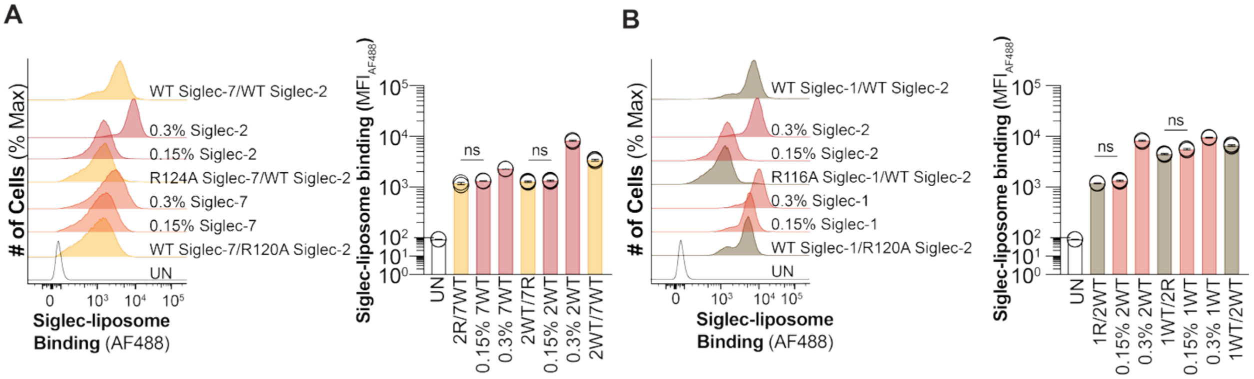


**Fig. S20. Binding of multiplexed liposomes with Siglec-2/Siglec-7 and Siglec-1/Siglec-2 on one liposome to U937 cells.** Binding of (**A**) Siglec-2 and Siglec-7 in 0.15 mol% and 0.3 mol% of SC-liposomes and multiplexed Siglec-2/Siglec-7-liposomes, and (**B**) Siglec-1 and Siglec-2 in 0.15 mol% and 0.3 mol% of SC-liposomes and multiplexed Siglec-1/Siglec-2-liposomes. Data is presented as flow cytometry histograms (*left* panel) and MFI of Siglec binding (*right* panel). The $p$ value for three technical replicates was calculated using an unpaired one-way ANOVA. Not significant (ns) $p>0.05$.

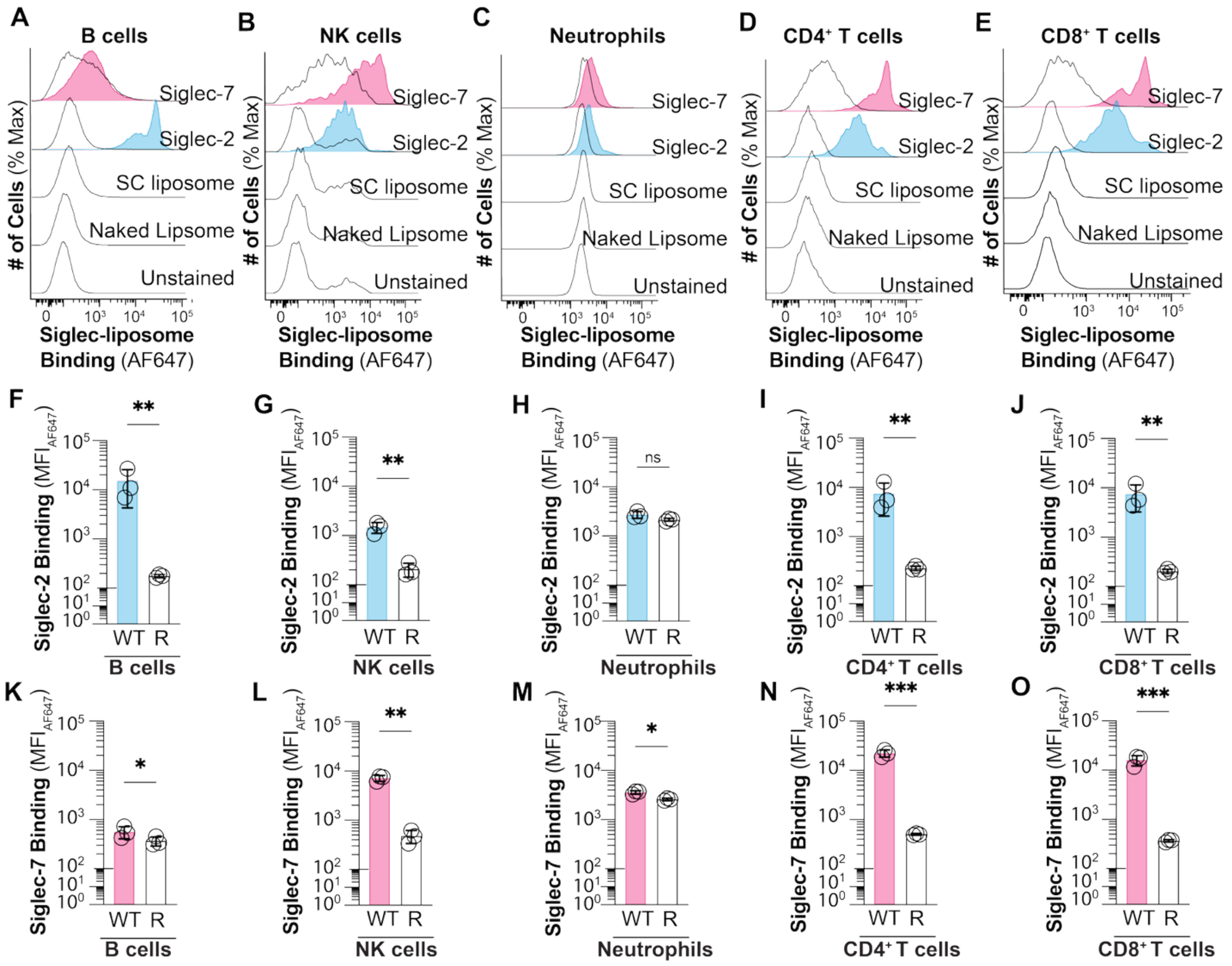


**Fig. S21.** ***In vitro*** **binding of Siglec-2-liposomes and Siglec-7-liposomes to immune cells isolated from mouse splenocytes.** Binding of WT and arginine mutated Siglec-2- and Siglec-7-liposomes to (**A**) B cells, (**B**) NK cells, (**C**) Neutrophils, (**D**) $CD4^+$ T cells, and (**E**) $CD8^+$ T cells. Data is presented as flow cytometry histograms. Binding of WT and R120A Siglec-2-liposomes to (**F**) B cells, (**G**) NK cells, (**H**) Neutrophils, (**I**) $CD4^+$ T cells, and (**J**) $CD8^+$ T cells. Data is presented as MFI of Siglec-2 binding. Binding of WT and R124A Siglec-7-liposomes to (**K**) B cells, (**L**) NK cells, (**M**) Neutrophils, (**N**) $CD4^+$ T cells, and (**O**) $CD8^+$ T cells. Data is presented as MFI of Siglec-7 binding. The $p$ value for four biological replicates was calculated using a paired Student's $t$ test. *$0.05>p\geq 0.01$, **$0.01>p\geq 0.001$, ***$0.001>p\geq 0.0001$.

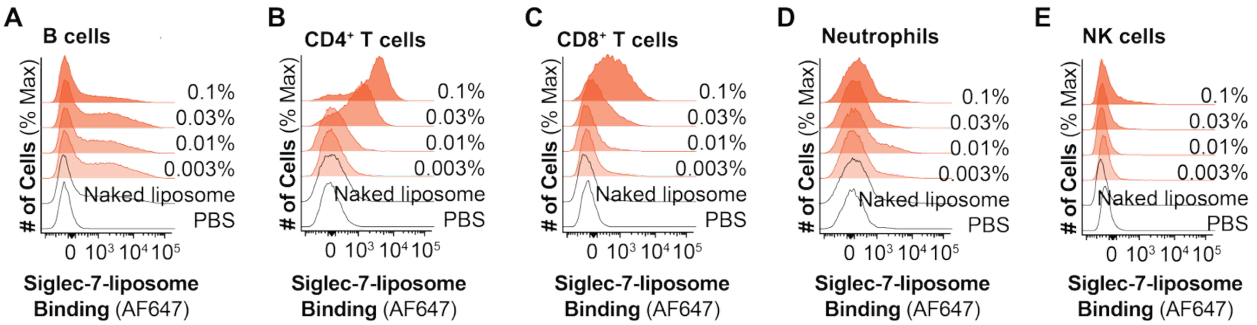


**Fig. S22. *In vivo* binding of Siglec-7-liposomes to mouse splenocytes**. Binding of Naked liposome, 0.003%, 0.01%, 0.03%, and 0.1% of WT Siglec-7-liposomes to (**A**) B cells, (**B**) CD4$^{+}$ T cells, (**C**) CD8$^{+}$ T cells, (**D**) neutrophils, and (**E**) NK cells. Data is presented as flow cytometry histograms.

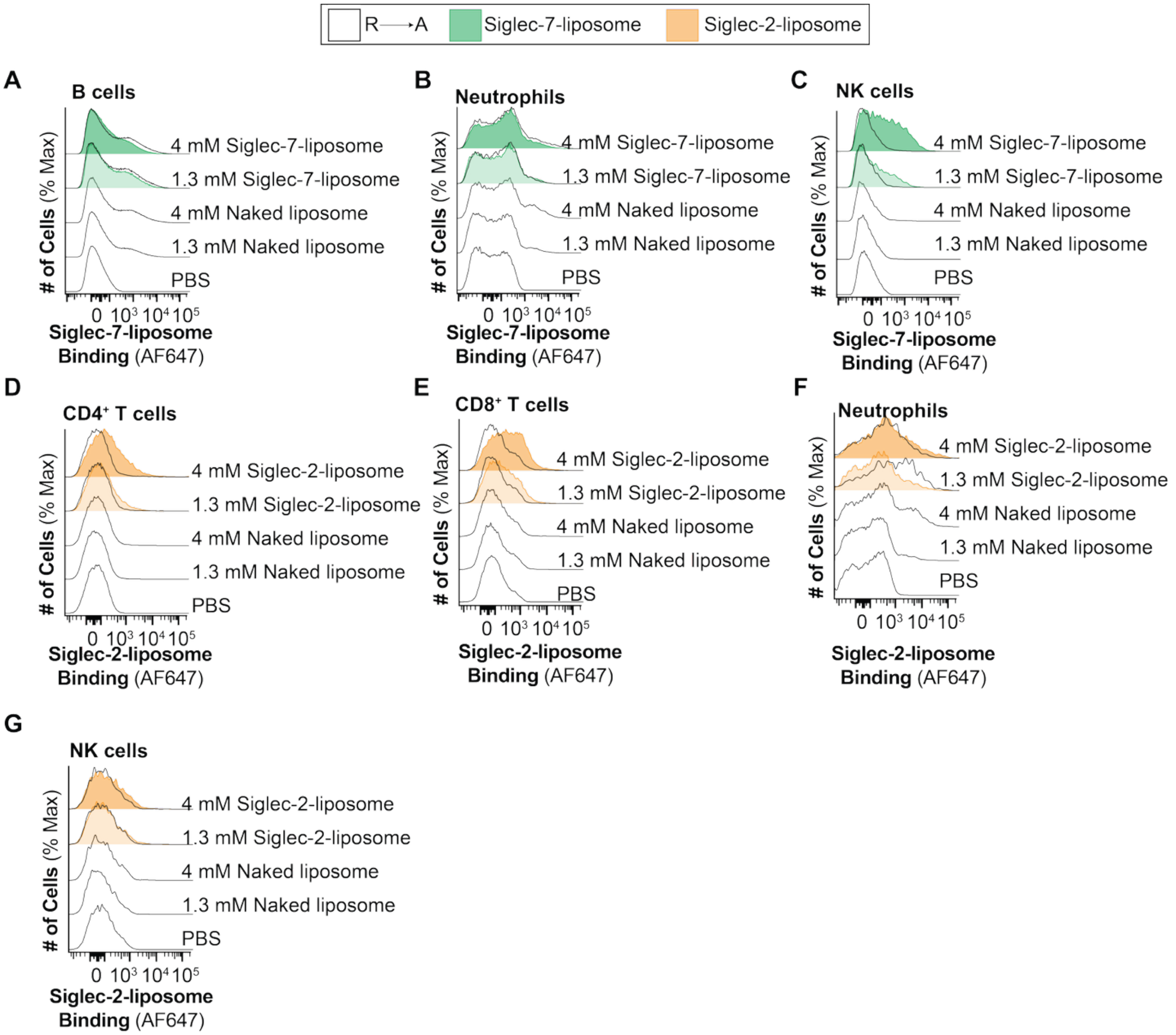


**Fig. S23. *In vivo* binding of Siglec-7-liposomes and Siglec-2-liposomes to mouse splenocytes.** Binding of 1.3 mM and 4mm of Naked liposome, R124A, and WT Siglec-7-liposomes to (**A**) B cells, (**B**) neutrophils, and (**C**) NK cells. Binding of 1.3 mM and 4mm of Naked liposome, R120A and WT Siglec-2-liposomes to (**D**) CD4$^+$ T cells, (**E**) CD8$^+$ T cells, (**F**) neutrophils, and (**G**) NK cells. Data is presented as flow cytometry histograms.

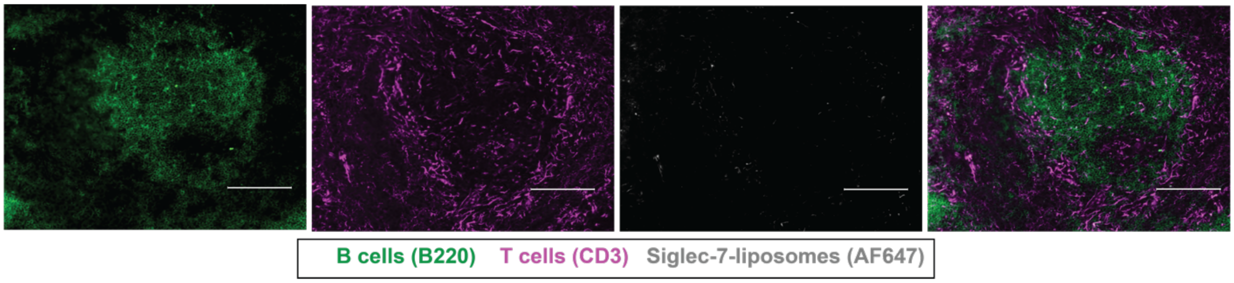


**Fig. S24.** IF microscopy images of mice spleen tissues representing intravenously injection and binding of R124A Siglec-7-liposomes to T cells and B cells. Scale bar: 100 μm.

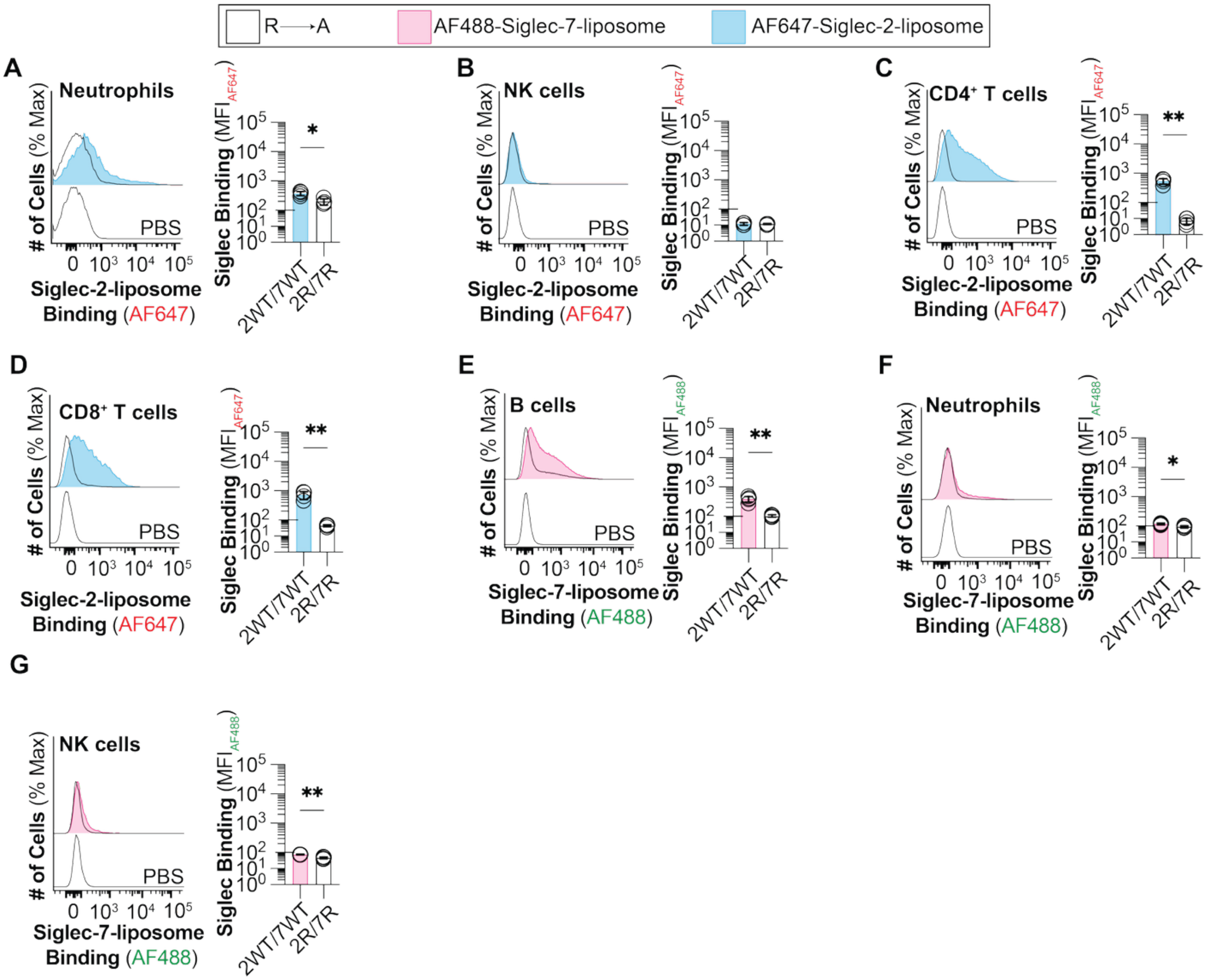


**Fig. S25. *In vivo* binding of multiplexed AF488-Siglec-7-liposomes and AF647-Siglec-2-liposomes to mouse splenocytes.** Binding of multiplexed WT and R120A Siglec-2-liposomes to (**A**) neutrophils, (**B**) NK cells, (**C**) $CD4^+$ T cells, and (**D**) $CD8^+$ T cells. Binding of multiplexed WT and R124A Siglec-7-liposomes to (**E**) B cells, (**F**) neutrophils, and (**G**) NK cells. Data is presented as representative flow cytometry histograms and MFI of Siglec binding. The $p$ value for four biological replicates was calculated using a paired Student's $t$ test. *$0.05>p\geq0.01$, c**$0.01>p\geq0.001$.

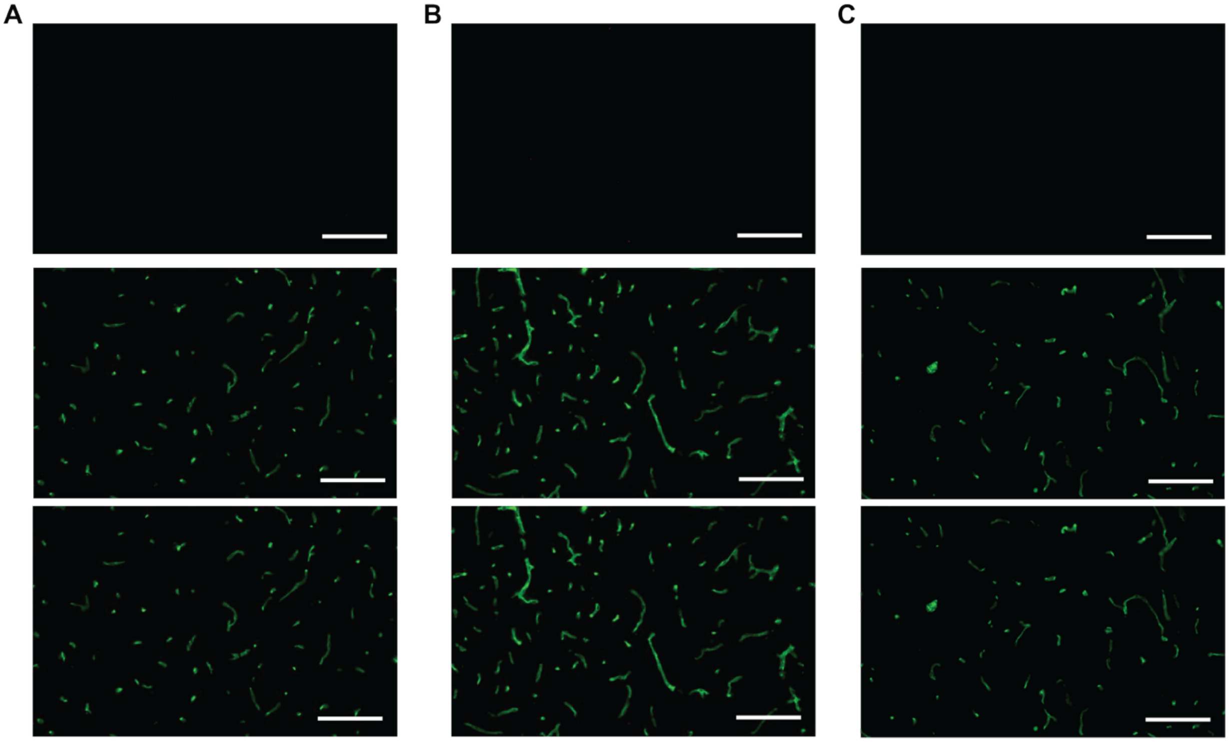


**Fig. S26.** IF microscopy images of mice brain tissues representing intravenously injection and binding of (**A**) PBS, (**B**) naked liposome, and (**C**) R124A Siglec-7-liposomes to endothelial cells in mouse brain. Scale bar: 100 μm.

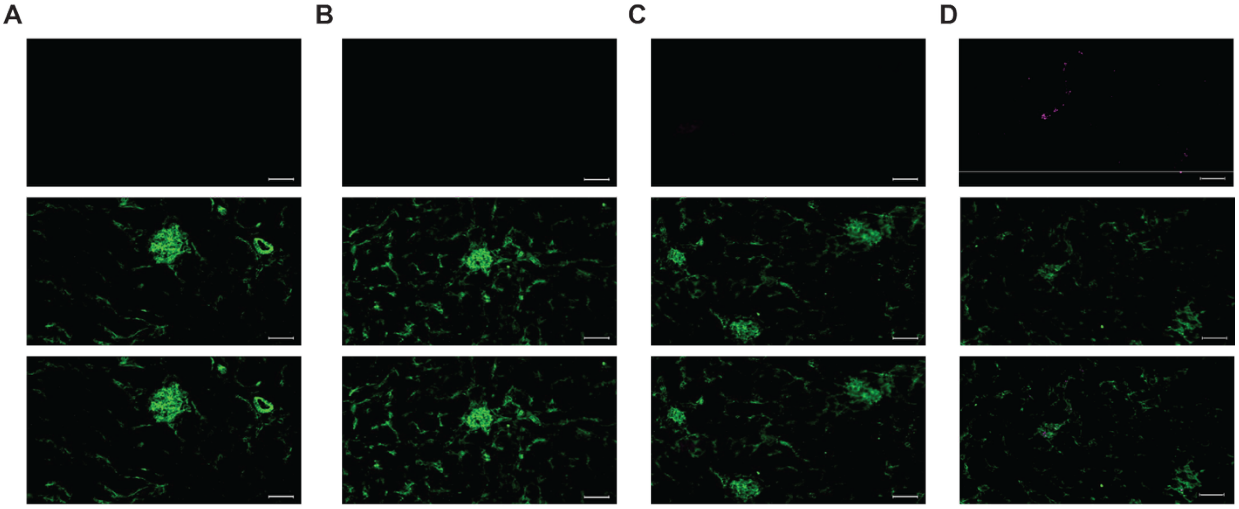


Siglec-7-liposomes (AF647) Endothelial cells (CD31)

**Fig. S27.** IF microscopy images of mouse kidney tissues representing intravenously injection and binding of (**A**) PBS, (**B**) naked liposome, (**C**) R124A Siglec-7-liposomes, and (**D**) WT Siglec-7-liposomes to endothelial cells in mouse kidney tissues. Scale bar: 50 μm.

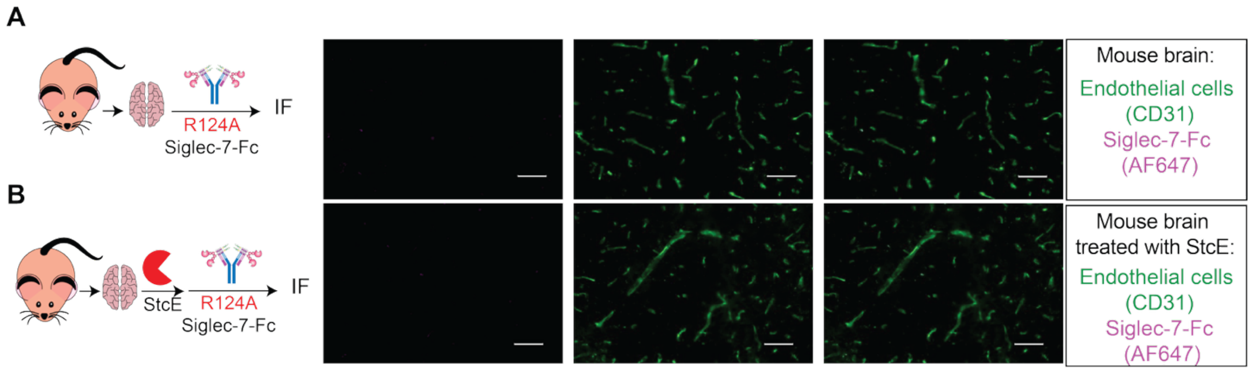


**Fig. S28.** IF microscopy images of *ex vivo* staining of mouse brain tissues representing binding of (**A**) R124A Siglec-7-Fc, (**B**) treatment with StcE mucinase enzyme followed with staining with R124A Siglec-7-Fc and αCD31. Scale bar: 50 μm.

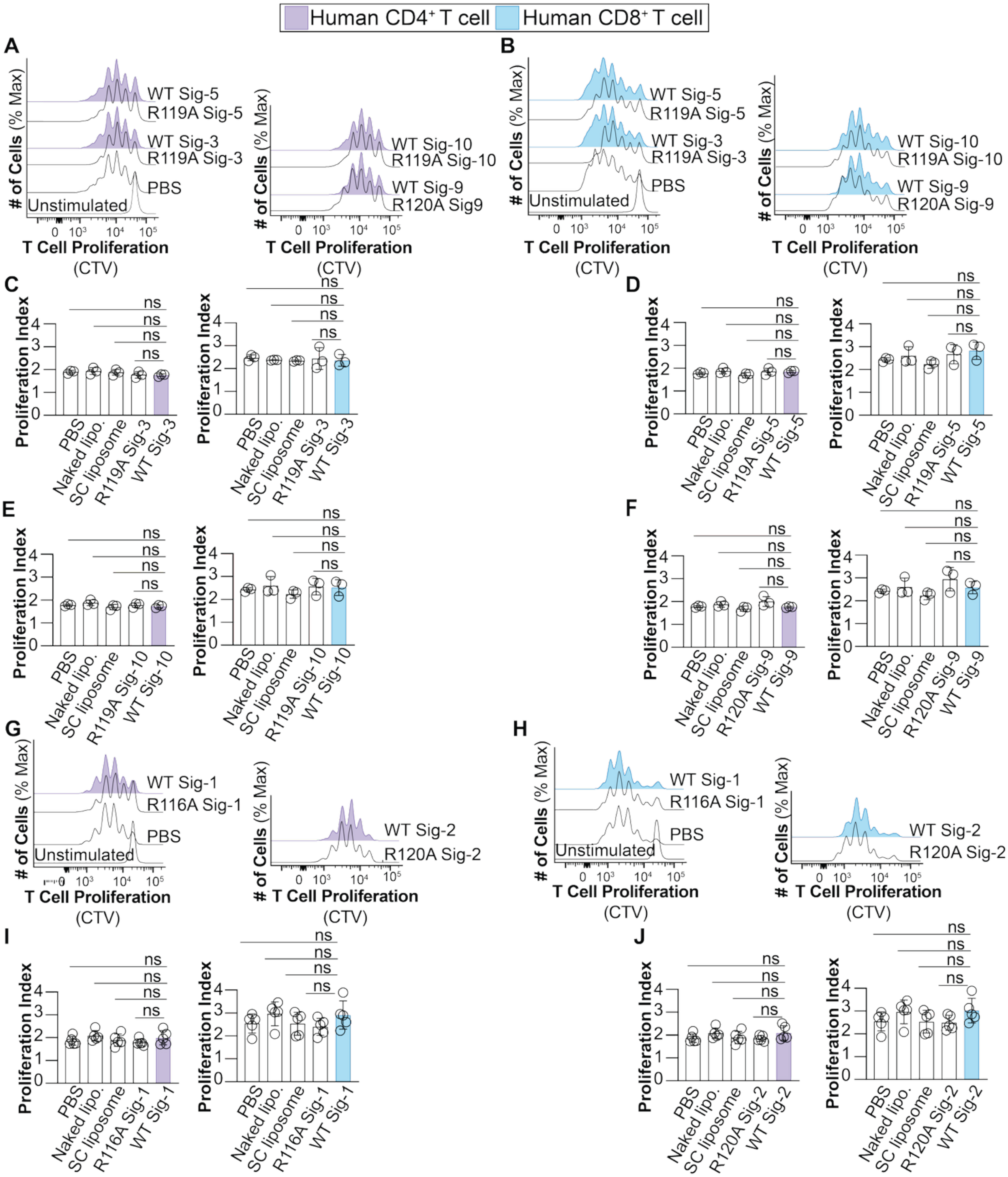

**Fig. S29. Impact of Siglec-1, -2, -3, -5, -9, and -10 -liposomes on human T cell proliferation.** (**A-F**) T cell Proliferation of human CD4$^+$ and CD8$^+$ T cells activated with αCD3/28 beads and co-cultured with WT and R119A Siglec-3-, WT and R119A Siglec-5-, WT and R120A Siglec-9-, WT and R119A Siglec-10-liposomes. (**G-J**) T cell Proliferation of human CD4$^+$ and CD8$^+$ T cells activated with αCD3/28 beads and co-cultured with WT and R116A Siglec-1-, WT and R120A Siglec-2-liposomes. Data is presented as flow cytometry histograms (*upper* panels) and proliferation index (*lower* panels). The *p* value for three (C-F) and five (I,J) biological replicates was calculated using a paired one-way ANOVA Not significant (ns) $p>0.05$.

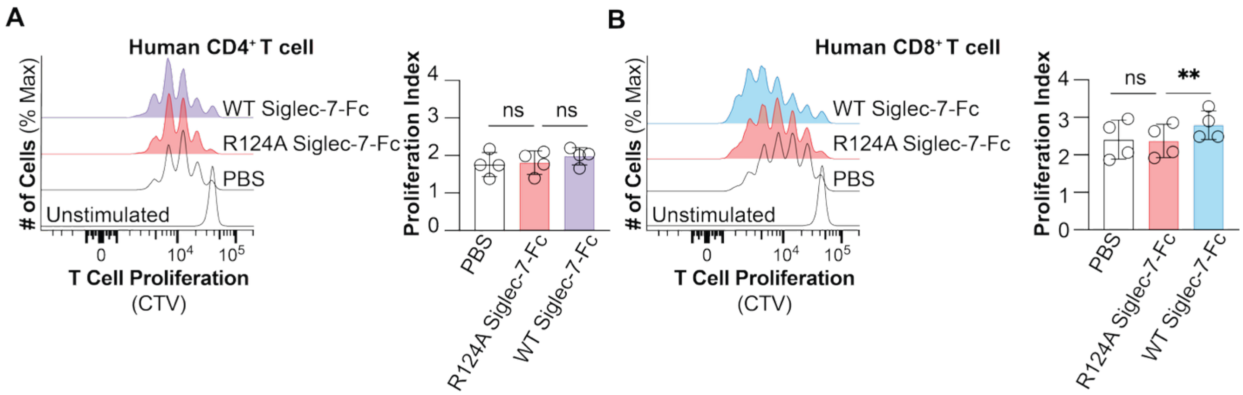


**Fig. S30. Modulation of T cell proliferation by soluble Siglec-7-Fc.** T cell proliferation of (**A**) human CD4$^+$ and (**B**) CD8$^+$ T cells activated with αCD3/28 beads and co-cultured with soluble WT and R124A Siglec-7-Fc. Data is presented as flow cytometry histograms (*left* panel), and proliferation index of T cells (*right* panel). The *p* value for four biological replicates was calculated using a paired one-way ANOVA. Not significant (ns) $p>0.05$, $^{**}0.01>p\geq0.001$.

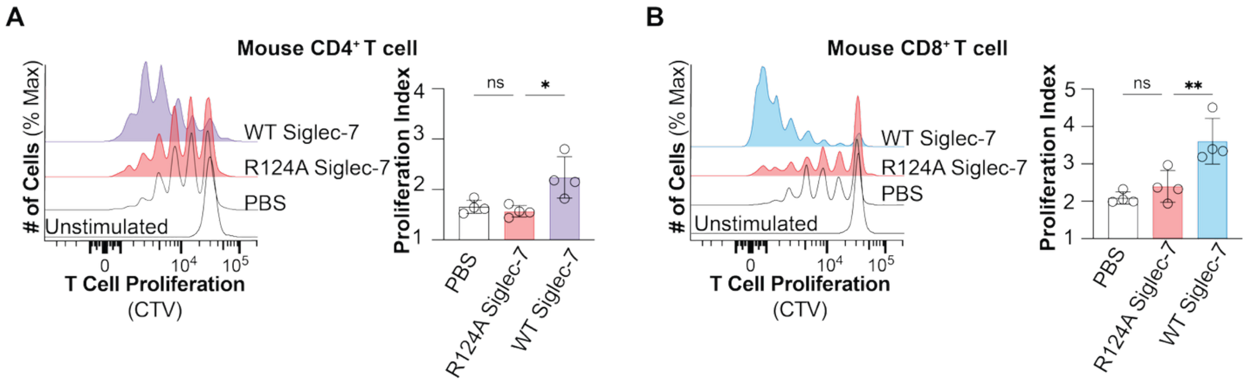


**Fig. S31. Modulation of mouse T cell proliferation by Siglec-7-liposomes after 96 hours.** (**A**) T cell proliferation of mouse $CD4^+$ T cells activated with αCD3/28 beads and co-cultured with WT and R124A Siglec-7-liposomes after 96 hours. (**B**) T cell proliferation of mouse $CD8^+$ T cells activated with αCD3/28 beads and co-cultured with WT and R124A Siglec-7-liposomes after 96 hours. Data is presented as flow cytometry histograms (*left* panel), and T cell proliferation index (*right* panel). The *p* value for four biological replicates was calculated using a paired one-way ANOVA. Not significant (ns) $p>0.05$, $*0.05>p\geq0.01$, $**0.01>p\geq0.001$.

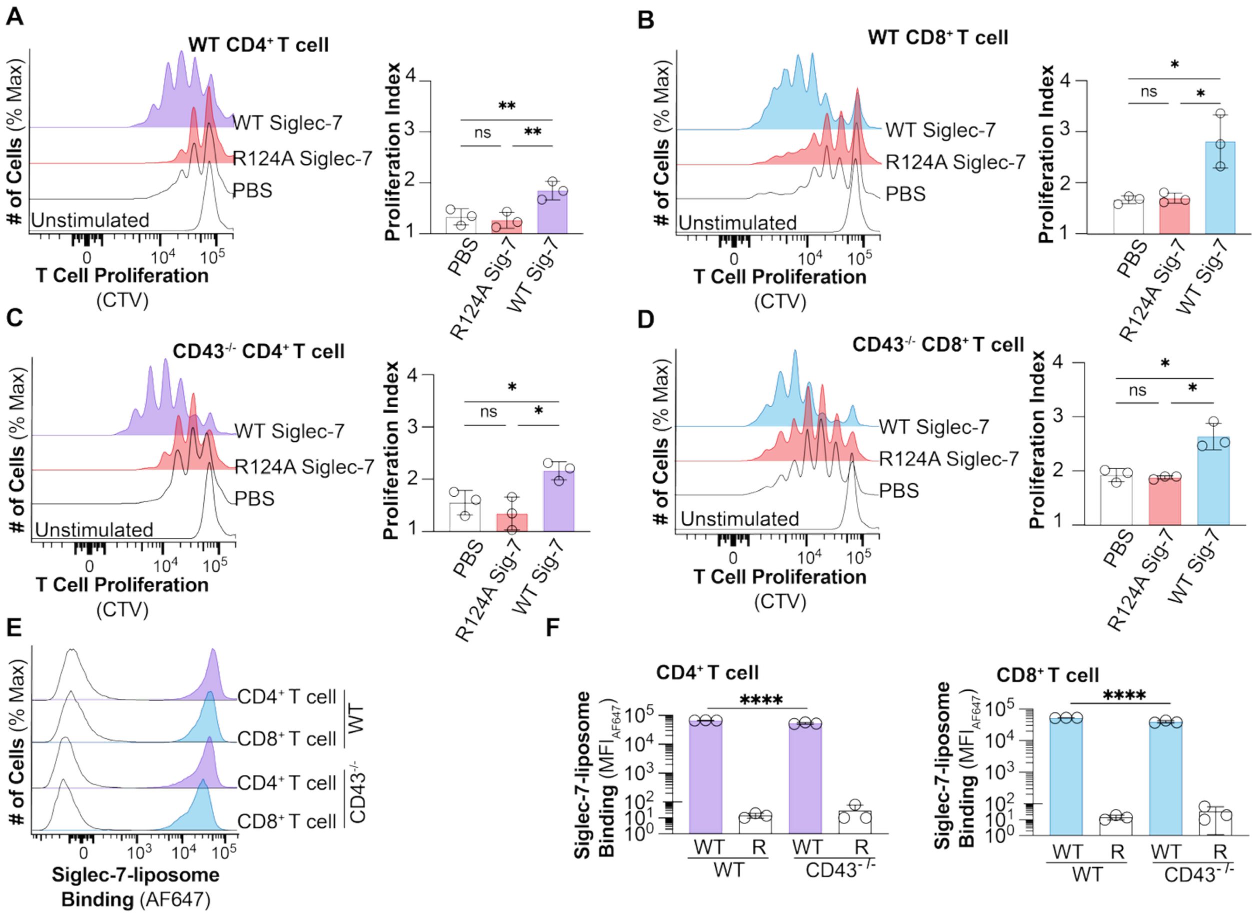


**Fig. S32. Impact of Siglec-7-liposomes on CD43$^{-/-}$ T cell proliferation.** T cell proliferation of (**A**) WT CD4$^{+}$ T cells, (**B**) WT CD8$^{+}$ T cells, (**C**) CD43$^{-/-}$ CD4$^{+}$ T cells, and (**D**) CD43$^{-/-}$ CD8$^{+}$ T cells activated with αCD3/28 beads and co-cultured with PBS, R124A, and WT Siglec-7-liposomes. Data is presented as flow cytometry histograms (*left* panels) and T cell proliferation index (*right* panels). Binding of WT and R124A Siglec-7-liposomes to WT and CD43$^{-/-}$ T cells. Data is presented as (**E**) flow cytometry histograms and (**F**) MFI of Siglec-7 binding. The *p* value for three biological replicates was calculated using a paired one-way ANOVA in panels a-d. The *p* value for three technical replicates was calculated using an unpaired one-way ANOVA in panels f. Not significant (ns) $p>0.05$, $*0.05>p\geq0.01$, $**0.01>p\geq0.001$, $****p<0.0001$.

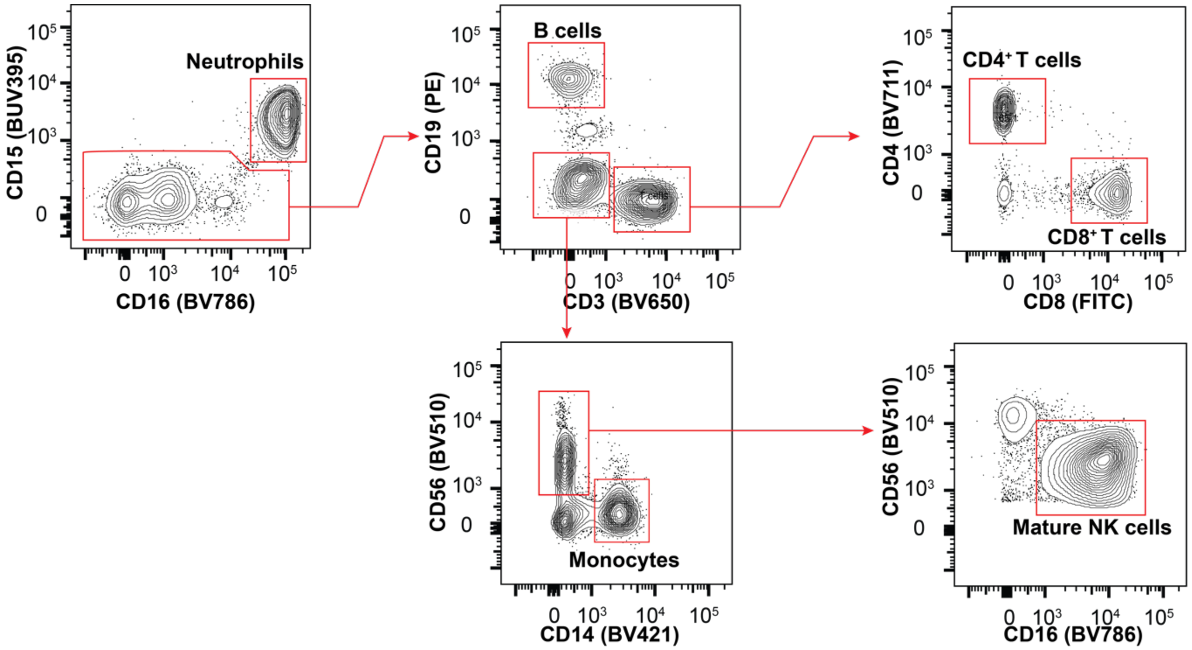


**Fig. S33. Flow cytometry gating strategy to isolate primary immune cells from human blood.** Neutrophils ($CD15^+CD16^+$), B cells ($CD19^+$), $CD4^+$ T cells ($CD3^+CD4^+$) and $CD8^+$ T cells ($CD3^+CD8^+$), monocytes ($CD14^+$), and mature natural killer cells ($CD56^+CD16^+$).

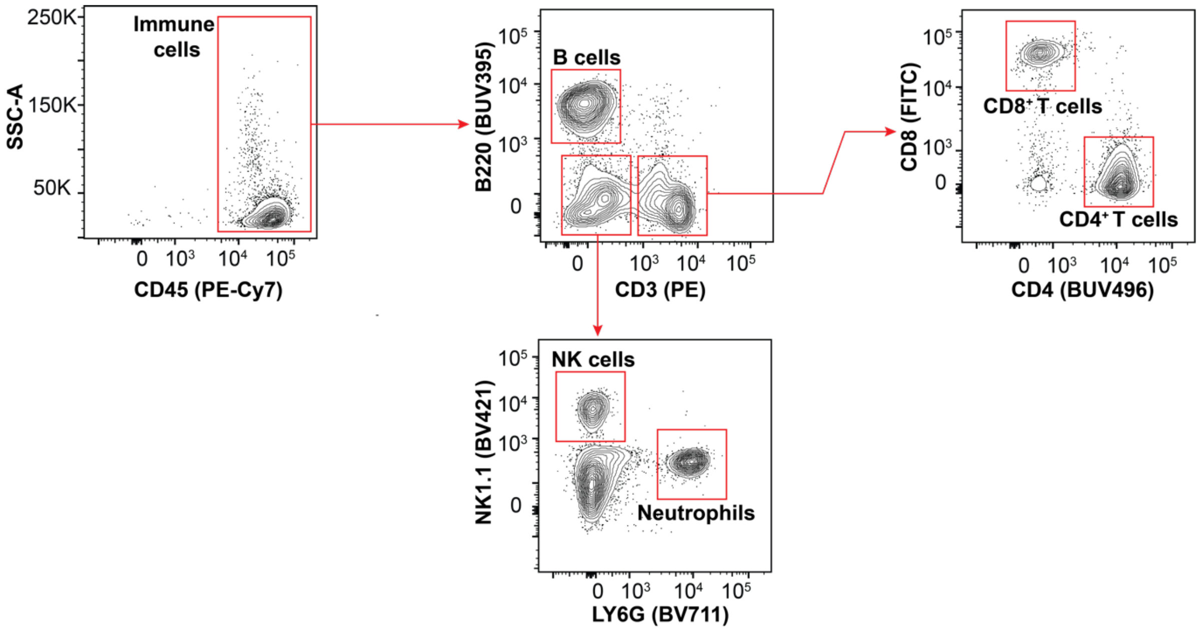


**Fig. S34. Flow Cytometry gating strategy for primary immune cells from mouse splenocytes.** Leukocytes ($CD45^+$) are initially selected, followed by gating for B cells ($B220^+$), $CD4^+$ T cells ($CD3^+CD4^+$) and $CD8^+$ T cells ($CD3^+CD8^+$), natural killer cells ($NK1.1^+$), and neutrophils ($LY6G^+$).

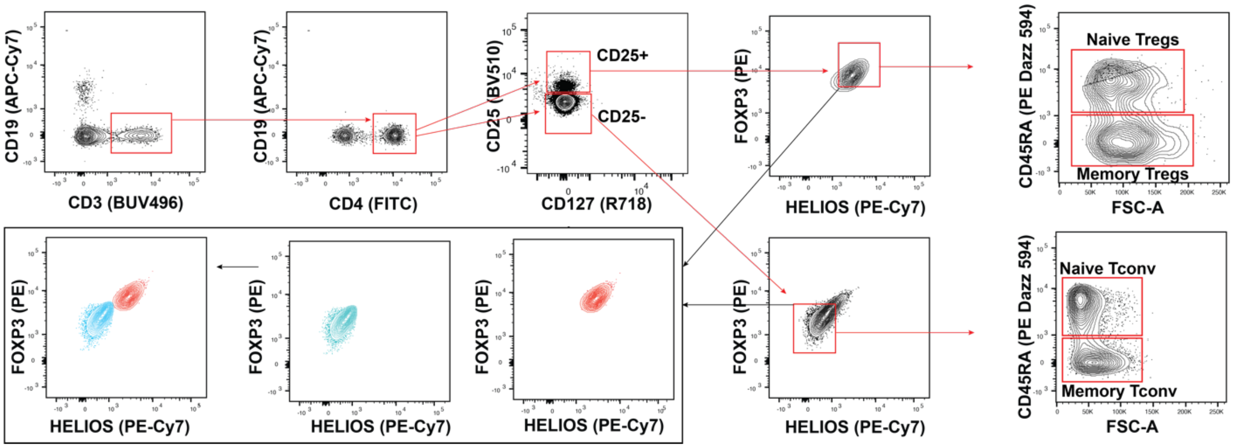


**Fig. S35. Flow cytometry gating strategy for regulatory T cells (Tregs) and conventional T cells (Tconv).** T cells ($CD3^+$) are first selected away from B cells ($CD19^+$). $CD4^+$ T cells are selected and gated on the $CD25^+$ and $CD25^-$ cells. A combination of FOXP3 and HELIOS are used to distinguish Treg ($FOXP3^+HELIOS^+$) and Tconv ($FOXP3^-$ $HELIOS^-$) cells followed by gating for naïve Treg and Tconv cells ($CD45RA^+$) and memory Treg and Tconv cells ($CD45RA^-$).

**Table S1. PCR primers for cloning Version 1 (Siglec-ST-Fc) and Version 2 (Siglec-Fc-ST) and development of CMAS$^{-/-}$ Siglecs.**

| | |
|---|---|
| **P1** (ST- Fc) | GCCTACAAACGCTATAAGGAGAACCTGTACTTCCAGG |
| **P2** (*XmaI*- Fc) | AGCAGCCCCGGGTCACTTCTCGAACTGGGGGTGGGA |
| **P3** (*AgeI*- ST) | AGCAGCACCGGTCGTGGGGTTCCACACATAGTAATGGTGGAT GCCTACAAACGCTATAAG |
| **P4** (*AgeI*- Fc) | AGCAGCACCGGTGAGAACCTGTACTTCCAGGGGGAC |
| **P5** (*XmaI*- ST) | AGCAGCCCCGGGTCACTTATAGCGTTTGTAGGCATCCACCATTACTA TGTGTGGAACCCCACGCTTCTCGAACTGGGGGTGGGA |
| **P6** | CCTAGTCAGGTCTGTTGTCACCAGGC |
| **P7** | TTCCTCGAATGGGGAGATGTGCG |

**Table S2. Human antibodies used in this study.**

| Antibody | Supplier | Cat. No. | Label | Clone | Isotype | Dilution |
|---|---|---|---|---|---|---|
| CD19 | BioLegend | 363004 | PE | SJ25C1 | Mouse IgG1, κ | 1:150 |
| CD19 | BioLegend | 363010 | APC-CY7 | SJ25C1 | Mouse IgG1, κ | 1:150 |
| CD3 | BioLegend | 317324 | BV650 | OKT3 | Mouse IgG2a, κ | 1:150 |
| CD3 | BD Bioscience | 612941 | BUV496 | UCHT1 | Mouse IgG1, κ | 1:150 |
| HLA DR | BioLegend | 307618 | APC Cy7 | L243 | Mouse IgG2a, κ | 1:150 |
| CD15 | BD Bioscience | 563872 | BUV395 | HI98 | Mouse IgM, κ | 1:150 |
| CD56 | BioLegend | 362533 | BV510 | 5.1 H11 | Mouse IgG1, κ | 1:150 |
| CD4 | BioLegend | 300558 | BV711 | RPA-T4 | Mouse IgG1, κ | 1:150 |
| CD4 | BioLegend | 980802 | FITC | SK3 | Mouse IgG1, κ | 1:150 |
| CD8a | BioLegend | 300914 | PE-Cy7 | HIT8a | Mouse IgG1, κ | 1:150 |
| CD8a | BioLegend | 300905 | FITC | HIT8a | Mouse IgG1, κ | 1:150 |
| CD14 | BioLegend | 301829 | BV421 | M5E2 | Mouse IgG2a, κ | 1:150 |
| CD16 | BD Bioscience | 563689 | BV786 | 3G8 | Mouse IgG1, κ | 1:150 |
| CD45RA | BioLegend | 304145 | PE-Daz-594 | HI100 | Mouse IgG2b, κ | 1:150 |
| FOXP3 | BioLegend | 364704 | PE | QA18A03 | Mouse IgG1, κ | 1:50 |
| HELIOS | BioLegend | 137235 | PE/CY7 | 22F6 | American Hamester IgG | 1:50 |
| CD25 | BioLegend | 302640 | BV510 | BC96 | Mouse IgG1, κ | 1:150 |
| FcX | BioLegend | 422302 | | | | 1:100 |

**Table S3. Mouse antibodies used in this study.**

| Antibody | Supplier | Cat. No. | Label | Clone | Isotype | Dilution |
|---|---|---|---|---|---|---|
| CD45 | BioLegend | 103114 | PE-Cy7 | 30-F11 | Rat IgG2b, κ | 1:200 |
| B220 | BD Bioscience | 563793 | BUV395 | RA3-6B2 | Rat IgG2a, κ | 1:200 |
| CD3 | BioLegend | 100206 | PE | 17A2 | Rat IgG2b, κ | 1:200 |
| CD4 | BD Bioscience | 612952 | BUV496 | GK1.5 | Rat LEW, also known as Lewis IgG2b, κ | 1:200 |
| CD4 | BioLegend | 100447 | BV711 | GK1.5 | Rat IgG2b, κ | 1:200 |
| CD8a | BioLegend | 100723 | AF488 | 53-6.7 | Rat IgG2a, κ | 1:200 |
| F4/80 | BD Bioscience | 743282 | BV650 | T45-2342 | Rat WI, also known as Wistar (outbred) IgG2a, κ | 1:200 |
| NK1.1 | BioLegend | 108741 | BV421 | PK136 | Mouse IgG2a, κ | 1:200 |
| Ly6G | BD Bioscience | 563979 | BV711 | 1A8 | Rat LEW, also known as Lewis IgG2a, κ | 1:200 |
| Ly6C | BioLegend | 128026 | APC-Cy7 | HK1.4 | Rat IgG2c, κ | 1:200 |
| CD11b | BioLegend | 101245 | BV510 | M1/70 | Rat IgG2b, κ | 1:200 |
| CD11c | BioLegend | 117336 | BV786 | N418 | American hamster IgG | 1:200 |
| CD43 | BioLegend | 143205 | PE | S11 | Rat IgG2b, κ | 1:200 |
| Biotin-B220 | BioLegend | 103204 | | RA3-6B2 | Rat IgG2a, κ | 1μg/mL |
| Biotin-CD11c | BioLegend | 117304 | | N418 | American hamster IgG | 1μg/mL |

| | | | | | | |
|---|---|---|---|---|---|---|
| Biotin-CD11b | BioLegend | 101204 | | M1/70 | Rat IgG2b, κ | 1μg/mL |
| Biotin-Ter119 | BioLegend | 116204 | | TER-119 | Rat IgG2b, κ | 1μg/mL |
| Biotin-Gr-1 | BioLegend | 108403 | | RB6-8C5 | Rat IgG2b, κ | 1μg/mL |
| Biotin-TCRγ/β | BioLegend | 118103 | | GL3 | American hamster IgG | 1μg/mL |
| Biotin-CD19 | BioLegend | 115504 | | 6D5 | Rat IgG2a, κ | 1μg/mL |
| Biotin-CD49b | BioLegend | 108904 | | DX5 | Rat IgM, κ | 1μg/mL |
